# BT-11 repurposing potential for Alzheimer’s disease and insights into its mode of actions

**DOI:** 10.1101/2024.10.29.620882

**Authors:** Emily M. Birnbaum, Lei Xie, Peter Serrano, Patricia Rockwell, Maria E. Figueiredo-Pereira

## Abstract

Neuroinflammation is a key pathological hallmark of Alzheimer’s disease (AD). Investigational and FDA approved drugs targeting inflammation already exist, thus drug repurposing for AD is a suitable approach. BT-11 is an investigational drug that reduces inflammation in the gut and improves cognitive function. BT-11 is orally active and binds to lanthionine synthetase C-like 2 (LANCL2), a glutathione-s-transferase, thus potentially reducing oxidative stress. We investigated the effects of BT-11 long-term treatment on the TgF344-AD rat model. BT-11 reduced hippocampal-dependent spatial memory deficits, Aβ plaque load and neuronal loss in males, and mitigated microglia numbers in females. BT-11 treatment led to hippocampal transcriptomic changes in signaling receptor, including G-protein coupled receptor pathways. We detected LANCL2 in hippocampal nuclear and cytoplasmic fractions with potential different post-translational modifications, suggesting distinct functions based on its subcellular localization. LANCL2 was present in oligodendrocytes, showing a role in oligodendrocyte function. To our knowledge, these last two findings have not been reported. Overall, our data suggest that targeting LANCL2 with BT-11 improves cognition and reduces AD-like pathology by potentially modulating G-protein signaling and oligodendrocyte function. Our studies contribute to the field of novel immunomodulatory AD therapeutics, and merit further research on the role of LANCL2 in this disease.

## Introduction

Chronic neuroinflammation is a key factor in AD progression.^1^ Focusing on regulators of neuroinflammation for treatment of AD is an important avenue of current research. LANCL2 is a novel regulator of inflammation, and BT-11 is a LANCL2 activator. BT-11 was selected based on a computational drug repurposing approach for AD candidates, and on its immunomodulatory effects knowing that inflammation plays a critical role in AD. BT-11 is currently in clinical trials for Irritable Bowel Disease (IBD).^2,3^ BT-11 [N,N-Bis(benzimadzolylpicolinoyl)piperazine] is a synthetic organic compound derived from the dried root extract of the *Polygala tenuifolia* wildenow plant.^4,5^ BT-11 was originally developed by designing analogs of abscisic acid (ABA), the endogenous ligand for LANCL2. Molecular docking and binding assays to LANCL2 revealed that BT-11 was the top lead compound.^4^

Multiple studies showed beneficial effects of BT-11 on cognitive deficits and in the context of aging but none, to our knowledge, in the context of AD. One study used an amnesia model in both male and female pregnant Sprague-Dawley rats and found that a 10mg/kg intra-peritoneal (i.p.) injection of BT-11 recovered performance on passive avoidance and water maze tasks. ^5^ Another group looked at the effect of BT-11 on elderly humans.^6^ This randomized, double-blind, placebo-controlled comparison study used the Consortium to Establish a Registry for Alzheimer’s Disease (CERAD) and the Mini-Mental State Examination (MMSE). Total CERAD scores were significantly improved in the BT-11 treatment group as compared to placebo, demonstrating that BT-11 enhances cognitive functions in aged adults.^6^ These studies merit further investigation of BT-11 in cognition as it occurs in AD. To our knowledge, our studies address for the first time the impact of BT-11 on AD-relevant spatial memory deficits and hippocampal pathology.

At the diagnostic stage, AD patients already show significant cognitive deficits, and substantial accumulation of brain pathology. This makes developing and investigating therapeutics very difficult and suggests a need for examining the neuropathology of AD in transgenic animal models.^7^ We used the TgF344-AD rat model that develops AD pathology including amyloid plaques, tauopathy, gliosis, neuronal loss, and cognitive deficits in a progressive age-dependent manner, an aspect of AD that is difficult to reproduce in animal models.^8,9^ TgF344-AD rats express the human Swedish amyloid precursor protein (APPsw) and Δ exon 9 presenilin-1 (PS1-ΔE9) genes, driven by the prion promoter.^8^ BT-11 was administered orally and daily (8 mg/kg bw/day) for a period of 6 months to transgenic and wild type male and female rats, and its effects were assessed on hippocampal-dependent spatial memory and AD-pathology.

In summary, our studies established that: (1) long-term oral treatment of TgF344-AD rats with BT- 11 improved spatial memory and reduced Aβ plaque load and neuronal loss in males, and microgliosis in females; (2) hippocampal LANCL2 is localized in the nuclear and cytoplasmic fractions with possible localization-specific post-translational modifications, such as myristoylation, phosphorylation and/or ubiquitination; (3) LANCL2 is specifically located in hippocampal oligodendrocytes, and not in neurons, astrocytes or microglia. Together, our results strongly support the potential of BT-11 to treat AD, and the need to continue investigating the role of the LANCL2 pathway in AD.

## Results

### BT-11 treatment improved spatial learning in 11-month male TgF344-AD rats

Our studies included four groups of rats for each sex: WTNT – Wild-type not treated; TGNT – transgenic not treated; WTTR – Wild-type RG2833-treated; TGTR – transgenic RG2833-treated. After six consecutive months of drug treatment, all rats were assessed for cognitive behavior at 11 months of age, as previously described by our group.^10^ For this study, three different cohorts of rats were included, and males and females were analyzed separately. Across these cohorts there were WTNT (n=14), TGNT (n=17), WTTR (n=13), and TGTR (n=13) males, as well as WTNT (n=28), TGNT (n=16), WTTR (n=12), and TGTR (n=11) females. To show that any differences in behavior were due to cognitive abilities and not movement or motivation, we analyzed the total path length of all rats in their habituation trial. During this trial, all rats should be freely exploring the arena. There were no significant differences in path length across all male groups and across all female groups (Supplemental Table 9, Supplemental Figure 1).

All male rats were able to learn across the six trials as showed by a significant training effect (Fig 1b-e, p<0.0001). There was a significant deficit in spatial learning in the TGNT male rats as compared to the WTNT male rats in the measure of percent time in the target zone with significant post-hoc effects in the first two trials (Fig 1b, genotype effect p<0.01). This deficit was mitigated by BT-11 treatment, with TGTR male rats performing significantly better than TGNT male rats (Fig 1d, drug effect p<0.05). There were no significant differences between WTTR and TGTR (Fig. 1c) or WTNT and WTTR males (Fig 1e). Representative traces from trial three in a WTNT and TGNT male rats are shown in Fig. 1f.

**Fig. 1.**
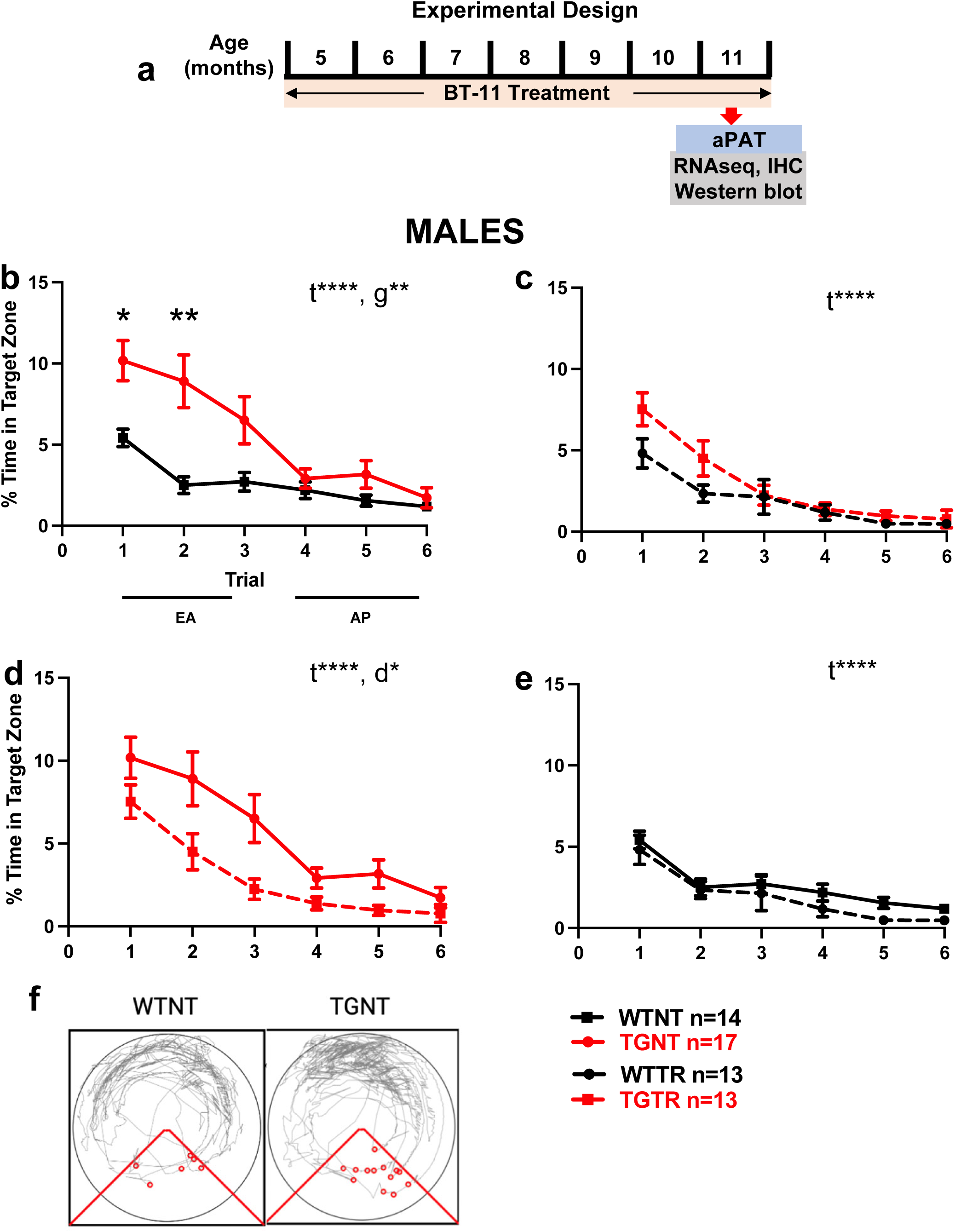
BT-11 improved spatial learning in TgF344-AD males. **a**, Experimental design. All rats were exposed to the aPAT at 11 months of age. Rats were tasked with avoiding the shock zone over a course of six trials. The measure used to analyze this data was percent (%) time in the target zone during a 600 second trial. Following behavioral analysis, the rats were sacrificed (details under methods). **b,** a significant difference was seen during the first three trials (early acquisition) in which the WTNT rats outperform the TGNT rats. **c,** there was no genotype difference between WTTR and TGTR rats. **d,** TGTR rats outperformed the TGNT rats. **e,** there was no treatment difference between WTTR and WTNT rats. **f,** Representative track tracings of a WTNT and a TGNT male performance for trial 3. Two-way ANOVAs with Tukey’s post hoc test with GraphPad Prism 9 software were used, \**P*<0.05, ** *P* <0.01, **** *P* <0.0001. EA - Early Acquisition; AP – Asymptotic Performance; WTNT – Wild-type not treated; TGNT – transgenic not treated; WTTR – Wild-type BT-11-treated; TGTR – transgenic BT-11-treated; t=training effect, g=genotype effect, d=drug effect; aPAT, active place avoidance test; RNAseq, RNA sequencing; IHC, immunohistochemistry.

All female rats were able to learn across the six trials as shown by significant training effects, WTNT vs. TGNT, TGNT vs. TGTR, WTTR vs. TGTR (Fig. 2a to c, p<0.0001), and WTTR vs. WTNT (Fig. 2d, p<0.001). However, there were no significant genotype or behavior differences across all female groups (Fig 2, Supplemental Table 2). Representative traces from trial three in a WTNT and TGNT female rats are shown in Fig. 2e.

**Fig. 2.**
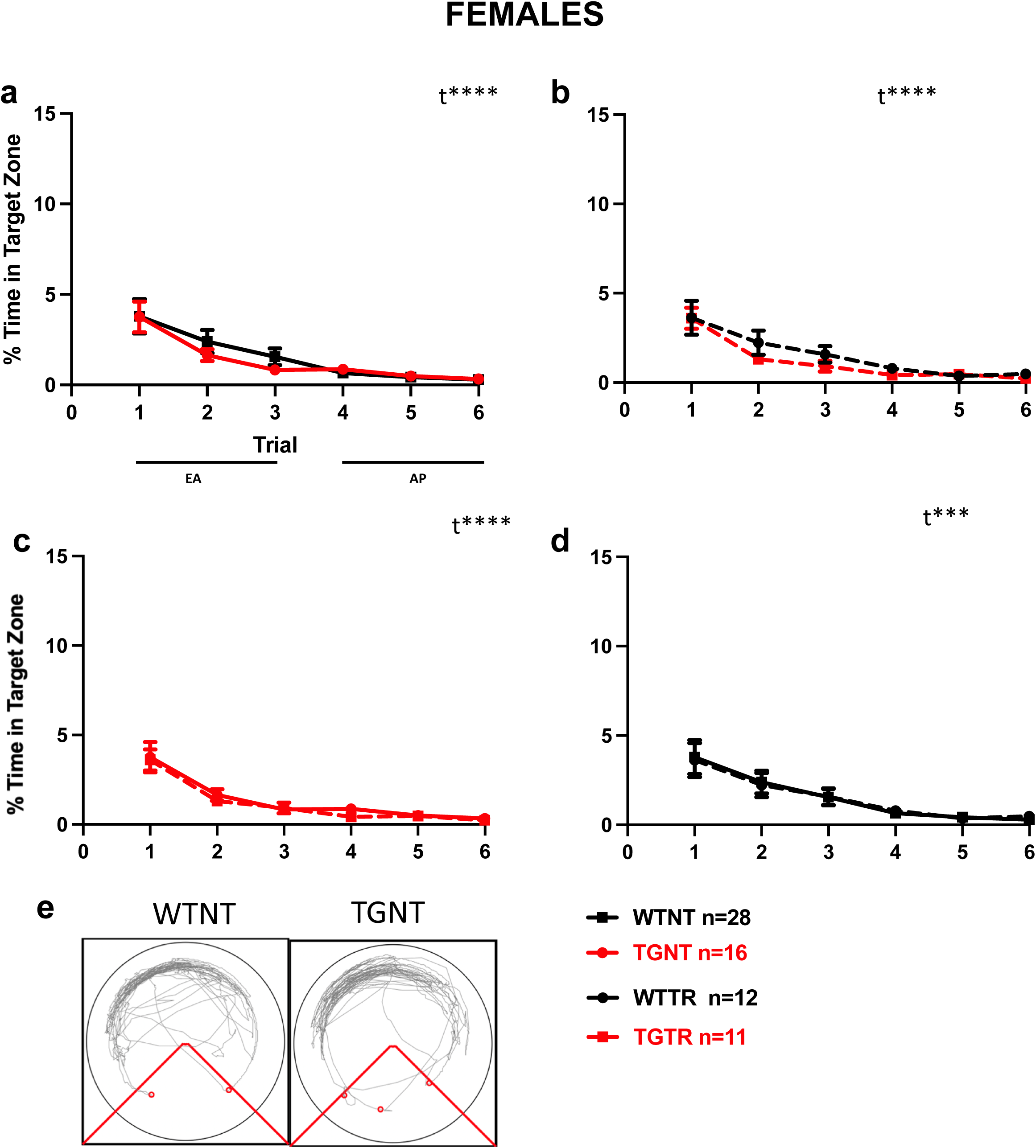
BT-11 did not improve spatial learning in TgF344-AD females. All females showed learning across six trials with a significant trial effect. The measure used to analyze this data was percent (%) time in the target zone during a 600 second trial. There were no significant genotype differences between **a**, WTNT and TGNT, **b**, WTTR and TGTR, nor treatment differences between **c**, TGNT and TGTR, **d**, WTNT and WTTR. **e,** Representative track tracings of a WTNT and a TGNT female performance for trial 3. Two-way ANOVA with Tukey’s post hoc test with GraphPad Prism 9 software were used, **** *P* <0.0001. EA - Early Acquisition; AP – Asymptotic Performance; WTNT – Wild-type not treated; TGNT – transgenic not treated; WTTR – Wild-type BT-11-treated; TGTR – transgenic BT-11-treated; t= training effect.

### BT-11 treatment reduced Aβ plaque load in 11-month old male TgF344-AD rats

We assessed Aβ plaque burden in TGNT and TGTR rats by IHC analyses (mouse monoclonal antibody 4G8) of the dorsal hippocampus as well as across each of its four subregions: CA1, CA3, DG and SB (Fig 3). In each group, 8 or 9 rats were analyzed. Earlier studies reported that there are no Aβ plaques in WT rats of this AD model ^15^. Images of dorsal hippocampi from a TGNT and a TGTR male are shown in Fig 3a and b, respectively. Aβ plaque load was quantified as % area, which is the % area of the dorsal hippocampus that has Aβ signal. TgF344-AD male**s** treated with BT-11 (n=8) showed reduced plaque load as compared to TGNT males (n=8) in the dorsal hippocampus (Fig 3c, p<0.05, t=1.981). There were no significant differences for each of the hippocampal subregions: CA1 (p=0.535, t=0.637), CA3 (p=0.7140, t=0.375), DG (p=0.289, t=0.570), SB (p=0.734, t=0.347). Females were also assessed for Aβ plaque load and there was no significant difference between TGNT (n=9) and TGTR (n=8) rats in the dorsal hippocampus (Supplemental Table 3, p=0.201, t=0.863), or in each of its subregions: CA1 (p=0.736, t=0.343), CA3 (p=0.781, t=0.283), DG (p=0.991, t=0.012), SB (p=0.178, t=1.413).

**Fig. 3.**
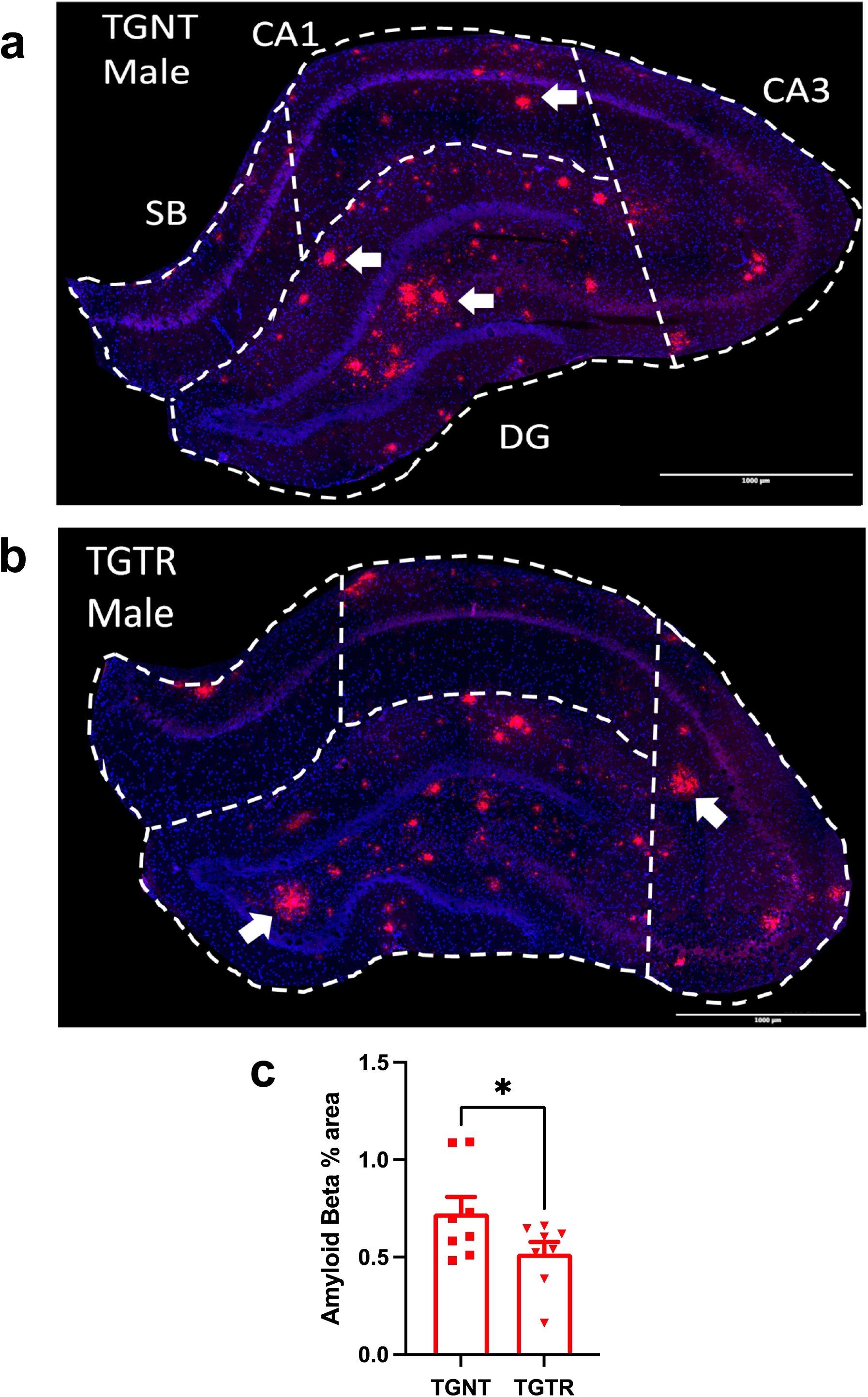
BT-11 treatment reduced Aβ plaque load in 11-month old male TgF344-AD rats. Immunohistochemical analyses for Aβ plaque load (*red*, 4G8 antibody) and DAPI (*blue*) in **a** TGNT and **b** TGTR males. White arrows point to Aβ plaques. Scale bar = 1000µm. **c,** Plaque load was reduced in the hippocampus of TGTR (n=8) compared to TGNT (n=8) males. BT-11 treatment reduced percent (%) area of Aβ signal in the dorsal hippocampus of TGTR male rats. Unpaired one-tail *t*-tests with Welch’s corrections were used for quantification, \**P*<0.05. For simplicity, error bars are shown in one direction only. TGNT – transgenic not treated; TGTR – transgenic BT-11 treated. CA = cornu ammonis, DG = dentate gyrus, SB = subiculum.

### BT-11 treatment reduced total and ramified microglia levels in female TgF344-AD rats

The three morphological states of microglia (Fig. 4a) were quantified in WTNT (males n=8; females n=7), TGNT (males n=8; females n=9), WTTR (males n=8; females n=6), and TGTR (males n=7; females n=8) rats by IHC with the rabbit antibody Iba1. Microglia levels were quantified as % area, representing the % of the total area with positive signal. Levels of all and each subtype of microglia were analyzed in dorsal hippocampi as well as across their four subregions: CA1, CA3, DG and SB. Representative IHC images for males are shown in Fig. 4b-g. Statistical analyses graphs are shown in Fig 4h-i for males and Fig. 4j-m for females.

**Fig. 4.**
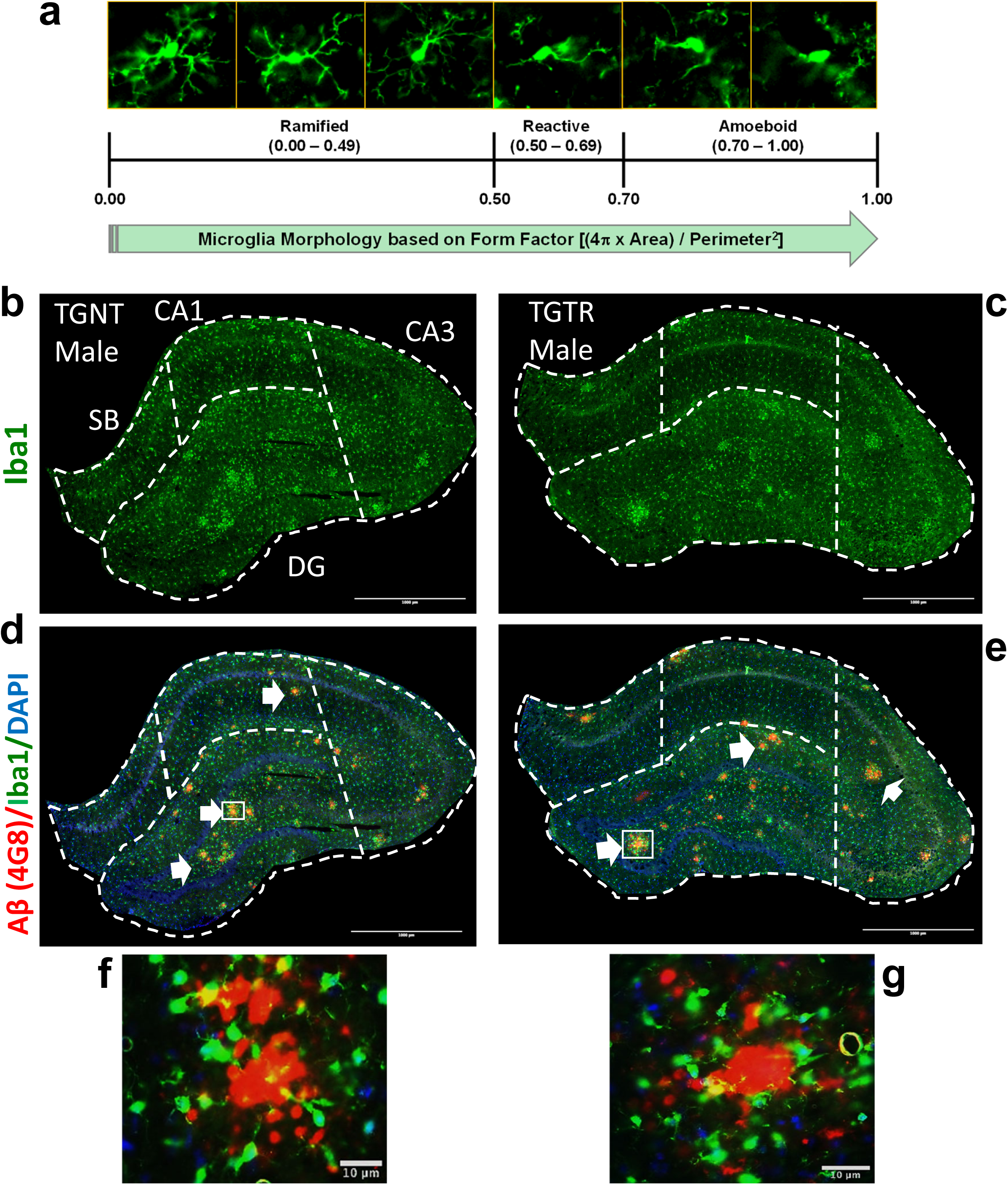
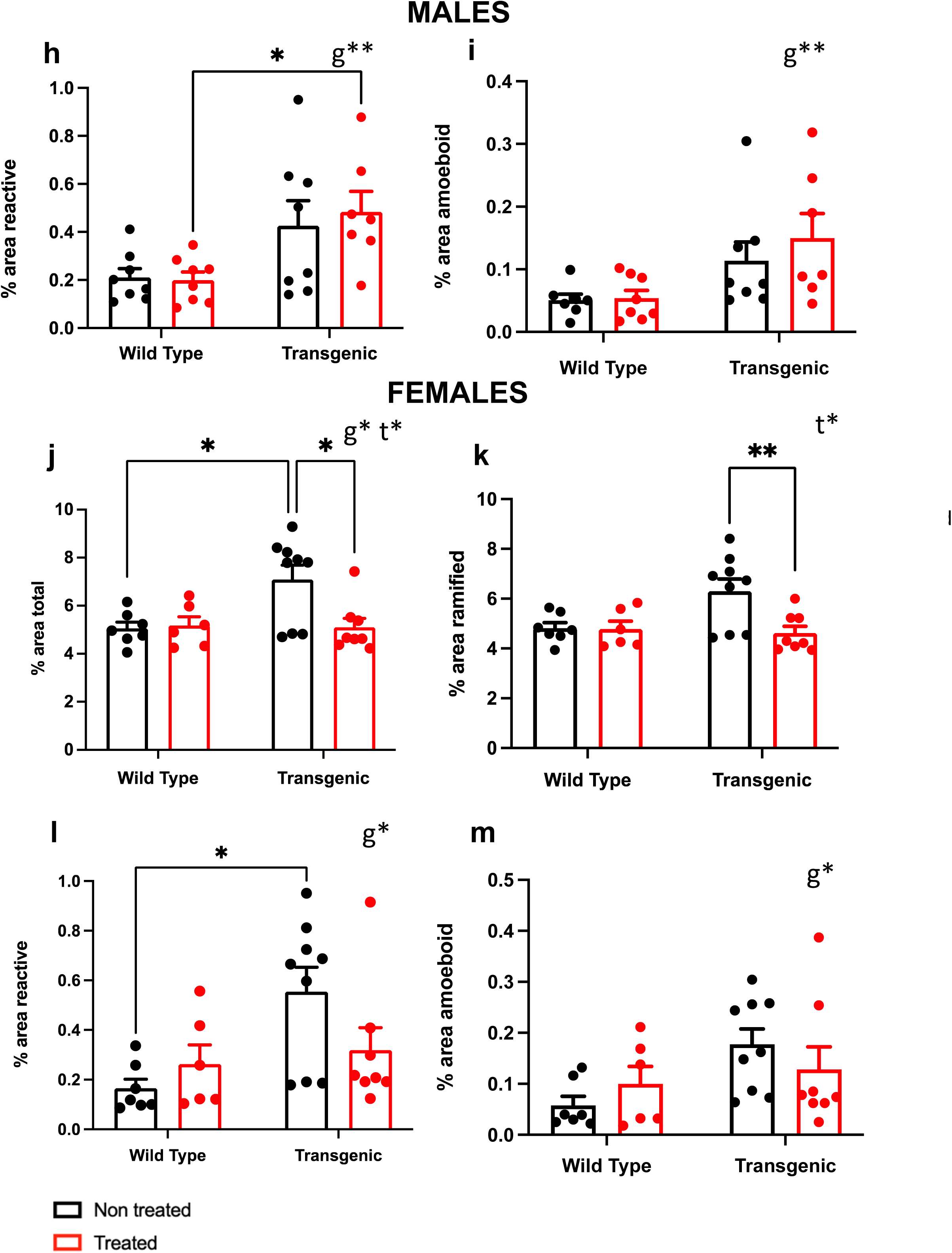
BT-11 treatment reduced total and ramified microglia levels in female TgF344-AD rats. **a,** Microglia immunohistochemical analyses with the Iba1 antibody (*green*) showed three distinct types of microglia morphology expressed by specific form factor ranges for circularity: ramified, reactive and amoeboid. Iba1 staining is shown for TGNT **b** and TGTR **c** males. Co-localization staining for Iba1 (*green*) Aβ plaques (*red*) and nuclei (DAPI, *blue*), is shown for TGNT **d** and TGTR **e** males. White arrows point to Aβ plaques. Bottom panels **f** and **g** represent the magnification of the respective small white boxes depicted in **d** and **e**. Scale bar = 1000µm **b-e**, and 10µm **f, g**. Male TgF344-AD rats had increased % area of reactive **h** and amoeboid **i** microglia in the dorsal hippocampus as compared to WT rats. Female TGNT rats had increased % area of Iba1 positive signal across the dorsal hippocampus for total **j**, reactive **l**, and amoeboid **m** microglia as compared to WTNT rats. BT-11 treatment reduced total **j** and ramified **k** microglia levels in female TGTR rats. Ordinary two-way ANOVA with Sidak’s *post hoc* tests were used for quantification. For simplicity, error bars are shown in one direction only. \**P*<0.05, ** *P* <0.01. TGNT – transgenic not treated; TGTR – transgenic BT-11 treated. CA = cornu ammonis, DG = dentate gyrus, SB = subiculum. t=treatment effect, g=genotype effect.

Male TgF344-AD rats had increased % area of reactive (p=0.0015) and amoeboid (p=0.0045) microglia in the dorsal hippocampus as compared to WT rats (Fig. 4h-i). There were also significant genotype differences across multiple subregions with TgF344-AD rats having increased microglia % area as compared to WT rats: CA1 reactive (p=0.0263), CA3 amoeboid (p=0.0370), DG reactive (p=0.0046), DG amoeboid (p=0.0509), with no changes in the subiculum (Supplemental Table 4). There were no significant differences induced by BT-11 treatment across all male groups (Supplemental Table 4).

Female TGNT rats had increased % area of Iba1 positive signal across the dorsal hippocampus for total microglia (p=0.0414), reactive (p=0.0146), and amoeboid (p=0.0402) as compared to WTNT rats (Fig. 4j-m, Supplemental Table 4). Significant genotype differences were also detected in the CA1, CA3, DG, and SB subregions (Supplemental Table 4). Additionally, BT-11 treated TgF344-AD females showed significantly reduced total microglia (p=0.0515) and ramified microglia (p=0.0282) levels in the dorsal hippocampus compared to TGNT female rats. Significant BT-11 treatment differences were also observed for CA1 ramified (p=0.0271), CA3 total (p=0.0446) and ramified (0.0308), and SB ramified microglia levels (p=0.0180), (Supplemental Table 4).

### Mature neurons are more abundant in the DG of TGTR than in TGNT male rats

To assess hippocampal neuronal numbers in male and female WTNT (males n=7; females n=7), TGNT (males n=7; females n=7), WTTR (males n=8; females n=6), and TGTR (males n=7; females n=7) rats, we used the NeuN antibody, which detects post-mitotic mature neurons (Fig. 5a-b). The images on the right represent the NeuN positive signal for quantification from immunofluorescent images. There was no significant genotype (g) or treatment (t) effects on neuronal counts/nm^2^ in the dorsal hippocampus of male rats [p=0.3572 (g), p=0.1017 (t)]. However, BT-11 treated TgF344-AD male rats compared to TGNT males had increased neuronal counts/nm^2^ in the DG hippocampal subregion (p=0.0192) (Fig. 5c). We analyzed NeuN counts across all hippocampal subregions, including the hilar region and the granule cell layer and are detailed in Supplemental Table 5.

**Fig. 5.**
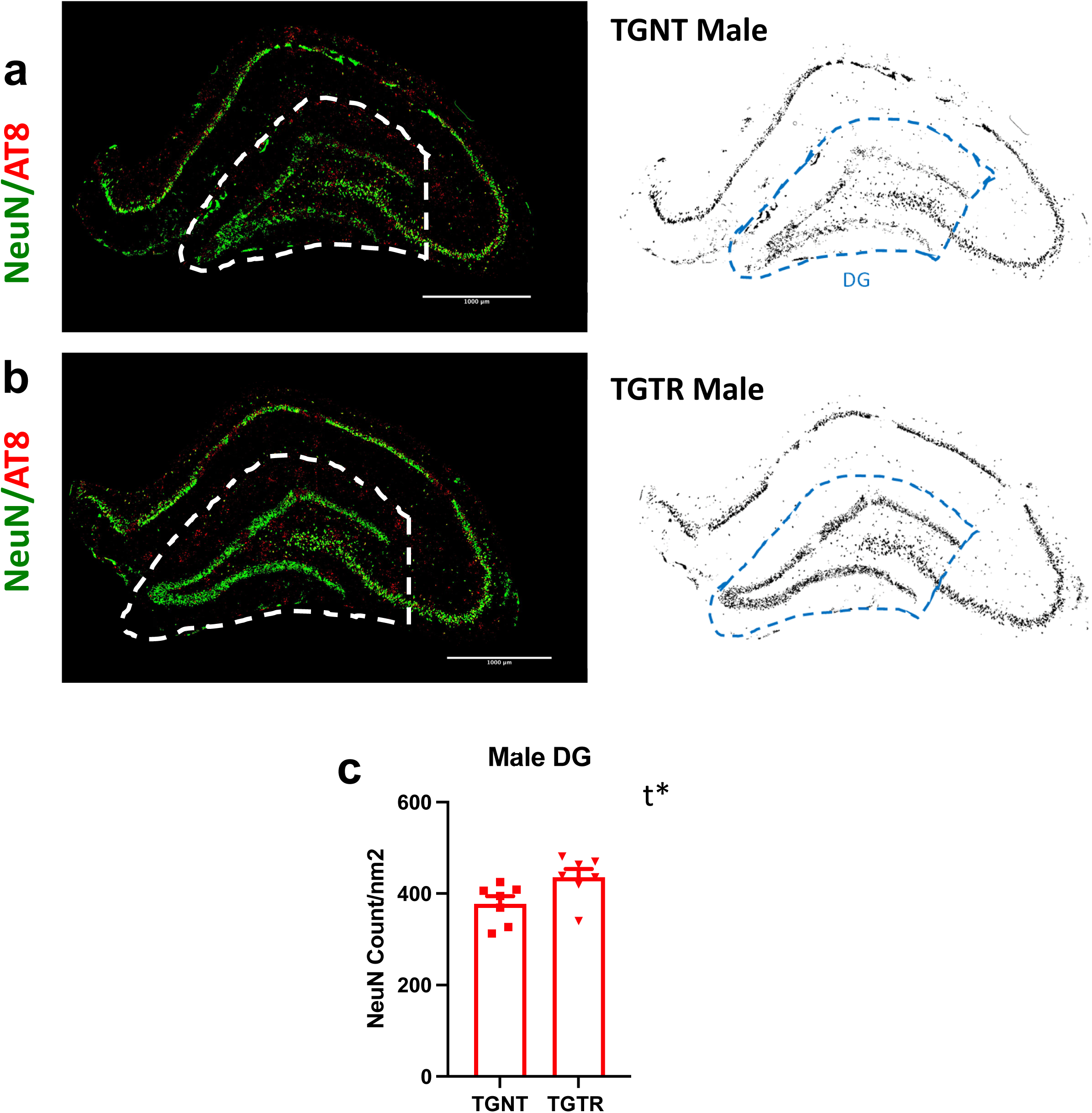
The number of mature neurons in the hippocampal dentate gyrus (DG) of TGTR males is higher than in TGNT males. NeuN staining (*green*) is shown for TGNT (**a,** n=7) and TGTR (**b,** n=7) males. Images on the right represent positive NeuN signal derived for quantification. Scale bars = 1000μm. TGTR males had increased NeuN counts/nm^2^ in the DG hippocampal subregion than TGNT males. Neurons were also stained for phosphoTau (*red,* AT8 antibody, Supplemental Fig. 5 shows phosphoTau staining alone). There were no significant differences of hyperphosphorylated tau levels in male rats across genotypes and treatments. Unpaired two-tailed *t*-tests with Welch’s corrections were used for quantification, \**P*<0.05. For simplicity, error bars are shown in one direction only. TGNT – transgenic not treated; TGTR – transgenic BT-11 treated. DG = dentate gyrus; t=treatment effect.

There were no significant genotype or treatment effects on neuronal counts/nm^2^ in the dorsal hippocampus of female rats [p=0.3485 (g), p=0.4544 (t)]. Additionally, there were no significant genotype or treatment effects across all hippocampal subregions (Supplemental Table 5).

### Hyperphosphorylated tau levels were equivalent in all groups of rats

We assessed hyperphosphorylated tau levels with the mouse monoclonal AT8 antibody (Supplemental Fig 5). There were no significant treatment (t) or genotype (g) differences of hyperphosphorylated tau levels in male rats [(p=0.1576 (g), p=0.9961 (t)] or female rats [(p=0.5119 (g), p=0.2030 (t)] of the four groups WTNT (males n=7; females n=7), TGNT (males n=7; females n=7), WTTR (males n=8; females n=6), and TGTR (males n=7; females n=7) (Supplemental Table 6a). Additionally, there were no significant treatment differences in hyperphosphorylated tau levels in male or female TgF344- AD rats across multiple subregions: CA1, CA3, DG, SB, HL, and GCL (Supplemental Table 6b-h).

### BT-11 induced changes in gene expression profiles and enriched pathways in TGTR vs TGNT rats

The differential expression profiles and enriched pathways in the dorsal hippocampi of TGTR vs TGNT male and female rats were assessed. Gene analysis generated a list of 11,671 genes for males and 12,561 genes for females. There were 195 genes in males and 77 genes in females that were differentially expressed in TGTR vs TGNT rats, considering the following parameters: *P* < 0.05, and a fold-change ≥ 1.5 or ≤-1.5. There were 12 overlapping genes between males and females (Fig 6a). In males, 123 genes were downregulated while 75 were upregulated, and three overlapping likely represented splice variants. In females, 56 genes were downregulated while 21 were upregulated (Fig 6a-c).

**Fig. 6.**
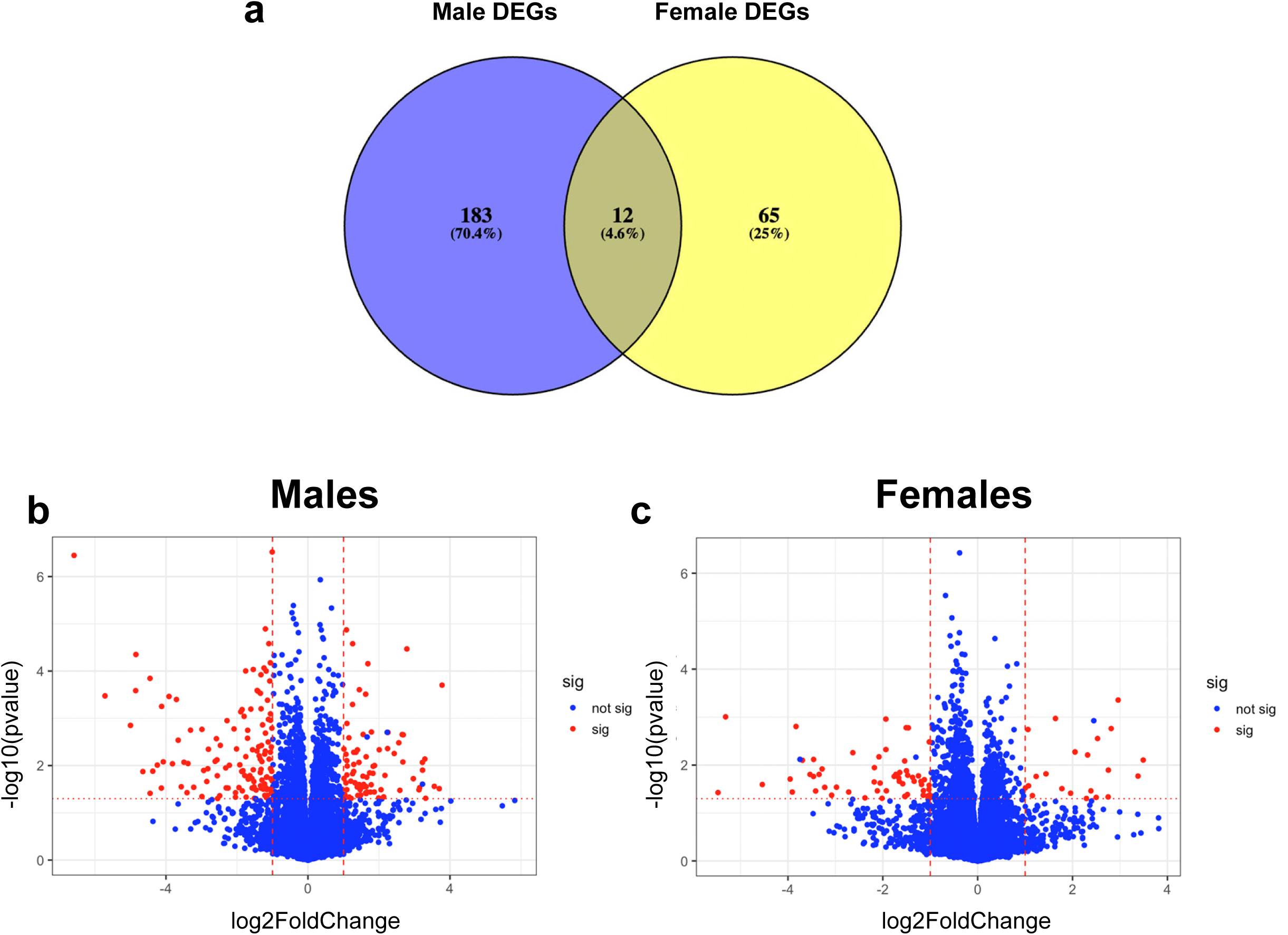
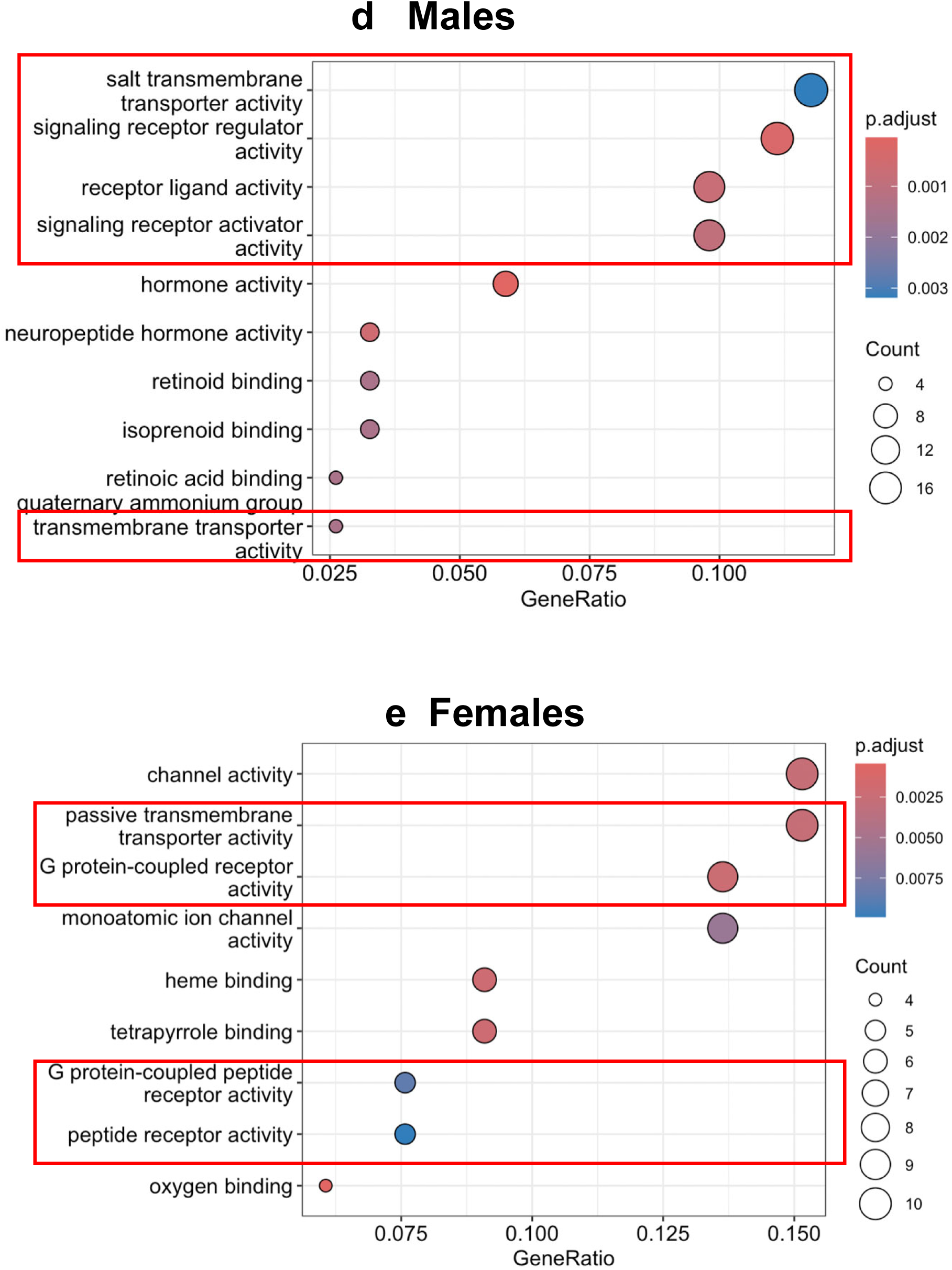
BT-11 treatment alters the transcriptome of male and female TgF344-AD rats. We used RNA sequencing (RNAseq) to analyze the transcriptomes of BT-11 treated and untreated TgF344-AD males and females separately. Five biological replicates for TGNT and TGTR males as well as females were analyzed. **a,** Venn diagram of differentially expressed genes (DEGs) shows a total of 195 genes for TGTR males (*lavender*), and 77 genes for TGTR females (*yellow*), with 12 genes overlapping between males and females (*brown*). **b,** Male and **c,** female volcano plots depicting significant (*red*) and non-significant (*blue*) DEGs from treated vs untreated TgF344-AD rats. Each point represents one gene. Y-axis, *P* values, x-axis, fold change. **d**, Male and **e**, female enriched pathway analysis of TGTR vs TGNT rats with the Gene Set Enrichment Analysis (GSEA) for transcriptional data. The size of circles represents the count of genes in the pathway differentially regulated. Color represents the *P* adjusted value of the pathway. Red boxes show relevant pathways relating to transporter and receptor activity. TGNT – transgenic not treated; TGTR – transgenic BT-11 treated.

Enriched pathways in males and females were determined by gene set enrichment analysis (GSEA, Fig 6d-e). Some enriched pathways for males included salt transmembrane transporter activity, signaling receptor regulator activity, receptor ligand activity, signaling receptor activator activity, and transmembrane transporter activity. Some enriched pathways for females included passive transmembrane transporter activity, G protein-coupled receptor activity, G protein-coupled peptide receptor activity.

### LANCL2 total, cytoplasmic and nuclear fraction levels were not altered in TGTR vs TGNT rats

Depending on post-translational modifications, LANCL2 is found in multiple locations of the cell, and this likely affects its function. We determined LANCL2 total, nuclear, and cytoplasmic fraction levels in dorsal hippocampi of male and female rats, by subcellular fractionation across each of the four groups WTNT, WTTR, TGNT, and TGTR, three rats per group (Fig. 7). LANCL2 levels associated with the plasma membrane were not detectable.

**Fig. 7.**
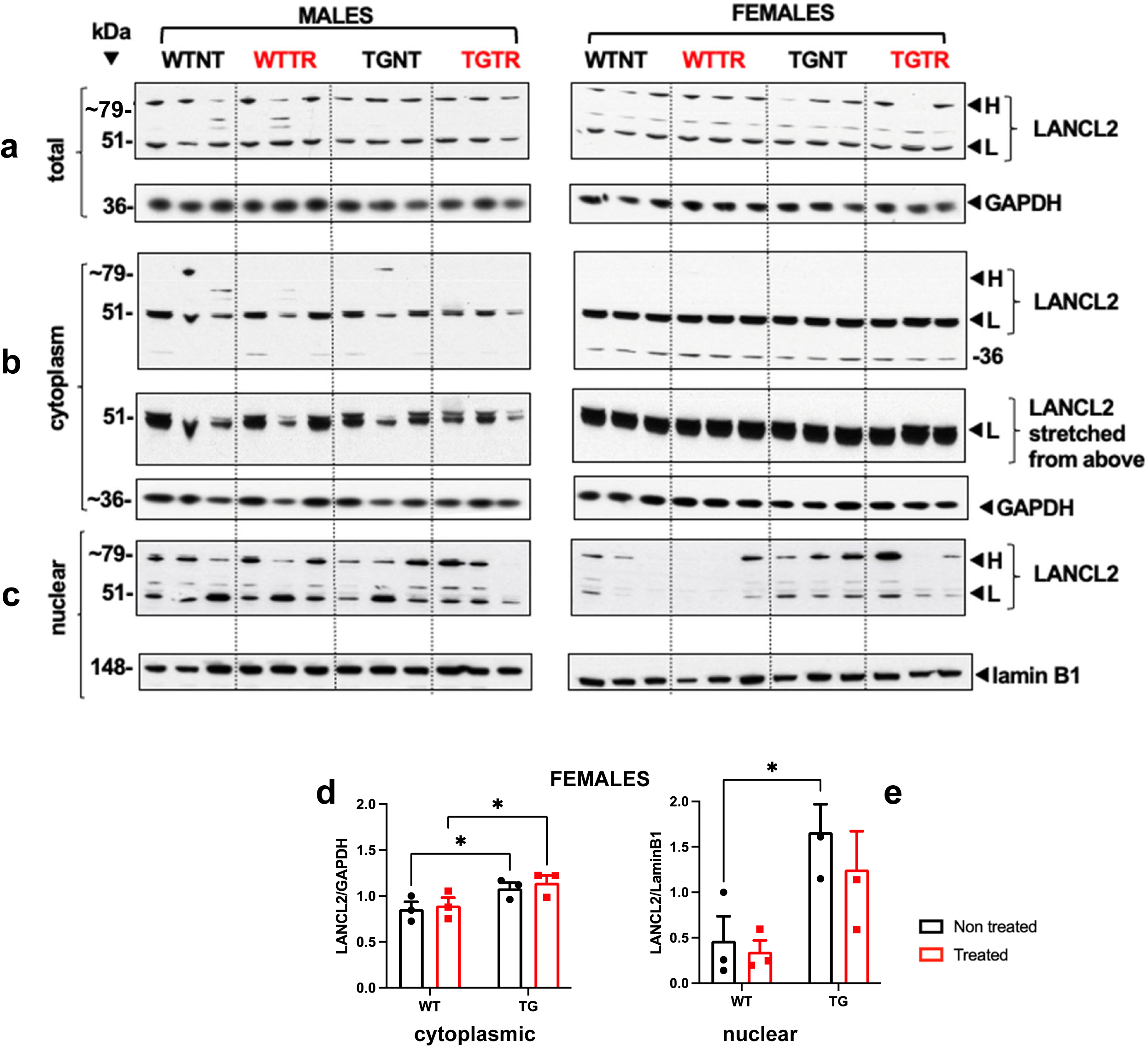
BT-11 treatment does not change LANCL2 total, cytoplasmic and nuclear levels in TgF344- AD male rats. Male and female rat dorsal hippocampal tissue homogenates were used to assess BT-11 treatment effects on the subcellular localization and levels of **a,** total, **b,** cytoplasmic and **c**, nuclear LANCL2 by western blot analysis. **d,** and **e,** Represent the percentage of the pixel ratio for LANCL2 over the respective loading controls, GAPDH for total and cytoplasmic fractions, and lamin B1 for the nuclear fraction. **d,** Analysis was performed for the four groups WTNT, WTTR, TGNT, TGTR at 11 months of age, three rats per group. Signal was quantified using relative density measurement on ImageJ. Only the data for females is shown. Values are means + SEM. Significance (asterisks shown on graphs) represent post-hoc effects following an ordinary two-way ANOVA with Sidak’s post-hoc tests. \**P* < 0.05. WTNT, wild-type not treated, TGNT, transgenic not treated, WTTR, wild-type BT-11 treated, TGTR, transgenic BT-11 treated; H = high molecular weight LANCL2; L = low molecular weight LANCL2.

Interestingly, besides the ∼51 kDa band corresponding to the reported LANCL2 molecular mass, an additional band was detected in the total lysate at ∼79 kDa (Fig. 7a). An increase of 28 kDa in LANCL2 suggests a post-translational modification. While this is a relatively high shift, there is the potential for ubiquitination or hyperphosphorylation to be involved.

In the cytoplasmic fraction of both males and females, a LANCL2 double band was detected at ∼51 kDa (Fig 7b). This could represent myristoylation, which is a known modification of LANCL2 that localizes the protein to the plasma or intracellular membranes. The addition of a myristol group would add ∼0.2 kDa per group to the protein. The molecular mass shift seen in the doublet could also be due to phosphorylation, which adds ∼0.8 kDa per phosphate group to a protein. There is also a ∼36 kDa band detected in the cytoplasmic fraction that could be a product of cleavage or degradation. In the cytoplasmic fraction, TgF344-AD females have significantly higher levels of LANCL2 than WT females regardless of treatment (Fig. 7d). There are no significant differences in LANCL2 levels in the male cytoplasmic fraction (Supplemental Table 7b).

The nuclear fraction (Fig. 7c) shows the ∼51 kDa LANCL2 band, and the ∼79 kDa band that is not detected in the cytoplasmic fraction (Fig 7a). It is possible that different post-translational modifications, such as ubiquitination, phosphorylation and myristoylation, occur on LANCL2 in different intracellular compartments. Female TgF344-AD rats have higher LANCL2 levels in the nuclear fraction than WT females (Fig. 7e). This is not seen in males (Supplemental Table 7C). The whole western blots with the LANCL2 signal are shown in Supplemental Fig. 6 – 11.

### BT-11 treatment did not alter the levels of pCREB in male and female TgF344-AD rats

To address the mechanisms mediating the effects of BT-11, we focused on the cAMP pathway because: (a) LANCL2 activation affects cAMP signaling leading to CREB phosphorylation by PKA ^11,12^ and (b) our transcriptome analysis revealed enrichment of signaling receptor pathways with BT-11 treatment (see above under “gene expression”). The same total lysate samples described above for male and female rats across the four groups WTNT, WTTR, TGNT, and TGTR were analyzed for total CREB and pCREB (Supplemental Fig. 12 - 15). There were no significant differences in pCREB levels across groups (Supplemental Table 8) for male and female rats.

### LANCL2 is detected in oligodendrocytes in dorsal hippocampi of TgF344-AD and WT rats

The cell type localization of LANCL2 in the dorsal hippocampus is only predicted by high throughput transcriptomic analyses.^13^ To further investigate this question, IHC was performed on dorsal hippocampal sections including the surrounding *corpus callosum.* Sections were co-immunostained with the LANCL2 antibody together with each of the cell type specific antibodies: NeuN for neurons, Iba1 for microglia, GFAP for astrocytes, or Olig2 for oligodendrocytes (Supplemental Table 1). WTNT and TGNT male rats (three rats per group) were included in the analysis. LANCL2 cell type localization was similar across each group.

LANCL2 signal was clearly detected in oligodendrocytes, although its signal was also found near astrocytes, neurons, and microglia suggesting possible interactions (Fig 8a - p). Olig2 is a transcription factor, and its signal is mainly localized in the nucleus while LANCL2 stain is found mostly in the cytoplasm leading to the signal appearing as red LANCL2 wrapped around the green Olig2 of the same cell (Fig. 8h and 8p). ^13,14^

**Fig. 8.**
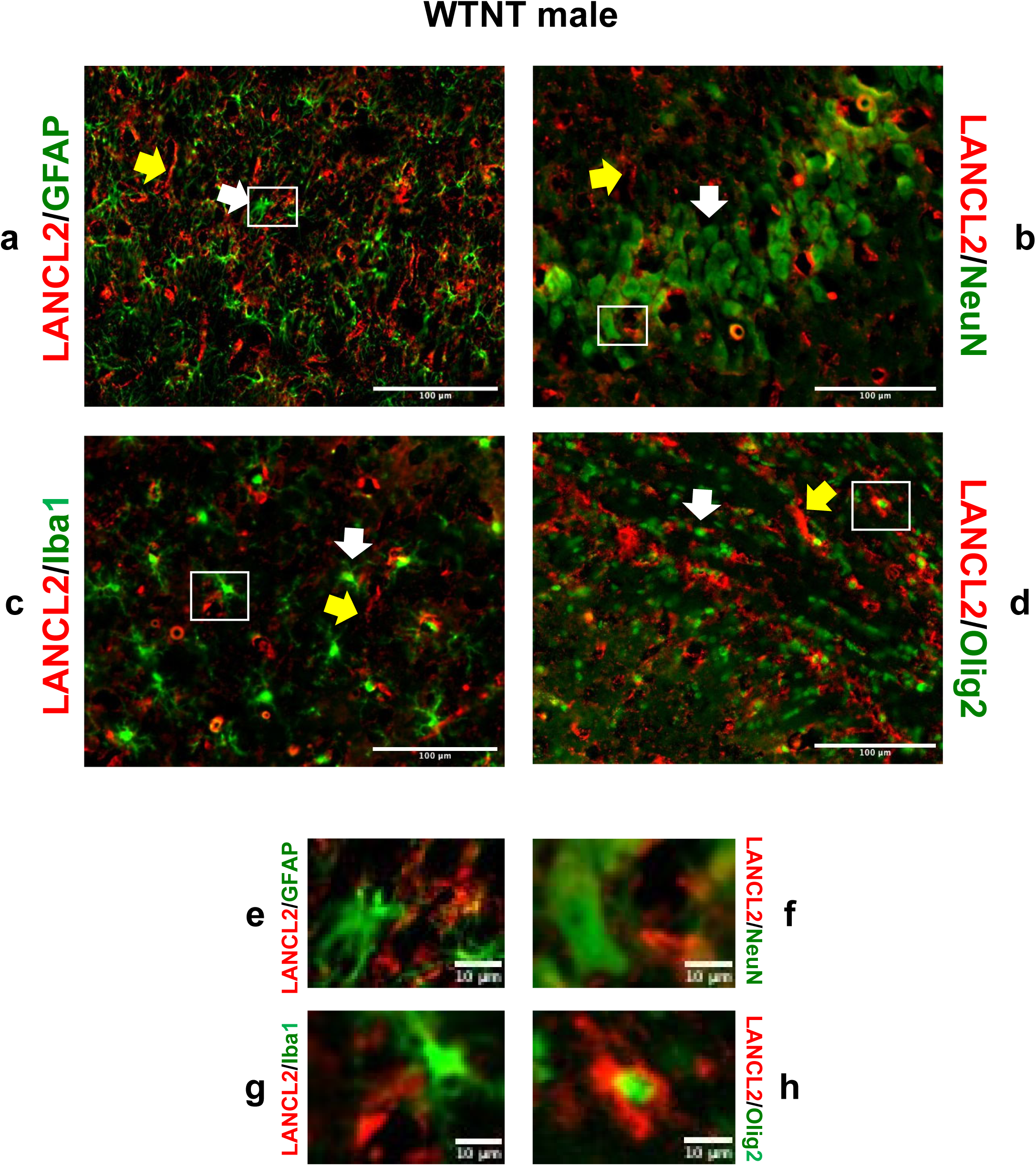
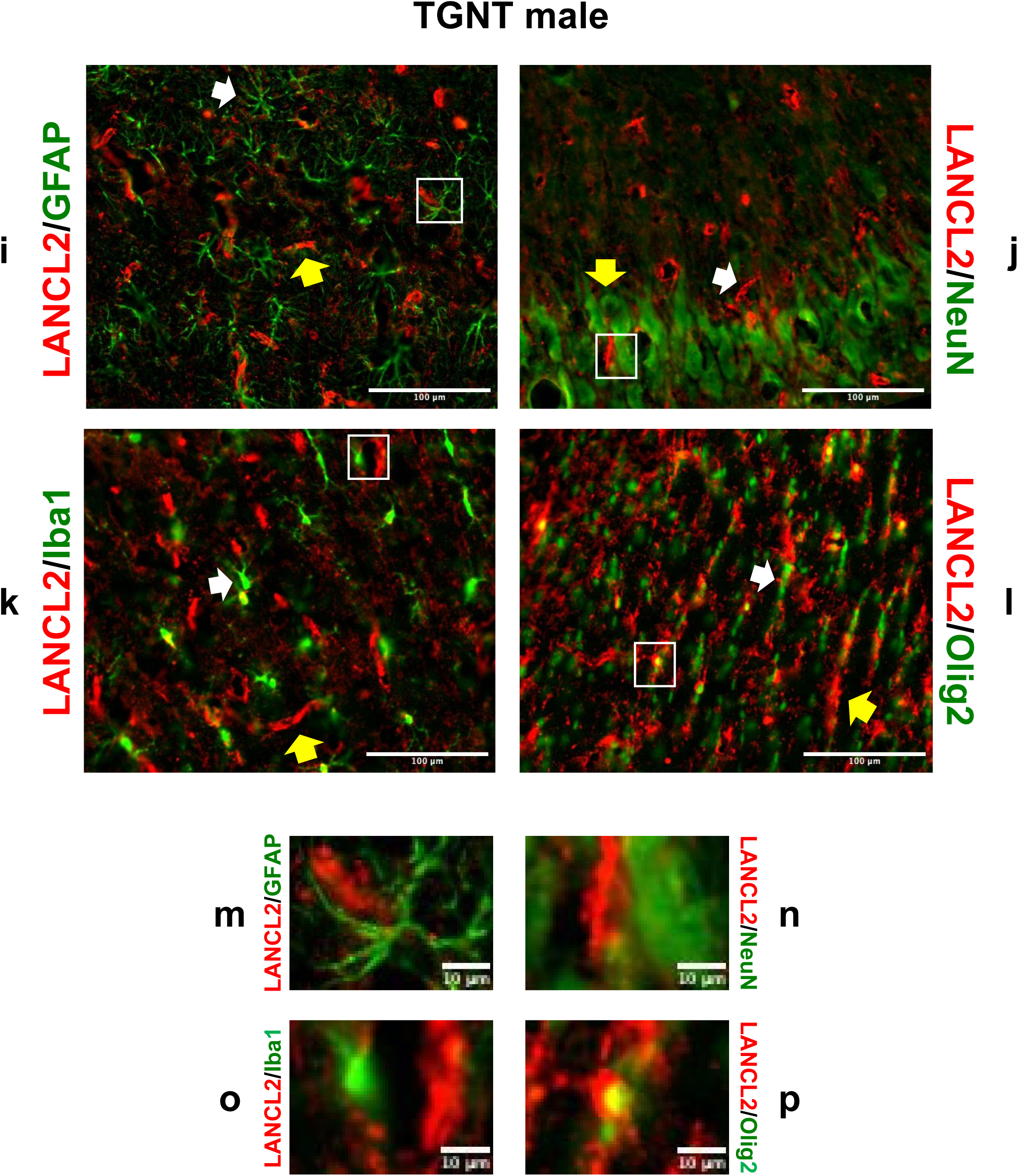
LANCL2 is localized in oligodendrocytes in the dorsal hippocampus of WT and TgF344-AD male rats. LANCL2 is detected in oligodendrocytes found in the dorsal hippocampus and neighboring corpus callosum of WTNT (**a-h**) and TGNT (**i-p**) male rats. Co-localization of LANCL2 (*red*) with different cell specific markers (*green*): **a, i,** GFAP for astrocytes, **b, j**, NeuN for neurons, **c, k,** Iba1 for microglia, and **d, l,** Olig2 for oligodendrocytes. Bottom panels (**e-h**) represent the magnification of the respective small white boxes depicted in **a-d**. Bottom panels (**m-p**) represent the magnification of the respective small white boxes depicted in **i-l**. Scale bar = 1000µm (**a-d, i-1**), and 10µm (**e-h, m-o**). LANCL2 *red* staining is wrapped around the *green* Olig2 nuclear stain, clearly shown in **h a**nd **p**. White arrows point to the cell type staining shown in *green*, yellow arrows point to LANCL2 staining shown in *red*. WTNT, wild type not treated, TGNT, transgenic not treated.

## Discussion

Overall, our studies investigated the effects of long-term oral treatment with the LANCL2 activator BT-11, on cognitive performance and AD pathology in male and female TgF344-AD rats and WT controls. Moreover, we characterized LANCL2 cellular and subcellular localization in the hippocampus.

Hippocampal-dependent spatial learning was assessed with the aPAT.^15^ As previously described, rats learn to avoid a shock zone by associating it with cues placed around the behavior testing room. This makes the task hippocampal-dependent, given that the hippocampus plays a key role in remembering past locations.^16^ The 11-month TgF344-AD male rats showed a significant deficit in performance as compared to WT controls. Interestingly, the same deficit was absent in the female rats. The initial report of this model found no significant behavioral sex differences.^8,9^ However, more recent studies looking at a range of behavioral tasks demonstrated significant differences in cognitive performance between male and female TgF344-AD rats, such as females showed no deficit in a buried food task at 12 months of age, while males exhibited a significant deficit.^9^ Additionally, our previous studies demonstrated that female TgF344-AD rats outperform males in the aPAT with no significant behavior changes in either group at 9 months of age^29^. There are conflicting results on the behavioral characterization of the TgF344-AD rats that seem to relate to the task being evaluated. For example, in a study that focused on females, TgF344-AD rats showed spatial memory impairment in a Morris water maze at 12 months, however showed no significant deficits at 12 months during the acquisition phase of a radial arm water maze task.^17^ These studies suggest that the behavior task chosen to assess spatial learning influences the behavior performance. Thus, in future studies it will be important to use a variety of behavioral tasks to show changes in spatial learning.

While the male TgF344-AD rats performed worse on the aPAT than WT rats, the TGTR rats performed significantly better than the TGNT rats. These results show that BT-11 treatment starting early at 5 months of age, mitigated the spatial learning deficits showed in male TGNT rats. Many AD patients maintain cognitive function demonstrating a potential ‘resilience’ to AD pathology, and our previous study showed improved behavior performance in Tg-F344AD females despite increased pathology as compared to males.^18,19^

There was a significant reduction of hippocampal Aβ plaque load in TGTR compared to TGNT males. In addition, the number of mature neurons in the hippocampal dentate gyrus (DG) of TGTR males was higher than in TGNT males, supporting a relationship between neuronal numbers and improved spatial memory. This shows that BT-11 treatment likely ameliorates cognitive deficits by reducing classical AD pathology in male TGTR rats. There was no BT-11 treatment effect on Aβ plaque load or neuronal levels in female transgenic rats, corresponding to no effect of the drug on spatial memory.

Significantly, a key factor limits the drug translatability of female rats as compared to male rats for both behavior and pathology effects. Female rats, unlike humans, undergo estropause instead of menopause. Estropause occurs around 9-12 months of age, corresponding to when our rats were evaluated. Unlike menopause where there is a significant loss of estrogen signaling, estropause is characterized by moderate to high levels of estrogen.^20^ Importantly, estrogen is neuroprotective, increasing cognitive performance and reducing oxidative stress.^21^ This creates a potential source of protection against neurodegeneration in the TgF344-AD females, not present in males and not recapitulated in humans. Since we are focused on the effect of drugs with an eventual goal of translation, future studies could include models to induce menopause such as ovariectomy or drug induced menopause.^20^ However, these methods can introduce additional variables to the study. For the sake of translatability, it is still important to evaluate both male and female rats despite this key difference. The sex-dependent effects of BT-11 may be explained by sex-specific mechanisms mediated by BT-11, as well as possible differences in drug metabolism by males and females. Translational science researchers must continue to address this issue.

Microglia analysis showed an increase in reactive and amoeboid microglia in males in the dorsal hippocampus of TgF344-AD rats compared to WT rats. BT-11 treatment had no effect on microglia in males. However, female TgF344-AD rats showed an increase in total, reactive and amoeboid microglia in the dorsal hippocampus as compared to WT rats, as well as a reduction in total and ramified microglia with BT-11 treatment. BT-11 treatment in female TgF344-AD rats also caused a reduction in ramified microglia in the CA1, CA3 and subiculum. Ramified microglia are commonly associated with surveillance.^22^ There were no significant treatment effects on reactive or amoeboid microglia, which are known to be involved in phagocytic activity and associated with disease.^23^ There are many ways to analyze microglia and microglial signaling including presence of inflammatory markers such as CD68 as well as utilizing single cell sequencing to assess the transcriptomic nature of different types of microglia.^23–25^ Future studies focusing on more detailed and multiple ways of assessing microglial activity and function should provide additional support and explanations for our studies. Moreover, as previously mentioned, BT-11 reduces inflammation in the gut.^26^ It is possible that BT-11 influences inflammation in TgF344-AD rats in ways that may not be demonstrated by analyzing microglial levels. It may relate to an overall reduction in systemic inflammation, or presence of other immune cells, such as T cells, and other signaling pathways.

There were no genotype or treatment changes in hyperphosphorylated tau in male or female rats. Previous studies on this TgF344-AD rat model suggest that the 11-month timepoint is too early to demonstrate tau pathology, proposed in the initial report to be detected at 16 months of age.^8^ A more recent study showed increased levels of hyperphosphorylated tau in rats aged 18-21 months and no change in rats 6-8 months of age.^8,27^

So far, we showed that oral treatment with BT-11 ameliorated cognitive deficits and reduced some of the pathological hallmarks of AD in the TgF344-AD rat model. Based on these findings, we investigated the molecular and cellular mechanisms potentially mediating the effects of BT-11 treatment on the dorsal hippocampus of WT and TgF344-AD rats. Specifically, we focused on the dorsal hippocampal expression as well as the intracellular and cell type localization of the BT-11 target LANCL2, to gain a better understanding of its role in AD.

Firstly, we investigated potential CNS effects of BT-11, by comparing mRNA expression in the dorsal hippocampus of TGTR vs TGNT rats. Treating TgF344-AD rats orally with BT-11 led to enrichment in G-protein coupled receptor and other signaling receptor pathways. This suggests that activating LANCL2 with BT-11 likely works by these signaling mechanisms in the brain in addition to systemic effects.

Secondly, we explored LANCL2 protein expression in the brain at the subcellular level. In the dorsal hippocampal tissue, as expected, LANCL2 was detected as a 51 kDa protein band in both the nuclear and cytoplasmic fractions. Interestingly, an ∼51 kDa doublet was found in the cytoplasmic fraction, and an additional ∼79 kDa band was visible in the nuclear fraction. These other bands could hint at post-translational modifications.

The cytoplasmic ∼51 kDa doublet could correspond to LANCL2 myristoylation.^28^ LANCL2 myristoylation adds a hydrophobic handle to the protein and induces its localization to cellular membranes, leading to it being identified as a non-transmembrane G-protein coupled receptor.^29,30^ Additionally, when localized to a cellular membrane and activated by its endogenous ligand, ABA, LANCL2 activates adenylyl cyclase, thus increasing cAMP levels.^31,32^ This was demonstrated in human red blood cells and granulocytes, but to my knowledge, has not been previously shown in the brain.

In the nuclear fraction, the ∼79 kDa LANCL2 upper band could be the result of multiple phosphorylation and/or polyubiquitination translational modifications. Treatment of glioblastoma cells with ABA, the endogenous LANCL2 ligand, triggered its translocation to the nucleus.^33^ Phosphoproteomic profiling of glioblastoma cells showed that ABA-induced LANCL2 phosphorylation at Tyr295 drives its activation and nuclear localization.^33^ Using the PhosphoSitePlus tool (https://www.phosphosite.org) as in ^33^, we identified potential LANCL2 phosphorylation, acetylation, and ubiquitination sites based on broad mass spectrometry studies of heart, brain tissue and multiple cell lines. It is possible that some of the other bands detected in our western blot experiments represent known or yet to be determined post-translational modifications. Future studies looking at LANCL2 post-translation modifications in conjunction with subcellular localizations in the context of AD, could lead to the clarification of the role of LANCL2 in the disease.

Thirdly, we showed for the first time to our knowledge that LANCL2 is present in dorsal hippocampal oligodendrocytes, which engage in the formation of myelin sheaths to speed up neural transmission and provide metabolic support to axons. Oligodendrocyte loss occurs in the context of AD, however much is unknown about the characterization of this pathology.^34^ The fact that LANCL2 is found in oligodendrocytes in the dorsal hippocampus could point to a previously unstudied brain-specific function of LANCL2. Interestingly, a recent study described an intrinsic signaling pathway involving a glutathione-s-transferase that modulates oxidative stress in premyelinating oligodendrocytes.^35^ Given the identification of LANCL2 as a glutathione-s-transferase, this could be a possible role LANCL2 plays in oligodendrocyte function.^36^ This oligodendrocyte finding merits future studies such as elucidating at what stages of oligodendrocyte development LANCL2 is present, if there are changes in expression of LANCL2 on oligodendrocytes in ageing or AD conditions, and assess the role of activating LANCL2 in oxidative stress in the context of AD.

BT-11 was predicted to cross the blood brain barrier suggesting that it can act locally, as it seems to have been detected in rat brain.^5^ Thus, BT-11 may be working both at the CNS and systemically, as BT- 11 treatment reduced inflammation in the gut.^2,26,37^ It is well known that gut inflammation affects brain pathology ^50^. The brain is not immune-privileged and systemic inflammation influences AD pathology. ^38,39^ It is possible that after treatment with BT-11, the vagus nerve senses a reduced inflammatory state in the gut and transmits signals to the brain that ultimately lead to modulation of neuroinflammation. To address this potential mechanism, future studies should look at levels of inflammatory markers in the brain, gut, and blood after oral treatment with BT-11.

In conclusion, our studies highlight the therapeutic potential of targeting LANCL2 to improve cognitive behavior and reduce Aβ plaques and neuronal loss, especially in male AD patients. Our data suggest multiple mechanisms by which LANCL2 activation by BT-11 could ameliorate cognition and reduce AD pathology: (1) Showing that BT-11 treatment enriches G-protein signaling pathways, points to the possibility of a previously identified mechanism involving cAMP in human blood cells and granulocytes, being present in the hippocampus. While BT-11 treatment did not change LANCL2 expression, it likely changes its activity levels leading to changes in downstream effectors. (2) Determining that, in the hippocampus, LANCL2 is present in both the nuclear and cytoplasmic fractions with potential subcellular specific post-translational modifications that could influence its role in different subcellular compartments. (3) Finding that LANCL2 is present in oligodendrocytes of the dorsal hippocampus of WT and TgF344-AD rats points to a role in supporting oligodendrocyte function in both normal and disease contexts. As LANCL2 is a glutathione-*s*-transferase, its activation could reduce oxidative stress in oligodendrocytes. Overall, given the multifactorial nature of AD and the variety of potential mechanisms by which LANCL2 activation could improve pathology, we show that LANCL2 has significant translational potential as an AD target and BT-11 as an AD therapeutic.

## Materials and Methods

### TgF344-AD rat model

TgF344-AD transgenic rats express the human APP Swedish (APPswe) and Δ exon 9 presenelin-1 (PS1ΔE9) mutations driven by the prion promoter, at 2.6- and 6.2-fold higher levels than the respective endogenous rat proteins.^8^ These rats exhibit AD-like pathology and navigational deficits similar to those found in pre-clinical AD, and in an age-dependent progressive manner.^40^ TgF344-AD rats were purchased from the Rat Resource and Research Center (Columbia, MO). TgF344-AD and WT rats of both sexes were housed in pairs and maintained on a 12-hour light/dark cycle with food and water available *ad libitum*. All animal procedures were performed in compliance with the relevant guidelines and regulations of the Institutional Animal Care and Use Committee (IACUC) at Hunter College. All experimental procedures were approved by the IACUC and agreed with the standards outlined in the ARRIVE guidelines.

### Experimental Design

Our aim was to figure out if BT-11 could mitigate, prevent, or delay AD symptoms. BT-11 treatment started at 5 months of age (prior to the known onset of pathology) and continued for 6 months until 11 months of age, when the TgF344-AD rats show moderate AD-pathology (Fig. 1a, experimental design). A total of 124 rats across multiple cohorts were included in these studies. Males: WTNT n=14, WTTR n=13, TGNT n=17, TGTR n=13, and Females: WTNT n=28, WTTR n=12, TGNT n=16, TGTR n=11.

Rats were treated with 8 mg/kg bw/day of BT-11, based on efficacy studies at 8 mg/kg bw/day in multiple mouse models of IBD, and safety at 80 mg/kg per day in Sprague Dawley rats.^4,41^ BT-11 was delivered orally through *ad libitum* 5001 Purina rodent chow from Research Diets, Inc. Efficacy assessment included treatment with 2 types of chow: BT-11 and BT-11-free (control) chow x 2 genotypes (WT and TgF344-AD) x 2 sexes (males and females).

At 11 months of age, all rats were evaluated for hippocampal-dependent spatial memory. Following cognitive behavior assessment, rats were sacrificed, and brains were isolated, bisected and prepared for IHC, RNA sequencing (RNAseq), and western blot analyses.

### Spatial memory assessment

All rats were tested at 11 months of age with the active place avoidance test (aPAT), as previously described^15,42^. The aPAT assesses hippocampal-dependent spatial learning and memory function relevant to AD.^15^ The hippocampus plays a role in spatial memory, and it is one of the first brain regions affected by AD. The PAT is active, meaning that the rodent must take part.

Performance in the behavior test was analyzed based on the following measures: percent time spent in shock zone, latency to first entrance of shock zone, number of entrances, number of shocks administered, and maximum time spent avoiding the shock zone. A well-performing rat, indicative of increased learning and spatial memory, will spend less time in the shock zone over the course of 6 trials. As in our previous studies, rats generally learned during the first 3 trials (*early acquisition, trials 1-3*) and then reached an asymptote at which their performance levels off during trials 4-6 (*asymptotic performance, trials 4-6)*.^10,18,43^

### Immunohistochemistry

Tissue collection, preparation, and left hippocampal immunohistochemistry (IHC) were performed in 11-month-old rats, as previously described^42,44^. Hippocampal sections chosen for IHC analysis corresponded to Figures 60 – 70 of the rat brain atlas (bregma - 3.36 through - 4.44).^45^ Sections were viewed under a wide-field fluorescence microscope (Zeiss AxioImager M2) using a Zeiss AxioCam MRm Rev. 3 camera connected to a motorized stage. The software AxioVision 4 module MosaiX was used to capture whole hippocampal region mosaics (5x, 10x and 20x magnifications). Each channel was analyzed to an antibody specific threshold. DSRed set was used for Aβ imaging, GFP spectrum filter set was used for Iba1 imaging, DAPI set was used for DAPI imaging. Exposure time for each channel was kept consistent among sections. For each captured image, ZVI files were loaded onto FIJI (Fiji Is Just ImageJ, NIH, Bethesda, MD) and converted to .tif files for future use in optical density and co-localization analyses.

Hippocampal subregions (CA1, CA3, DG and subiculum) were isolated, cropped, and saved as .tif files for use in pixel-intensity area analyses as described in.^46^ Three tissue sections per treatment/genotype group were used for quantification. Nonspecific background density was corrected using ImageJ rolling-ball method.^47^ Primary and secondary antibodies are listed in Supplemental Table 1.

Images were analyzed to extract the positive signal from each image with custom batch-processing macroscripts created for each channel/marker. Pixel-intensity statistics were calculated from the images at 16-bit intensity bins.^48^ Positive signal areas were measured, and masks were created and merged when co-localization analyses were required. Co-localization measurements were achieved by measuring the overlap of two merged masks from an analyzed image crop.

Ramified, reactive, and amoeboid microglia were analyzed by circularity based on the ImageJ form factor as previously described. ^42,44^ Microglia exhibit a remarkable variety of morphologies that can be associated with their functions.^49^ Using binary images of individual microglia silhouettes, activated microglia were distributed into three different groups according to their form factor (FF) value, which was defined as 4π X area/perimeter^2^: *<u>ramified</u>*, FF: 0 to 0.499; *<u>reactive</u>*, FF: > 0.5 to 0.699, and *<u>amoeboid</u>*, FF > 0.70 to 1. Images stained with the Iba1 antibody were used for morphological analyses, to characterize microglia into the three distinct shape classes. Particles within 50–300 μm^2^ were chosen for FF analyses. Quantities of each microglia class were collected and analyzed independently and in ratios of amoeboid normalized to ramified and then compared across treatment/genotype.

### RNA sequencing analyses

Right hippocampi from the same rats used for IHC, were analyzed for gene expression by RNA sequencing (RNAseq) at the UCLA Technology Center for Genomics and Bioinformatics (Los Angeles, CA). Samples from five male TGNT and five TGTR were compared, and the same was done for female rats. Libraries for RNAseq were prepared with the KAPA mRNA-Seq Hyper Prep Kit. cDNA was prepared and amplified, and sequencing was performed on Illumina NovaSeq6000 for a PE 2×50 run. Data quality was checked using Illumina SAV and demultiplexing was performed with Illumina Bcl2fast1 v2.19.1.403 software. Reads were mapped by STAR 2.7.9a and read counts per gene were quantified using the rat genome rn6. Read counts were normalized by median ratio. Differentially expressed genes between TGNT and TGTR for each sex were determined using the DESeq2 program on R.^50^ Normalized read counts were analyzed for fold-change, *p* values, and *p* adjusted values (based on false discovery rate) for each gene. Gene set enrichment analysis including genes with p value <0.05 and fold change >1.5 or <-1.5 was used to generate enriched pathways.

### Subcellular fractionation

Approximately 50mg of frozen right hippocampal tissue were added to 4μl of homogenizing buffer (0.5% 10X protein inhibitor cocktail in TEE buffer) per mg of tissue. The tissue was homogenized using 20 pumps on the Cole Palmer homogenizer at 40rpm at room temperature. Approximately half of the total homogenate was aliquoted into an Eppendorf tube, labeled total lysate, and stored at −80°C. The other half of the homogenate was placed in an Eppendorf tube labeled ‘nuclear’ and centrifuged for 5 min at 3,000 g at 4°C. The supernatant was transferred to an ultracentrifuge tube and spun at 100,000 g for 30 min at 4°C in the ultracentrifuge. While the ultracentrifuge was running, 100 µl of homogenizing buffer was added to the nuclear Eppendorf tube and it centrifuged again at 3,000 g for 10 min to remove debris. The supernatant was discarded, and the pellet resuspended in 100 µl of homogenizing buffer, labeled nuclear fraction and stored at −80°C. The supernatant from the ultracentrifuge tubes were transferred to an Eppendorf tube, labeled cytoplasmic fraction, and stored at −80°C.

This procedure was performed for three female and three male rats of the following four groups: WTNT, WTTR, TGNT, and TGTR. This procedure yielded three cellular fractions, total lysate, nuclear and cytoplasmic fractions all used for western blot analyses.

### Western blotting

Protein concentrations for total, nuclear, and cytoplasmic fractions were determined with the BCA assay. 30μg of sample were loaded on a Novex Tris-Glycine 4-12% gel, ran for approximately one hour and 50 min, and transferred to nitrocellulose membranes with the iBlot® dry blotting system (Life Technologies) for 6.5 min. Membranes were washed with TBS/Triton and blocked with 5% BSA. Membranes were then probed overnight at 4°C with primary antibody solutions in 1% BSA, followed by secondary antibodies for one hour, prior to developing with an enhanced chemiluminescence (ECL) substrate (SuperSignalTM West Pico PLUS, ThermoFisher #34580), and detected by autoradiography (film from Midwest Scientific, cat# XC6A2). Primary and secondary antibodies are listed in Supplemental Table 1. ImageJ software was used for semi quantification of protein bands by densitometry. Loading controls used were GAPDH for total and cytoplasmic fractions and lamin B1 for nuclear fractions.

### Statistical analysis

Statistical analysis was performed with GraphPad Prism 9 or 10 (GraphPad Software, San Diego, CA). All *p* values and relevant statistics are shown on graphs and supplemental tables. Data are the mean ± standard error of the mean (*SEM*) with *n* being the number of rats. Three-way ANOVAs were used to determine significant effects across independent variables for behavior prior to two-way repeated measures ANOVAs across all four groups for both males and female separately (WTNT, TGNT, WTTR, TGTR), as well as for WTNT vs. TGNT, TGNT vs. TGTR, WTNT vs. WTTR, and WTTR vs. TGTR, followed by Tukey or Sidak post-hoc tests to assess differences across individual trials (Supplemental Tables 2a and b and 9).

For IHC analyses unpaired two-tailed *t-tests* were used to assess Aβ % area between TGNT and TGTR rats (Supplemental Table 3). Additionally, two-way ANOVAs were used to assess tau count/nm^2^ for WTNT, TGNT, WTTR, and TGTR rats for the dorsal hippocampus (Supplemental Table 6a). Unpaired t-tests were used to assess tau count/nm^2^ between TGNT and TGTR rats across multiple subregions (Supplemental Tables 6b-h). For all microglia analyses two-way ANOVAs with Tukey multiple comparisons tests were performed (Supplemental Tables 4a-t). For NeuN IHC analyses two-way ANOVAs with Tukey multiple comparisons were performed for WTNT, TGNT, WTTR, and TGTR rats (Supplemental Tables 5a-g).

For western blotting two-way ANOVAs followed by Tukey’s multiple comparison tests were used to compare normalized densities of proteins across four groups for both males and females (WTNT, WTTR, TGNT, TGTR, Supplemental Tables 7a-c and 8).

## Author contributions

E.B., P.S., P.R., and M.F.P., conceived the project and designed the experiments. L.X. performed the computational analysis supporting BT-11 treatment for AD. E.B. performed all experiments and analyzed the data. E.B. and M.F.P wrote the manuscript. P.S. and P.R edited the manuscript. All authors approved the manuscript for submission.

## Data Availability

The datasets generated and/or analyzed during the current study are available in the NIH GEO repository, ACCESSION NUMBERS: GSE280023 and GSE280024

