## Supplemental Tables and Figures for "BT-11 repurposing potential for Alzheimer’s disease and insights into its mode of actions"

**Supplemental Table 1 – Antibodies used for IHC and WB analyses of hippocampal tissue.**

| Antibody | Company | Catalog Number | Species | Dilution | Assay |
|---|---|---|---|---|---|
| | | Primaries | | | |
| LANCL2 | Novus Bio | NBP1-83339 | Rabbit | 1:2000 | WB |
| Aβ (4G8) | Biolegend | 800708 | Mouse | 1:1000 | IHC |
| Iba1 | Wako | 019-19741 | Rabbit | 1:500 | IHC |
| Phospho Tau (AT8) | Thermo Fisher Sci | MN1020 | Mouse | 1:500 | IHC |
| Lamin B1 | Thermo Fisher Sci | PA5-19468 | Rabbit | 1:1000 | WB |
| GAPDH | Millipore Sigma | MAB374 | Mouse | 1:2000 | WB |
| NeuN | Millipore Sigma | ABN91 | Chicken | 1:200 | IHC |
| GFAP | Millipore Sigma | AB5541 | Chicken | 1:200 | IHC |
| Iba1 | Synaptic Systems | 234006 | Chicken | 1:500 | IHC |
| Olig2 | Millipore Sigma | MABN50 | Mouse | 1:500 | IHC |
| LANCL2 | Invitrogen | PA5-72884 | Rabbit | 1:250 | IHC |
| | | Secondaries | | | |
| Alexa Fluor 568 Goat anti-rabbit IgG (H+L) | Thermo Fisher Sci | A-11036 | Rabbit | 1:250 | IHC |
| Alexa Fluor 568 Goat anti-mouse IgG (H+L) | Thermo Fisher Sci | A-11031 | Mouse | 1:250 | IHC |
| Alexa Fluor 488 Goat anti-rabbit IgG (H+L) | Thermo Fisher Sci | A-11008 | Rabbit | 1:250 | IHC |
| Alexa Fluor 488 Goat anti-mouse IgG (H+L) | Thermo Fisher Sci | A-11029 | Mouse | 1:250 | IHC |
| Alexa Fluor 488 Goat anti-chicken IgY (H+L) | Thermo Fisher Sci | A-11039 | Chicken | 1:250 | IHC |
| Alexa Fluor 488 Goat anti-chicken IgG (H+L) | Thermo Fisher Sci | A-11040 | Chicken | 1:250 | IHC |
| Goat anti-rabbit IgG (H+L)-HRP | Cell Signaling | 7074 | Rabbit | 1:3000 | WB |
| Goat anti-mouse IgG (H+L)-HRP | Cell Signaling | 7076 | Mouse | 1:3000 | WB |

**Supplemental Table 2A – aPAT % Time in Target zone across treatment and genotype.**

| Three-way ANVOA analysis (Males) | | | | | |
|---|---|---|---|---|---|
| **Source of Variation** | **% of total variation** | **P value** | **P value summary** | **Significant?** | **Geisser-Greenhouse's epsilon** |
| Trial | 24.61 | <0.0001 | **** | Yes | 0.6735 |
| Treatment | 4.115 | 0.0078 | ** | Yes | |
| Genotype | 5.737 | 0.0019 | ** | Yes | |

| | | | | | |
|---|---|---|---|---|---|
| Trial x Treatment | 0.4388 | 0.5174 | ns | No | |
| Trial x Genotype | 3.411 | <0.0001 | **** | Yes | |
| Treatment x Genotype | 1.424 | 0.1097 | ns | No | |
| Trial x Treatment x Genotype | 0.8466 | 0.1511 | ns | No | |
| Subject | 28.52 | | | | |
| **ANOVA table** | **SS** | **DF** | **MS** | **F (DFn, DFd)** | **P value** |
| Trial | 1422 | 5 | 284.4 | F (3.367, 178.5) = 47.52 | P<0.0001 |
| Treatment | 237.7 | 1 | 237.7 | F (1, 53) = 7.646 | P=0.0078 |
| Genotype | 331.5 | 1 | 331.5 | F (1, 53) = 10.66 | P=0.0019 |
| Trial x Treatment | 25.35 | 5 | 5.070 | F (5, 265) = 0.8471 | P=0.5174 |
| Trial x Genotype | 197.1 | 5 | 39.42 | F (5, 265) = 6.586 | P<0.0001 |
| Treatment x Genotype | 82.27 | 1 | 82.27 | F (1, 53) = 2.646 | P=0.1097 |
| Trial x Treatment x Genotype | 48.91 | 5 | 9.783 | F (5, 265) = 1.635 | P=0.1511 |
| Subject | 1648 | 53 | 31.09 | | |

### Supplemental Table 2B: aPAT % Time in Target zone across treatment or genotype.

| Two-way ANOVA | WTNT vs. TGNT (Males) | | | | |
|---|---|---|---|---|---|
| **Source of Variation** | **% of total variation** | **P value** | **P value summary** | **Significant?** | **Geisser-Greenhouse's epsilon** |
| Trial x Genotype | 5.529 | <0.0001 | **** | Yes | |
| Trial | 22.47 | <0.0001 | **** | Yes | 0.5902 |
| Genotype | 10.27 | 0.0045 | ** | Yes | |
| Subject | 31.37 | <0.0001 | **** | Yes | |
| **ANOVA table** | **SS** | **DF** | **MS** | **F (DFn, DFd)** | **P value** |
| Trial x Genotype | 218.5 | 5 | 43.70 | F (5, 145) = 5.682 | P<0.0001 |
| Trial | 887.8 | 5 | 177.6 | F (2.951, 85.59) = 23.09 | P<0.0001 |
| Genotype | 405.7 | 1 | 405.7 | F (1, 29) = 9.492 | P=0.0045 |
| Subject | 1240 | 29 | 42.74 | F (29, 145) = 5.558 | P<0.0001 |

| Two-way ANOVA | TGNT vs. TGTR (Males) | | | | |
|---|---|---|---|---|---|
| **Source of Variation** | **% of total variation** | **P value** | **P value summary** | **Significant?** | **Geisser-Greenhouse's epsilon** |
| Trial x Treatment | 1.674 | 0.1356 | ns | No | |
| Trial | 30.41 | <0.0001 | **** | Yes | 0.5865 |
| Treatment | 7.132 | 0.0181 | * | Yes | |
| Subject | 31.65 | <0.0001 | **** | Yes | |
| **ANOVA table** | **SS** | **DF** | **MS** | **F (DFn, DFd)** | **P value** |
| Trial x Treatment | 73.67 | 5 | 14.73 | F (5, 140) = 1.713 | P=0.1356 |
| Trial | 1338 | 5 | 267.5 | F (2.933, 82.11) = 31.10 | P<0.0001 |
| Treatment | 313.8 | 1 | 313.8 | F (1, 28) = 6.308 | P=0.0181 |

| | | | | | |
|---|---|---|---|---|---|
| Subject | 1393 | 28 | 49.74 | F (28, 140) = 5.781 | P<0.0001 |

| Two-way ANOVA | WTNT vs. WTTR (Males) | | | | |
|---|---|---|---|---|---|
| **Source of Variation** | **% of total variation** | **P value** | **P value summary** | **Significant?** | **Geisser-Greenhouse's epsilon** |
| Trial x Treatment | 0.3756 | 0.9428 | ns | No | |
| Trial | 33.11 | <0.0001 | **** | Yes | 0.5445 |
| Treatment | 1.958 | 0.1813 | ns | No | |
| Subject | 25.89 | <0.0001 | **** | Yes | |
| **ANOVA table** | **SS** | **DF** | **MS** | **F (DFn, DFd)** | **P value** |
| Trial x Treatment | 3.700 | 5 | 0.7401 | F (5, 125) = 0.2425 | P=0.9428 |
| Trial | 326.3 | 5 | 65.25 | F (2.722, 68.06) = 21.38 | P<0.0001 |
| Treatment | 19.29 | 1 | 19.29 | F (1, 25) = 1.891 | P=0.1813 |
| Subject | 255.1 | 25 | 10.20 | F (25, 125) = 3.343 | P<0.0001 |

| **Two-way ANOVA** | **WTTR vs. TGTR (Males)** | | | | |
|---|---|---|---|---|---|
| **Source of Variation** | **% of total variation** | **P value** | **P value summary** | **Significant?** | **Geisser-Greenhouse's epsilon** |
| Trial x Genotype | 2.729 | 0.0643 | ns | No | |
| Trial | 37.86 | <0.0001 | **** | Yes | 0.5779 |
| Genotype | 2.495 | 0.1453 | ns | No | |
| Subject | 26.43 | <0.0001 | **** | Yes | |
| **ANOVA table** | **SS** | **DF** | **MS** | **F (DFn, DFd)** | **P value** |
| Trial x Genotype | 42.15 | 5 | 8.430 | F (5, 120) = 2.148 | P=0.0643 |
| Trial | 584.7 | 5 | 116.9 | F (2.889, 69.34) = 29.80 | P<0.0001 |
| Genotype | 38.53 | 1 | 38.53 | F (1, 24) = 2.266 | P=0.1453 |
| Subject | 408.2 | 24 | 17.01 | F (24, 120) = 4.334 | P<0.0001 |

**Three-way ANVOA analysis (Females)**

| Source of Variation | % of total variation | P value | P value summary | Significant? | Geisser-Greenhouse's epsilon |
|---|---|---|---|---|---|
| Trial | 21.04 | <0.0001 | **** | Yes | 0.3806 |
| Treatment | 0.02661 | 0.7780 | ns | No | |
| Genotype | 0.2990 | 0.3463 | ns | No | |
| Trial x Treatment | 0.04944 | 0.9978 | ns | No | |
| Trial x Genotype | 0.4805 | 0.7252 | ns | No | |
| Treatment x Genotype | 0.02303 | 0.7932 | ns | No | |
| Trial x Treatment x Genotype | 0.05120 | 0.9976 | ns | No | |
| Subject | 20.92 | | | | |

| ANOVA table | SS | DF | MS | F (DFn, DFd) | P value |
|---|---|---|---|---|---|
| Trial | 476.2 | 5 | 95.23 | F (1.903, 119.9) = 24.83 | P<0.0001 |
| Treatment | 0.6023 | 1 | 0.6023 | F (1, 63) = 0.08013 | P=0.7780 |
| Genotype | 6.767 | 1 | 6.767 | F (1, 63) = 0.9003 | P=0.3463 |
| Trial x Treatment | 1.119 | 5 | 0.2238 | F (5, 315) = 0.05835 | P=0.9978 |
| Trial x Genotype | 10.88 | 5 | 2.175 | F (5, 315) = 0.5671 | P=0.7252 |
| Treatment x Genotype | 0.5212 | 1 | 0.5212 | F (1, 63) = 0.06933 | P=0.7932 |
| Trial x Treatment x Genotype | 1.159 | 5 | 0.2318 | F (5, 315) = 0.06043 | P=0.9976 |
| Subject | 473.6 | 63 | 7.517 | | |

| Two-way ANOVA | WTNT vs. TGNT (Females) | | | | |
|---|---|---|---|---|---|
| **Source of Variation** | **% of total variation** | **P value** | **P value summary** | **Significant?** | **Geisser-Greenhouse's epsilon** |
| Trial x Genotype | 0.5095 | 0.8671 | ns | No | |
| Trial | 19.34 | <0.0001 | **** | Yes | 0.3722 |
| Genotype | 0.1367 | 0.6000 | ns | No | |
| Subject | 20.57 | 0.0042 | ** | Yes | |
| **ANOVA table** | **SS** | **DF** | **MS** | **F (DFn, DFd)** | **P value** |
| Trial x Genotype | 9.129 | 5 | 1.826 | F (5, 210) = 0.3726 | P=0.8671 |
| Trial | 346.6 | 5 | 69.32 | F (1.861, 78.15) = 14.15 | P<0.0001 |
| Genotype | 2.450 | 1 | 2.450 | F (1, 42) = 0.2792 | P=0.6000 |
| Subject | 368.5 | 42 | 8.774 | F (42, 210) = 1.791 | P=0.0042 |

| Two-way ANOVA | TGNT vs. TGTR (Females) | | | | |
|---|---|---|---|---|---|
| **Source of Variation** | **% of total variation** | **P value** | **P value summary** | **Significant?** | **Geisser-Greenhouse's epsilon** |
| Trial x Treatment | 0.2576 | 0.9776 | ns | No | |
| Trial | 40.92 | <0.0001 | **** | Yes | 0.3094 |
| Treatment | 0.1957 | 0.5851 | ns | No | |
| Subject | 15.99 | 0.0090 | ** | Yes | |
| **ANOVA table** | **SS** | **DF** | **MS** | **F (DFn, DFd)** | **P value** |
| Trial x Treatment | 1.312 | 5 | 0.2624 | F (5, 125) = 0.1567 | P=0.9776 |
| Trial | 208.4 | 5 | 41.67 | F (1.547, 38.68) = 24.89 | P<0.0001 |
| Treatment | 0.9964 | 1 | 0.9964 | F (1, 25) = 0.3059 | P=0.5851 |
| Subject | 81.43 | 25 | 3.257 | F (25, 125) = 1.946 | P=0.0090 |

| Two-way ANOVA | WTNT vs. WTTR (Females) | | | | |
|---|---|---|---|---|---|
| **Source of Variation** | **% of total variation** | **P value** | **P value summary** | **Significant?** | **Geisser-Greenhouse's epsilon** |
| Trial x Treatment | 0.05242 | 0.9994 | ns | No | |
| Trial | 16.53 | 0.0001 | *** | Yes | 0.3629 |
| Treatment | 9.615e-005 | 0.9899 | ns | No | |
| Subject | 22.44 | 0.0017 | ** | Yes | |
| **ANOVA table** | **SS** | **DF** | **MS** | **F (DFn, DFd)** | **P value** |
| Trial x Treatment | 0.9160 | 5 | 0.1832 | F (5, 190) = 0.03485 | P=0.9994 |
| Trial | 288.8 | 5 | 57.77 | F (1.814, 68.95) = 10.99 | P=0.0001 |
| Treatment | 0.001680 | 1 | 0.001680 | F (1, 38) = 0.0001628 | P=0.9899 |
| Subject | 392.1 | 38 | 10.32 | F (38, 190) = 1.963 | P=0.0017 |

| Two-way ANOVA | WTTR vs. TGTR (Females) | | | | |
|---|---|---|---|---|---|
| **Source of Variation** | **% of total variation** | **P value** | **P value summary** | **Significant?** | **Geisser-Greenhouse's epsilon** |
| Trial x Genotype | 0.9054 | 0.7757 | ns | No | |
| Trial | 37.76 | <0.0001 | **** | Yes | 0.3957 |
| Genotype | 0.9170 | 0.3634 | ns | No | |

| | | | | | |
|---|---|---|---|---|---|
| Subject | 22.31 | 0.0002 | *** | Yes | |
| **ANOVA table** | **SS** | **DF** | **MS** | **F (DFn, DFd)** | **P value** |
| Trial x Genotype | 4.263 | 5 | 0.8526 | F (5, 105) = 0.5000 | P=0.7757 |
| Trial | 177.8 | 5 | 35.56 | F (1.979, 41.55) = 20.85 | P<0.0001 |
| Genotype | 4.317 | 1 | 4.317 | F (1, 21) = 0.8630 | P=0.3634 |
| Subject | 105.1 | 21 | 5.003 | F (21, 105) = 2.934 | P=0.0002 |

**Supplemental Table 2: BT-11 mitigates spatial learning deficits in male TgF344 rats.** The % time spent in the target zone during an active place avoidance task across 6 trials was determined for each experimental group (WTNT, TGNT, WTTR, and TGTR) for both males and females. 3-way ANOVA analysis is used to assess effects of drug treatment, genotype, and trial (A). Individual 2-way ANOVAs were performed to assess genotype or treatment differences across trials between 2 experimental groups (WTNT vs. TGNT, TGNT vs. TGTR, WTNT vs. WTTR, and WTTR vs. TGTR) for both males and females (B). Abbreviations: WTNT= wild type not treated, TGNT= transgenic not treated, WTTR= wild type BT-11 treated, TGTR= transgenic BT-11 treated.

#### Supplemental Table 3 – Unpaired *t* test Aβ % area

| **Males** | TGNT vs. TGTR |
|---|---|
| **Unpaired t test** | |
| P value | 0.0338 |
| P value summary | * |
| Significantly different (P < 0.05)? | Yes |
| One- or two-tailed P value? | One-tailed |
| t, df | t=1.981, df=14 |

| **Females** | TGNT vs. TGTR |
|---|---|
| **Unpaired t test** | |
| P value | 0.2008 |
| P value summary | ns |
| Significantly different (P < 0.05)? | No |
| One- or two-tailed P value? | One-tailed |
| t, df | t=0.8633, df=15 |

**Supplemental Table 3 --** Amyloid Beta (antibody Aβ 4G8) levels across the hippocampus in Tg-AD rats. Values represent percent positive signal (out of 100%) as explained in materials and methods. Data was analyzed using unpaired t-tests assessing the dorsal hippocampus. Abbreviations TGNT – transgenic not treated, males *n* = 8, females n=9; TGTR – transgenic BT-11 treated males *n* = 8, females n=8.

#### Supplemental Table 4A – Total Microglia across whole Dorsal hippocampus across genotype and treatment.

| **Two-way ANOVA analysis** | **Whole hippocampus- total Microglia** | **Males** | | |
|---|---|---|---|---|
| **Source of Variation** | **% of total variation** | **P value** | **P value summary** | **Significant?** |
| Interaction | 3.183 | 0.3471 | ns | No |
| Genotype | 3.043 | 0.3577 | ns | No |

| | | | | | |
|---|---|---|---|---|---|
| Treatment | 0.2490 | 0.7910 | ns | No | |
| **ANOVA table** | **SS (Type III)** | **DF** | **MS** | **F (DFn, DFd)** | **P value** |
| Interaction | 6.417 | 1 | 6.417 | F (1, 27) = 0.9158 | P=0.3471 |
| Genotype | 6.135 | 1 | 6.135 | F (1, 27) = 0.8756 | P=0.3577 |
| Treatment | 0.5019 | 1 | 0.5019 | F (1, 27) = 0.07163 | P=0.7910 |

| **Two-way ANOVA analysis** | **Whole hippocampus- total Microglia** | **Females** | | | |
|---|---|---|---|---|---|
| **Source of Variation** | **% of total variation** | **P value** | **P value summary** | **Significant?** | |
| Interaction | 12.62 | 0.0292 | * | Yes | |
| Genotype | 10.91 | 0.0414 | * | Yes | |
| Treatment | 9.877 | 0.0515 | ns | No | |
| **ANOVA table** | **SS (Type III)** | **DF** | **MS** | **F (DFn, DFd)** | **P value** |
| Interaction | 8.138 | 1 | 8.138 | F (1, 26) = 5.326 | P=0.0292 |
| Genotype | 7.033 | 1 | 7.033 | F (1, 26) = 4.603 | P=0.0414 |
| Treatment | 6.369 | 1 | 6.369 | F (1, 26) = 4.168 | P=0.0515 |

### Supplemental Table 4B – Ramified Microglia across whole Dorsal hippocampus across genotype and treatment.

| **Two-way ANOVA analysis** | **Whole hippocampus-ramified Microglia** | **Males** | | | |
|---|---|---|---|---|---|
| **Source of Variation** | **% of total variation** | **P value** | **P value summary** | **Significant?** | |
| Interaction | 2.641 | 0.3891 | ns | No | |
| Genotype | 3.278 | 0.3380 | ns | No | |
| Treatment | 1.518 | 0.5124 | ns | No | |
| **ANOVA table** | **SS (Type III)** | **DF** | **MS** | **F (DFn, DFd)** | **P value** |
| Interaction | 5.255 | 1 | 5.255 | F (1, 27) = 0.7664 | P=0.3891 |
| Genotype | 6.523 | 1 | 6.523 | F (1, 27) = 0.9513 | P=0.3380 |
| Treatment | 3.022 | 1 | 3.022 | F (1, 27) = 0.4407 | P=0.5124 |

| **Two-way ANOVA analysis** | **Whole hippocampus- ramified Microglia** | **Females** | | | |
|---|---|---|---|---|---|
| Source of Variation | % of total variation | P value | P value summary | Significant? | |
| Interaction | 11.93 | 0.0345 | * | Yes | |
| Genotype | 7.703 | 0.0847 | ns | No | |
| Treatment | 12.95 | 0.0282 | * | Yes | |
| **ANOVA table** | **SS (Type III)** | **DF** | **MS** | **F (DFn, DFd)** | **P value** |
| Interaction | 4.987 | 1 | 4.987 | F (1, 26) = 4.976 | P=0.0345 |
| Genotype | 3.220 | 1 | 3.220 | F (1, 26) = 3.212 | P=0.0847 |
| Treatment | 5.415 | 1 | 5.415 | F (1, 26) = 5.402 | P=0.0282 |

### Supplemental Table 4C – Reactive Microglia across whole Dorsal hippocampus across genotype and treatment.

| **Two-way ANOVA analysis** | **Whole hippocampus-reactive Microglia** | **Males** |
|---|---|---|

| **Source of Variation** | **% of total variation** | **P value** | **P value summary** | **Significant?** | |
|---|---|---|---|---|---|
| Interaction | 0.5996 | 0.6298 | ns | No | |
| Genotype | 31.45 | 0.0015 | ** | Yes | |
| Treatment | 0.2721 | 0.7451 | ns | No | |
| **ANOVA table** | **SS (Type III)** | **DF** | **MS** | **F (DFn, DFd)** | **P value** |
| Interaction | 0.009179 | 1 | 0.009179 | F (1, 27) = 0.2377 | P=0.6298 |
| Genotype | 0.4814 | 1 | 0.4814 | F (1, 27) = 12.47 | P=0.0015 |
| Treatment | 0.004166 | 1 | 0.004166 | F (1, 27) = 0.1079 | P=0.7451 |

| **Two-way ANOVA analysis** | **Whole hippocampus-reactive Microglia** | **Females** | | | |
|---|---|---|---|---|---|
| Source of Variation | % of total variation | P value | P value summary | Significant? | |
| Interaction | 9.936 | 0.0611 | ns | No | |
| Genotype | 17.74 | 0.0146 | * | Yes | |
| Treatment | 1.700 | 0.4255 | ns | No | |
| ANOVA table | SS (Type III) | DF | MS | F (DFn, DFd) | P value |
| Interaction | 0.2032 | 1 | 0.2032 | F (1, 26) = 3.831 | P=0.0611 |
| Genotype | 0.3628 | 1 | 0.3628 | F (1, 26) = 6.840 | P=0.0146 |
| Treatment | 0.03477 | 1 | 0.03477 | F (1, 26) = 0.6554 | P=0.4255 |

### Supplemental Table 4D – Amoeboid Microglia across whole Dorsal hippocampus across genotype and treatment.

| **Two-way ANOVA analysis** | **Whole hippocampus-amoeboid Microglia** | **Males** | | | |
|---|---|---|---|---|---|
| **Source of Variation** | **% of total variation** | **P value** | **P value summary** | **Significant?** | |
| Interaction | 1.146 | 0.5239 | ns | No | |
| Genotype | 26.58 | 0.0045 | ** | Yes | |
| Treatment | 1.649 | 0.4453 | ns | No | |
| **ANOVA table** | **SS (Type III)** | **DF** | **MS** | **F (DFn, DFd)** | **P value** |
| Interaction | 0.002034 | 1 | 0.002034 | F (1, 26) = 0.4174 | P=0.5239 |
| Genotype | 0.04718 | 1 | 0.04718 | F (1, 26) = 9.682 | P=0.0045 |
| Treatment | 0.002927 | 1 | 0.002927 | F (1, 26) = 0.6007 | P=0.4453 |

| **Two-way ANOVA analysis** | **Whole hippocampus-amoeboid Microglia** | **Females** | | | |
|---|---|---|---|---|---|
| Source of Variation | % of total variation | P value | P value summary | Significant? | |
| Interaction | 5.400 | 0.1939 | ns | No | |
| Genotype | 14.16 | 0.0402 | * | Yes | |
| Treatment | 0.02751 | 0.9249 | ns | No | |
| ANOVA table | SS (Type III) | DF | MS | F (DFn, DFd) | P value |
| Interaction | 0.01527 | 1 | 0.01527 | F (1, 26) = 1.779 | P=0.1939 |
| Genotype | 0.04005 | 1 | 0.04005 | F (1, 26) = 4.665 | P=0.0402 |

| | | | | | |
|---|---|---|---|---|---|
| Treatment | 7.779e-005 | 1 | 7.779e-005 | F (1, 26) = 0.009062 | P=0.9249 |

**Supplemental Table 4E – Total Microglia across CA1 region across genotype and treatment.**

| **Two-way ANOVA analysis** | **CA1- total Microglia** | **Males** | | | |
|---|---|---|---|---|---|
| **Source of Variation** | **% of total variation** | **P value** | **P value summary** | **Significant?** | |
| Interaction | 0.8416 | 0.6335 | ns | No | |
| Genotype | 1.189 | 0.5713 | ns | No | |
| Treatment | 0.4120 | 0.7384 | ns | No | |
| **ANOVA table** | **SS (Type III)** | **DF** | **MS** | **F (DFn, DFd)** | **P value** |
| Interaction | 1.902 | 1 | 1.902 | F (1, 27) = 0.2326 | P=0.6335 |
| Genotype | 2.686 | 1 | 2.686 | F (1, 27) = 0.3285 | P=0.5713 |
| Treatment | 0.9311 | 1 | 0.9311 | F (1, 27) = 0.1139 | P=0.7384 |

| **Two-way ANOVA analysis** | **CA1- total Microglia** | **Females** | | | |
|---|---|---|---|---|---|
| Source of Variation | % of total variation | P value | P value summary | Significant? | |
| Interaction | 12.39 | 0.0454 | * | Yes | |
| Genotype | 2.729 | 0.3329 | ns | No | |
| Treatment | 8.104 | 0.1010 | ns | No | |
| ANOVA table | SS (Type III) | DF | MS | F (DFn, DFd) | P value |
| Interaction | 8.633 | 1 | 8.633 | F (1, 26) = 4.419 | P=0.0454 |
| Genotype | 1.902 | 1 | 1.902 | F (1, 26) = 0.9736 | P=0.3329 |
| Treatment | 5.647 | 1 | 5.647 | F (1, 26) = 2.891 | P=0.1010 |

**Supplemental Table 4F – Ramified Microglia across CA1 region across genotype and treatment.**

| **Two-way ANOVA analysis** | **CA1- ramified Microglia** | **Males** | | | |
|---|---|---|---|---|---|
| **Source of Variation** | **% of total variation** | **P value** | **P value summary** | **Significant?** | |
| Interaction | 0.05908 | 0.9039 | ns | No | |
| Genotype | 0.3701 | 0.7626 | ns | No | |
| Treatment | 0.3631 | 0.7648 | ns | No | |
| **ANOVA table** | **SS (Type III)** | **DF** | **MS** | **F (DFn, DFd)** | **P value** |
| Interaction | 0.1237 | 1 | 0.1237 | F (1, 25) = 0.01489 | P=0.9039 |
| Genotype | 0.7751 | 1 | 0.7751 | F (1, 25) = 0.09328 | P=0.7626 |
| Treatment | 0.7603 | 1 | 0.7603 | F (1, 25) = 0.09150 | P=0.7648 |

| **Two-way ANOVA analysis** | **CA1- ramified Microglia** | **Females** | | | |
|---|---|---|---|---|---|
| **Source of Variation** | **% of total variation** | **P value** | **P value summary** | **Significant?** | |
| Interaction | 14.91 | 0.0225 | * | Yes | |
| Genotype | 0.4931 | 0.6627 | ns | No | |
| Treatment | 13.90 | 0.0271 | * | Yes | |
| **ANOVA table** | **SS (Type III)** | **DF** | **MS** | **F (DFn, DFd)** | **P value** |
| Interaction | 6.981 | 1 | 6.981 | F (1, 26) = 5.889 | P=0.0225 |

| | | | | | |
|---|---|---|---|---|---|
| Genotype | 0.2308 | 1 | 0.2308 | F (1, 26) = 0.1947 | P=0.6627 |
| Treatment | 6.506 | 1 | 6.506 | F (1, 26) = 5.489 | P=0.0271 |

### Supplemental Table 4G – Reactive Microglia across CA1 region across genotype and treatment.

| **Two-way ANOVA analysis** | **CA1- reactive Microglia** | **Males** | | | |
|---|---|---|---|---|---|
| **Source of Variation** | **% of total variation** | **P value** | **P value summary** | **Significant?** | |
| Interaction | 0.2941 | 0.7630 | ns | No | |
| Genotype | 17.66 | 0.0263 | * | Yes | |
| Treatment | 2.627 | 0.3710 | ns | No | |
| **ANOVA table** | **SS (Type III)** | **DF** | **MS** | **F (DFn, DFd)** | **P value** |
| Interaction | 0.002975 | 1 | 0.002975 | F (1, 25) = 0.09291 | P=0.7630 |
| Genotype | 0.1786 | 1 | 0.1786 | F (1, 25) = 5.577 | P=0.0263 |
| Treatment | 0.02657 | 1 | 0.02657 | F (1, 25) = 0.8298 | P=0.3710 |

| **Two-way ANOVA analysis** | **CA1- reactive Microglia** | **Females** | | | |
|---|---|---|---|---|---|
| **Source of Variation** | **% of total variation** | **P value** | **P value summary** | **Significant?** | |
| Interaction | 1.249 | 0.5159 | ns | No | |
| Genotype | 25.41 | 0.0064 | ** | Yes | |
| Treatment | 0.6996 | 0.6261 | ns | No | |
| **ANOVA table** | **SS (Type III)** | **DF** | **MS** | **F (DFn, DFd)** | **P value** |
| Interaction | 0.01847 | 1 | 0.01847 | F (1, 25) = 0.4344 | P=0.5159 |
| Genotype | 0.3758 | 1 | 0.3758 | F (1, 25) = 8.839 | P=0.0064 |
| Treatment | 0.01035 | 1 | 0.01035 | F (1, 25) = 0.2434 | P=0.6261 |

### Supplemental Table 4H – Amoeboid Microglia across CA1 region across genotype and treatment.

| **Two-way ANOVA analysis** | **CA1- amoeboid Microglia** | **Males** | | | |
|---|---|---|---|---|---|
| **Source of Variation** | **% of total variation** | **P value** | **P value summary** | **Significant?** | |
| Interaction | 9.261 | 0.0945 | ns | No | |
| Genotype | 8.211 | 0.1143 | ns | No | |
| Treatment | 6.449 | 0.1594 | ns | No | |
| **ANOVA table** | **SS (Type III)** | **DF** | **MS** | **F (DFn, DFd)** | **P value** |
| Interaction | 0.009721 | 1 | 0.009721 | F (1, 25) = 3.021 | P=0.0945 |
| Genotype | 0.008618 | 1 | 0.008618 | F (1, 25) = 2.678 | P=0.1143 |
| Treatment | 0.006769 | 1 | 0.006769 | F (1, 25) = 2.104 | P=0.1594 |

| **Two-way ANOVA analysis** | **CA1- amoeboid Microglia** | **Females** | | | |
|---|---|---|---|---|---|
| **Source of Variation** | **% of total variation** | **P value** | **P value summary** | **Significant?** | |
| Interaction | 0.2614 | 0.7875 | ns | No | |
| Genotype | 5.691 | 0.2149 | ns | No | |
| Treatment | 2.373 | 0.4192 | ns | No | |
| **ANOVA table** | **SS (Type III)** | **DF** | **MS** | **F (DFn, DFd)** | **P value** |
| Interaction | 0.002213 | 1 | 0.002213 | F (1, 26) = 0.07420 | P=0.7875 |
| Genotype | 0.04820 | 1 | 0.04820 | F (1, 26) = 1.616 | P=0.2149 |

| | | | | | |
|---|---|---|---|---|---|
| Treatment | 0.02010 | 1 | 0.02010 | F (1, 26) = 0.6738 | P=0.4192 |

### Supplemental Table 4I – Total Microglia across CA3 region across genotype and treatment.

| **Two-way ANOVA analysis** | **CA3- total Microglia** | **Males** | | | |
|---|---|---|---|---|---|
| Source of Variation | % of total variation | P value | P value summary | Significant? | |
| Interaction | 0.3981 | 0.7374 | ns | No | |
| Genotype | 5.537 | 0.2173 | ns | No | |
| Treatment | 0.6221 | 0.6753 | ns | No | |
| ANOVA table | SS (Type III) | DF | MS | F (DFn, DFd) | P value |
| Interaction | 0.8769 | 1 | 0.8769 | F (1, 27) = 0.1147 | P=0.7374 |
| Genotype | 12.19 | 1 | 12.19 | F (1, 27) = 1.596 | P=0.2173 |
| Treatment | 1.370 | 1 | 1.370 | F (1, 27) = 0.1793 | P=0.6753 |

| **Two-way ANOVA analysis** | **CA3- total Microglia** | **Females** | | | |
|---|---|---|---|---|---|
| **Source of Variation** | **% of total variation** | **P value** | **P value summary** | **Significant?** | |
| Interaction | 7.931 | 0.0903 | ns | No | |
| Genotype | 9.978 | 0.0592 | ns | No | |
| Treatment | 11.41 | 0.0446 | * | Yes | |
| **ANOVA table** | **SS (Type III)** | **DF** | **MS** | **F (DFn, DFd)** | **P value** |
| Interaction | 4.973 | 1 | 4.973 | F (1, 26) = 3.095 | P=0.0903 |
| Genotype | 6.257 | 1 | 6.257 | F (1, 26) = 3.894 | P=0.0592 |
| Treatment | 7.158 | 1 | 7.158 | F (1, 26) = 4.455 | P=0.0446 |

### Supplemental Table 4J – Ramified Microglia across CA3 region across genotype and treatment.

| **Two-way ANOVA analysis** | **CA3- ramified Microglia** | **Males** | | | |
|---|---|---|---|---|---|
| **Source of Variation** | **% of total variation** | **P value** | **P value summary** | **Significant?** | |
| Interaction | 0.1478 | 0.8411 | ns | No | |
| Genotype | 5.522 | 0.2268 | ns | No | |
| Treatment | 0.7464 | 0.6528 | ns | No | |
| **ANOVA table** | **SS (Type III)** | **DF** | **MS** | **F (DFn, DFd)** | **P value** |
| Interaction | 0.3312 | 1 | 0.3312 | F (1, 26) = 0.04103 | P=0.8411 |
| Genotype | 12.37 | 1 | 12.37 | F (1, 26) = 1.533 | P=0.2268 |
| Treatment | 1.672 | 1 | 1.672 | F (1, 26) = 0.2072 | P=0.6528 |

| **Two-way ANOVA analysis** | **CA3- ramified Microglia** | **Females** | | | |
|---|---|---|---|---|---|
| **Source of Variation** | **% of total variation** | **P value** | **P value summary** | **Significant?** | |
| Interaction | 6.521 | 0.1131 | ns | No | |
| Genotype | 13.79 | 0.0247 | * | Yes | |
| Treatment | 12.65 | 0.0308 | * | Yes | |
| **ANOVA table** | **SS (Type III)** | **DF** | **MS** | **F (DFn, DFd)** | **P value** |
| Interaction | 4.342 | 1 | 4.342 | F (1, 26) = 2.689 | P=0.1131 |

| | | | | | |
|---|---|---|---|---|---|
| Genotype | 9.184 | 1 | 9.184 | F (1, 26) = 5.688 | P=0.0247 |
| Treatment | 8.422 | 1 | 8.422 | F (1, 26) = 5.216 | P=0.0308 |

#### Supplemental Table 4K – Reactive Microglia across CA3 region across genotype and treatment.

| **Two-way ANOVA analysis** | **CA3- reactive Microglia** | **Males** | | | |
|---|---|---|---|---|---|
| **Source of Variation** | **% of total variation** | **P value** | **P value summary** | **Significant?** | |
| Interaction | 3.149 | 0.3247 | ns | No | |
| Genotype | 12.10 | 0.0598 | ns | No | |
| Treatment | 4.271 | 0.2530 | ns | No | |
| **ANOVA table** | **SS (Type III)** | **DF** | **MS** | **F (DFn, DFd)** | **P value** |
| Interaction | 0.02903 | 1 | 0.02903 | F (1, 26) = 1.008 | P=0.3247 |
| Genotype | 0.1115 | 1 | 0.1115 | F (1, 26) = 3.873 | P=0.0598 |
| Treatment | 0.03936 | 1 | 0.03936 | F (1, 26) = 1.367 | P=0.2530 |

| **Two-way ANOVA analysis** | **CA3- reactive Microglia** | **Females** | | | |
|---|---|---|---|---|---|
| **Source of Variation** | **% of total variation** | **P value** | **P value summary** | **Significant?** | |
| Interaction | 9.125 | 0.0833 | ns | No | |
| Genotype | 12.18 | 0.0475 | * | Yes | |
| Treatment | 2.598 | 0.3453 | ns | No | |
| **ANOVA table** | **SS (Type III)** | **DF** | **MS** | **F (DFn, DFd)** | **P value** |
| Interaction | 0.2332 | 1 | 0.2332 | F (1, 26) = 3.245 | P=0.0833 |
| Genotype | 0.3112 | 1 | 0.3112 | F (1, 26) = 4.329 | P=0.0475 |
| Treatment | 0.06642 | 1 | 0.06642 | F (1, 26) = 0.9239 | P=0.3453 |

#### Supplemental Table 4L – Amoeboid Microglia across CA3 region across genotype and treatment.

| **Two-way ANOVA analysis** | **CA3- amoeboid Microglia** | **Males** | | | |
|---|---|---|---|---|---|
| **Source of Variation** | **% of total variation** | **P value** | **P value summary** | **Significant?** | |
| Interaction | 0.0002265 | 0.9935 | ns | No | |
| Genotype | 16.17 | 0.0370 | * | Yes | |
| Treatment | 0.4215 | 0.7249 | ns | No | |
| **ANOVA table** | **SS (Type III)** | **DF** | **MS** | **F (DFn, DFd)** | **P value** |
| Interaction | 1.296e-007 | 1 | 1.296e-007 | F (1, 25) = 6.807e-005 | P=0.9935 |
| Genotype | 0.009249 | 1 | 0.009249 | F (1, 25) = 4.858 | P=0.0370 |
| Treatment | 0.0002411 | 1 | 0.0002411 | F (1, 25) = 0.1267 | P=0.7249 |

| **Two-way ANOVA analysis** | **CA3- amoeboid Microglia** | **Females** | | |
|---|---|---|---|---|
| **Source of Variation** | **% of total variation** | **P value** | **P value summary** | **Significant?** |
| Interaction | 6.410 | 0.1663 | ns | No |
| Genotype | 9.878 | 0.0888 | ns | No |
| Treatment | 0.07189 | 0.8813 | ns | No |

| ANOVA table | SS (Type III) | DF | MS | F (DFn, DFd) | P value |
|---|---|---|---|---|---|
| Interaction | 0.009587 | 1 | 0.009587 | F (1, 26) = 2.028 | P=0.1663 |
| Genotype | 0.01477 | 1 | 0.01477 | F (1, 26) = 3.125 | P=0.0888 |
| Treatment | 0.0001075 | 1 | 0.0001075 | F (1, 26) = 0.02274 | P=0.8813 |

### Supplemental Table 4M – Total Microglia across DG region across genotype and treatment.

| Two-way ANOVA analysis | DG- total Microglia | Males | | | |
|---|---|---|---|---|---|
| **Source of Variation** | **% of total variation** | **P value** | **P value summary** | **Significant?** | |
| Interaction | 1.979 | 0.4497 | ns | No | |
| Genotype | 6.500 | 0.1759 | ns | No | |
| Treatment | 1.197 | 0.5558 | ns | No | |
| **ANOVA table** | **SS (Type III)** | **DF** | **MS** | **F (DFn, DFd)** | **P value** |
| Interaction | 4.514 | 1 | 4.514 | F (1, 27) = 0.5883 | P=0.4497 |
| Genotype | 14.82 | 1 | 14.82 | F (1, 27) = 1.932 | P=0.1759 |
| Treatment | 2.729 | 1 | 2.729 | F (1, 27) = 0.3557 | P=0.5558 |

| Two-way ANOVA analysis | DG- total Microglia | Females | | | |
|---|---|---|---|---|---|
| **Source of Variation** | **% of total variation** | **P value** | **P value summary** | **Significant?** | |
| Interaction | 12.45 | 0.0229 | * | Yes | |
| Genotype | 19.85 | 0.0051 | ** | Yes | |
| Treatment | 7.362 | 0.0741 | ns | No | |
| **ANOVA table** | **SS (Type III)** | **DF** | **MS** | **F (DFn, DFd)** | **P value** |
| Interaction | 9.049 | 1 | 9.049 | F (1, 26) = 5.852 | P=0.0229 |
| Genotype | 14.43 | 1 | 14.43 | F (1, 26) = 9.333 | P=0.0051 |
| Treatment | 5.353 | 1 | 5.353 | F (1, 26) = 3.462 | P=0.0741 |

### Supplemental Table 4N – Ramified Microglia across DG region across genotype and treatment.

| Two-way ANOVA analysis | DG- ramified Microglia | Males | | | |
|---|---|---|---|---|---|
| **Source of Variation** | **% of total variation** | **P value** | **P value summary** | **Significant?** | |
| Interaction | 1.067 | 0.5848 | ns | No | |
| Genotype | 6.702 | 0.1774 | ns | No | |
| Treatment | 1.891 | 0.4681 | ns | No | |
| **ANOVA table** | **SS (Type III)** | **DF** | **MS** | **F (DFn, DFd)** | **P value** |
| Interaction | 2.486 | 1 | 2.486 | F (1, 26) = 0.3061 | P=0.5848 |
| Genotype | 15.61 | 1 | 15.61 | F (1, 26) = 1.922 | P=0.1774 |
| Treatment | 4.405 | 1 | 4.405 | F (1, 26) = 0.5423 | P=0.4681 |

| Two-way ANOVA analysis | DG- ramified Microglia | Females | | | |
|---|---|---|---|---|---|
| **Source of Variation** | **% of total variation** | **P value** | **P value summary** | **Significant?** | |
| Interaction | 9.172 | 0.0451 | * | Yes | |
| Genotype | 24.56 | 0.0020 | ** | Yes | |
| Treatment | 7.841 | 0.0626 | ns | No | |
| **ANOVA table** | **SS (Type III)** | **DF** | **MS** | **F (DFn, DFd)** | **P value** |
| Interaction | 7.252 | 1 | 7.252 | F (1, 26) = 4.429 | P=0.0451 |
| Genotype | 19.42 | 1 | 19.42 | F (1, 26) = 11.86 | P=0.0020 |
| Treatment | 6.200 | 1 | 6.200 | F (1, 26) = 3.786 | P=0.0626 |

### Supplemental Table 4O – Reactive Microglia across DG region across genotype and treatment.

| Two-way ANOVA analysis | DG- reactive Microglia | Males | | | |
|---|---|---|---|---|---|
| Source of Variation | % of total variation | P value | P value summary | Significant? | |
| Interaction | 6.051 | 0.1404 | ns | No | |
| Genotype | 25.12 | 0.0046 | ** | Yes | |
| Treatment | 2.312 | 0.3559 | ns | No | |
| ANOVA table | SS (Type III) | DF | MS | F (DFn, DFd) | P value |
| Interaction | 0.1227 | 1 | 0.1227 | F (1, 26) = 2.313 | P=0.1404 |
| Genotype | 0.5093 | 1 | 0.5093 | F (1, 26) = 9.603 | P=0.0046 |
| Treatment | 0.04686 | 1 | 0.04686 | F (1, 26) = 0.8835 | P=0.3559 |

| Two-way ANOVA analysis | DG- reactive Microglia | Females | | | |
|---|---|---|---|---|---|
| Source of Variation | % of total variation | P value | P value summary | Significant? | |
| Interaction | 11.31 | 0.0390 | * | Yes | |
| Genotype | 21.56 | 0.0059 | ** | Yes | |
| Treatment | 1.425 | 0.4472 | ns | No | |
| ANOVA table | SS (Type III) | DF | MS | F (DFn, DFd) | P value |
| Interaction | 0.3029 | 1 | 0.3029 | F (1, 26) = 4.727 | P=0.0390 |
| Genotype | 0.5776 | 1 | 0.5776 | F (1, 26) = 9.014 | P=0.0059 |
| Treatment | 0.03816 | 1 | 0.03816 | F (1, 26) = 0.5956 | P=0.4472 |

### Supplemental Table 4P – Amoeboid Microglia across DG region across genotype and treatment.

| Two-way ANOVA analysis | DG- amoeboid Microglia | Males | | | |
|---|---|---|---|---|---|
| Source of Variation | % of total variation | P value | P value summary | Significant? | |
| Interaction | 12.11 | 0.0494 | * | Yes | |
| Genotype | 11.95 | 0.0509 | ns | No | |
| Treatment | 3.296 | 0.2923 | ns | No | |
| ANOVA table | SS (Type III) | DF | MS | F (DFn, DFd) | P value |
| Interaction | 0.03948 | 1 | 0.03948 | F (1, 26) = 4.247 | P=0.0494 |
| Genotype | 0.03894 | 1 | 0.03894 | F (1, 26) = 4.189 | P=0.0509 |
| Treatment | 0.01074 | 1 | 0.01074 | F (1, 26) = 1.156 | P=0.2923 |

| Two-way ANOVA analysis | DG- amoeboid Microglia | Females | | | |
|---|---|---|---|---|---|
| Source of Variation | % of total variation | P value | P value summary | Significant? | |
| Interaction | 8.551 | 0.0842 | ns | No | |
| Genotype | 19.36 | 0.0120 | * | Yes | |
| Treatment | 0.5433 | 0.6546 | ns | No | |
| ANOVA table | SS (Type III) | DF | MS | F (DFn, DFd) | P value |
| Interaction | 0.02818 | 1 | 0.02818 | F (1, 26) = 3.224 | P=0.0842 |
| Genotype | 0.06380 | 1 | 0.06380 | F (1, 26) = 7.300 | P=0.0120 |
| Treatment | 0.001790 | 1 | 0.001790 | F (1, 26) = 0.2049 | P=0.6546 |

### Supplemental Table 4Q – Total Microglia across SB region across genotype and treatment.

| Two-way ANOVA analysis | SB- total Microglia | Males | | |
|---|---|---|---|---|
| Source of Variation | % of total variation | P value | P value summary | Significant? |
| Interaction | 2.186 | 0.4398 | ns | No |
| Genotype | 1.400 | 0.5356 | ns | No |
| Treatment | 0.6563 | 0.6709 | ns | No |

| ANOVA table | SS (Type III) | DF | MS | F (DFn, DFd) | P value |
|---|---|---|---|---|---|
| Interaction | 4.844 | 1 | 4.844 | F (1, 27) = 0.6148 | P=0.4398 |
| Genotype | 3.103 | 1 | 3.103 | F (1, 27) = 0.3938 | P=0.5356 |
| Treatment | 1.454 | 1 | 1.454 | F (1, 27) = 0.1846 | P=0.6709 |

| Two-way ANOVA analysis | SB- total Microglia | Females | | | |
|---|---|---|---|---|---|
| **Source of Variation** | **% of total variation** | **P value** | **P value summary** | **Significant?** | |
| Interaction | 5.400 | 0.1939 | ns | No | |
| Genotype | 14.16 | 0.0402 | * | Yes | |
| Treatment | 0.02751 | 0.9249 | ns | No | |
| **ANOVA table** | **SS (Type III)** | **DF** | **MS** | **F (DFn, DFd)** | **P value** |
| Interaction | 0.01527 | 1 | 0.01527 | F (1, 26) = 1.779 | P=0.1939 |
| Genotype | 0.04005 | 1 | 0.04005 | F (1, 26) = 4.665 | P=0.0402 |
| Treatment | 7.779e-005 | 1 | 7.779e-005 | F (1, 26) = 0.009062 | P=0.9249 |

### Supplemental Table 4R – Ramified Microglia across SB region across genotype and treatment.

| Two-way ANOVA analysis | SB- ramified Microglia | Males | | | |
|---|---|---|---|---|---|
| **Source of Variation** | **% of total variation** | **P value** | **P value summary** | **Significant?** | |
| Interaction | 1.244 | 0.5667 | ns | No | |
| Genotype | 1.816 | 0.4894 | ns | No | |
| Treatment | 1.132 | 0.5844 | ns | No | |
| **ANOVA table** | **SS (Type III)** | **DF** | **MS** | **F (DFn, DFd)** | **P value** |
| Interaction | 2.738 | 1 | 2.738 | F (1, 26) = 0.3368 | P=0.5667 |
| Genotype | 3.998 | 1 | 3.998 | F (1, 26) = 0.4918 | P=0.4894 |
| Treatment | 2.493 | 1 | 2.493 | F (1, 26) = 0.3067 | P=0.5844 |

| Two-way ANOVA analysis | SB- ramified Microglia | Females | | | |
|---|---|---|---|---|---|
| **Source of Variation** | **% of total variation** | **P value** | **P value summary** | **Significant?** | |
| Interaction | 13.75 | 0.0173 | * | Yes | |
| Genotype | 11.52 | 0.0281 | * | Yes | |
| Treatment | 13.56 | 0.0180 | * | Yes | |
| **ANOVA table** | **SS (Type III)** | **DF** | **MS** | **F (DFn, DFd)** | **P value** |
| Interaction | 10.63 | 1 | 10.63 | F (1, 26) = 6.462 | P=0.0173 |
| Genotype | 8.905 | 1 | 8.905 | F (1, 26) = 5.411 | P=0.0281 |
| Treatment | 10.49 | 1 | 10.49 | F (1, 26) = 6.373 | P=0.0180 |

### Supplemental Table 4S – Reactive Microglia across SB region across genotype and treatment.

| Two-way ANOVA analysis | SB- reactive Microglia | Males | | | |
|---|---|---|---|---|---|
| **Source of Variation** | **% of total variation** | **P value** | **P value summary** | **Significant?** | |
| Interaction | 2.061 | 0.4236 | ns | No | |
| Genotype | 11.33 | 0.0678 | ns | No | |
| Treatment | 6.113 | 0.1733 | ns | No | |
| **ANOVA table** | **SS (Type III)** | **DF** | **MS** | **F (DFn, DFd)** | **P value** |
| Interaction | 0.03793 | 1 | 0.03793 | F (1, 26) = 0.6610 | P=0.4236 |
| Genotype | 0.2084 | 1 | 0.2084 | F (1, 26) = 3.633 | P=0.0678 |
| Treatment | 0.1125 | 1 | 0.1125 | F (1, 26) = 1.960 | P=0.1733 |

| Two-way ANOVA analysis | SB- reactive Microglia | Females | | |
|---|---|---|---|---|
| **Source of Variation** | **% of total variation** | **P value** | **P value summary** | **Significant?** |
| Interaction | 13.65 | 0.0217 | * | Yes |

| | | | | | |
|---|---|---|---|---|---|
| Genotype | 15.79 | 0.0143 | * | Yes | |
| Treatment | 6.226 | 0.1111 | ns | No | |
| **ANOVA table** | **SS (Type III)** | **DF** | **MS** | **F (DFn, DFd)** | **P value** |
| Interaction | 0.5778 | 1 | 0.5778 | F (1, 26) = 5.963 | P=0.0217 |
| Genotype | 0.6686 | 1 | 0.6686 | F (1, 26) = 6.900 | P=0.0143 |
| Treatment | 0.2636 | 1 | 0.2636 | F (1, 26) = 2.721 | P=0.1111 |

**Supplemental Table 4T – Amoeboid Microglia across SB region across genotype and treatment.**

| **Two-way ANOVA analysis** | **SB- amoeboid Microglia** | **Males** | | | |
|---|---|---|---|---|---|
| **Source of Variation** | **% of total variation** | **P value** | **P value summary** | **Significant?** | |
| Interaction | 4.726 | 0.2331 | ns | No | |
| Genotype | 2.435 | 0.3889 | ns | No | |
| Treatment | 10.80 | 0.0764 | ns | No | |
| **ANOVA table** | **SS (Type III)** | **DF** | **MS** | **F (DFn, DFd)** | **P value** |
| Interaction | 0.008242 | 1 | 0.008242 | F (1, 26) = 1.490 | P=0.2331 |
| Genotype | 0.004247 | 1 | 0.004247 | F (1, 26) = 0.7678 | P=0.3889 |
| Treatment | 0.01883 | 1 | 0.01883 | F (1, 26) = 3.405 | P=0.0764 |

| **Two-way ANOVA analysis** | **SB- amoeboid Microglia** | **Females** | | | |
|---|---|---|---|---|---|
| **Source of Variation** | **% of total variation** | **P value** | **P value summary** | **Significant?** | |
| Interaction | 7.586 | 0.0974 | ns | No | |
| Genotype | 20.33 | 0.0092 | ** | Yes | |
| Treatment | 2.338 | 0.3486 | ns | No | |
| **ANOVA table** | **SS (Type III)** | **DF** | **MS** | **F (DFn, DFd)** | **P value** |
| Interaction | 0.04568 | 1 | 0.04568 | F (1, 26) = 2.957 | P=0.0974 |
| Genotype | 0.1224 | 1 | 0.1224 | F (1, 26) = 7.924 | P=0.0092 |
| Treatment | 0.01408 | 1 | 0.01408 | F (1, 26) = 0.9112 | P=0.3486 |

**Supplemental Table 4 – Microglia (antibody Iba1) levels across the hippocampus.** Values represent percent positive signal (out of 100%) across the hippocampus and its subregions in across different morphologies (total, ramified, reactive, and amoeboid) for both male and female rats across four experimental groups (WTNT, TGNT. WTTR, and TGTR). Data was analyzed using two- way ANOVAs. Abbreviations: HC= hippocampus; CA= cornu ammonis; DG= dentate gyrus; SB= subiculum; WTNT= wild type not treated, males n=8; females n=7; TGNT= transgenic not treated, males n=8; females n=9); WTTR= wild type BT-11 treated, males n=8; females n=6; TGTR= transgenic BT-11 treated, males n=7; females n=8.

**Supplemental Table 5A – NeuN (count/square nm) across Dorsal HC across genotype and treatment.**

| **Two-way ANOVA analysis** | **Males** | | | | |
|---|---|---|---|---|---|
| **Source of Variation** | **% of total variation** | **P value** | **P value summary** | **Significant?** | |
| Interaction | 0.07044 | 0.8881 | ns | No | |
| Genotype | 3.068 | 0.3572 | ns | No | |
| Treatment | 10.07 | 0.1017 | ns | No | |
| **ANOVA table** | **SS (Type III)** | **DF** | **MS** | **F (DFn, DFd)** | **P value** |
| Interaction | 72.01 | 1 | 72.01 | F (1, 25) = 0.02020 | P=0.8881 |
| Genotype | 3136 | 1 | 3136 | F (1, 25) = 0.8800 | P=0.3572 |
| Treatment | 10289 | 1 | 10289 | F (1, 25) = 2.887 | P=0.1017 |

| Two-way ANOVA analysis | Females | | | | |
|---|---|---|---|---|---|
| **Source of Variation** | **% of total variation** | **P value** | **P value summary** | **Significant?** | |
| Interaction | 0.1401 | 0.8547 | ns | No | |
| Genotype | 3.738 | 0.3485 | ns | No | |
| Treatment | 2.364 | 0.4544 | ns | No | |
| **ANOVA table** | **SS (Type III)** | **DF** | **MS** | **F (DFn, DFd)** | **P value** |
| Interaction | 117.7 | 1 | 117.7 | F (1, 23) = 0.03432 | P=0.8547 |
| Genotype | 3140 | 1 | 3140 | F (1, 23) = 0.9160 | P=0.3485 |
| Treatment | 1986 | 1 | 1986 | F (1, 23) = 0.5791 | P=0.4544 |

### Supplemental Table 5B – NeuN (count/square nm) across CA1 across genotype and treatment.

| Two-way ANOVA analysis | Males | | | | |
|---|---|---|---|---|---|
| **Source of Variation** | **% of total variation** | **P value** | **P value summary** | **Significant?** | |
| Interaction | 1.499 | 0.5239 | ns | No | |
| Genotype | 5.001 | 0.2488 | ns | No | |
| Treatment | 4.084 | 0.2962 | ns | No | |
| **ANOVA table** | **SS (Type III)** | **DF** | **MS** | **F (DFn, DFd)** | **P value** |
| Interaction | 1510 | 1 | 1510 | F (1, 25) = 0.4180 | P=0.5239 |
| Genotype | 5035 | 1 | 5035 | F (1, 25) = 1.394 | P=0.2488 |
| Treatment | 4112 | 1 | 4112 | F (1, 25) = 1.138 | P=0.2962 |

| Two-way ANOVA analysis | Females | | | | |
|---|---|---|---|---|---|
| **Source of Variation** | **% of total variation** | **P value** | **P value summary** | **Significant?** | |
| Interaction | 2.175 | 0.4794 | ns | No | |
| Genotype | 0.5478 | 0.7215 | ns | No | |
| Treatment | 0.7445 | 0.6779 | ns | No | |
| **ANOVA table** | **SS (Type III)** | **DF** | **MS** | **F (DFn, DFd)** | **P value** |
| Interaction | 1287 | 1 | 1287 | F (1, 23) = 0.5170 | P=0.4794 |
| Genotype | 324.2 | 1 | 324.2 | F (1, 23) = 0.1302 | P=0.7215 |
| Treatment | 440.6 | 1 | 440.6 | F (1, 23) = 0.1770 | P=0.6779 |

### Supplemental Table 5C – NeuN (count/square nm) across CA3 across genotype and treatment.

| Two-way ANOVA analysis | Males | | | | |
|---|---|---|---|---|---|
| **Source of Variation** | **% of total variation** | **P value** | **P value summary** | **Significant?** | |
| Interaction | 2.146 | 0.4423 | ns | No | |
| Genotype | 0.5909 | 0.6856 | ns | No | |
| Treatment | 9.428 | 0.1143 | ns | No | |
| **ANOVA table** | **SS (Type III)** | **DF** | **MS** | **F (DFn, DFd)** | **P value** |
| Interaction | 2599 | 1 | 2599 | F (1, 25) = 0.6094 | P=0.4423 |
| Genotype | 715.5 | 1 | 715.5 | F (1, 25) = 0.1678 | P=0.6856 |
| Treatment | 11417 | 1 | 11417 | F (1, 25) = 2.678 | P=0.1143 |

| Two-way ANOVA analysis | Females |
|---|---|

| Source of Variation | % of total variation | P value | P value summary | Significant? | |
|---|---|---|---|---|---|
| Interaction | 1.357 | 0.5673 | ns | No | |
| Genotype | 6.175 | 0.2282 | ns | No | |
| Treatment | 0.01490 | 0.9520 | ns | No | |
| **ANOVA table** | **SS (Type III)** | **DF** | **MS** | **F (DFn, DFd)** | **P value** |
| Interaction | 2189 | 1 | 2189 | F (1, 23) = 0.3368 | P=0.5673 |
| Genotype | 9960 | 1 | 9960 | F (1, 23) = 1.532 | P=0.2282 |
| Treatment | 24.03 | 1 | 24.03 | F (1, 23) = 0.003697 | P=0.9520 |

### Supplemental Table 5D – NeuN (count/square nm) across DG across genotype and treatment.

| Two-way ANOVA analysis | Males | | | | |
|---|---|---|---|---|---|
| **Source of Variation** | **% of total variation** | **P value** | **P value summary** | **Significant?** | |
| Interaction | 0.008966 | 0.9570 | ns | No | |
| Genotype | 4.674 | 0.2257 | ns | No | |
| Treatment | 18.97 | 0.0192 | * | Yes | |
| **ANOVA table** | **SS (Type III)** | **DF** | **MS** | **F (DFn, DFd)** | **P value** |
| Interaction | 11.02 | 1 | 11.02 | F (1, 25) = 0.002960 | P=0.9570 |
| Genotype | 5743 | 1 | 5743 | F (1, 25) = 1.543 | P=0.2257 |
| Treatment | 23303 | 1 | 23303 | F (1, 25) = 6.262 | P=0.0192 |

| Two-way ANOVA analysis | Females | | | | |
|---|---|---|---|---|---|
| **Source of Variation** | **% of total variation** | **P value** | **P value summary** | **Significant?** | |
| Interaction | 0.9804 | 0.6305 | ns | No | |
| Genotype | 0.05111 | 0.9123 | ns | No | |
| Treatment | 4.301 | 0.3178 | ns | No | |
| **ANOVA table** | **SS (Type III)** | **DF** | **MS** | **F (DFn, DFd)** | **P value** |
| Interaction | 1489 | 1 | 1489 | F (1, 23) = 0.2377 | P=0.6305 |
| Genotype | 77.62 | 1 | 77.62 | F (1, 23) = 0.01239 | P=0.9123 |
| Treatment | 6532 | 1 | 6532 | F (1, 23) = 1.043 | P=0.3178 |

### Supplemental Table 5E– NeuN (count/square nm) across SB across genotype and treatment.

| Two-way ANOVA analysis | Males | | | | |
|---|---|---|---|---|---|
| **Source of Variation** | **% of total variation** | **P value** | **P value summary** | **Significant?** | |
| Interaction | 1.017 | 0.5675 | ns | No | |
| Genotype | 22.47 | 0.0116 | * | Yes | |
| Treatment | 1.315 | 0.5160 | ns | No | |
| **ANOVA table** | **SS (Type III)** | **DF** | **MS** | **F (DFn, DFd)** | **P value** |
| Interaction | 2449 | 1 | 2449 | F (1, 25) = 0.3357 | P=0.5675 |
| Genotype | 54098 | 1 | 54098 | F (1, 25) = 7.416 | P=0.0116 |
| Treatment | 3166 | 1 | 3166 | F (1, 25) = 0.4340 | P=0.5160 |

| Two-way ANOVA analysis | Females | | | |
|---|---|---|---|---|
| **Source of Variation** | **% of total variation** | **P value** | **P value summary** | **Significant?** |
| Interaction | 1.811 | 0.5204 | ns | No |
| Genotype | 0.1142 | 0.8712 | ns | No |
| Treatment | 0.4078 | 0.7596 | ns | No |

| ANOVA table | SS (Type III) | DF | MS | F (DFn, DFd) | P value |
|---|---|---|---|---|---|
| Interaction | 1599 | 1 | 1599 | F (1, 23) = 0.4260 | P=0.5204 |
| Genotype | 100.9 | 1 | 100.9 | F (1, 23) = 0.02687 | P=0.8712 |
| Treatment | 360.1 | 1 | 360.1 | F (1, 23) = 0.09593 | P=0.7596 |

### Supplemental Table 5F – NeuN (count/square nm) across HL across genotype and treatment.

| Two-way ANOVA analysis | Males | | | | |
|---|---|---|---|---|---|
| Source of Variation | % of total variation | P value | P value summary | Significant? | |
| Interaction | 2.983 | 0.3617 | ns | No | |
| Genotype | 2.768 | 0.3793 | ns | No | |
| Treatment | 8.054 | 0.1393 | ns | No | |
| ANOVA table | SS (Type III) | DF | MS | F (DFn, DFd) | P value |
| Interaction | 3205 | 1 | 3205 | F (1, 25) = 0.8635 | P=0.3617 |
| Genotype | 2974 | 1 | 2974 | F (1, 25) = 0.8013 | P=0.3793 |
| Treatment | 8654 | 1 | 8654 | F (1, 25) = 2.331 | P=0.1393 |

| Two-way ANOVA analysis | Females | | | | |
|---|---|---|---|---|---|
| Source of Variation | % of total variation | P value | P value summary | Significant? | |
| Interaction | 11.05 | 0.0773 | ns | No | |
| Genotype | 15.48 | 0.0391 | * | Yes | |
| Treatment | 0.004287 | 0.9713 | ns | No | |
| ANOVA table | SS (Type III) | DF | MS | F (DFn, DFd) | P value |
| Interaction | 18735 | 1 | 18735 | F (1, 23) = 3.420 | P=0.0773 |
| Genotype | 26237 | 1 | 26237 | F (1, 23) = 4.789 | P=0.0391 |
| Treatment | 7.265 | 1 | 7.265 | F (1, 23) = 0.001326 | P=0.9713 |

### Supplemental Table 5G – NeuN (count/square nm) across GCL across genotype and treatment.

| Two-way ANOVA analysis | Males | | | | |
|---|---|---|---|---|---|
| Source of Variation | % of total variation | P value | P value summary | Significant? | |
| Interaction | 0.03311 | 0.9276 | ns | No | |
| Genotype | 1.290 | 0.5717 | ns | No | |
| Treatment | 0.4328 | 0.7427 | ns | No | |
| ANOVA table | SS (Type III) | DF | MS | F (DFn, DFd) | P value |
| Interaction | 368.8 | 1 | 368.8 | F (1, 25) = 0.008429 | P=0.9276 |
| Genotype | 14366 | 1 | 14366 | F (1, 25) = 0.3284 | P=0.5717 |
| Treatment | 4819 | 1 | 4819 | F (1, 25) = 0.1102 | P=0.7427 |

| Two-way ANOVA analysis | Females | | | | |
|---|---|---|---|---|---|
| Source of Variation | % of total variation | P value | P value summary | Significant? | |
| Interaction | 0.3930 | 0.7624 | ns | No | |
| Genotype | 2.428 | 0.4547 | ns | No | |
| Treatment | 0.8357 | 0.6597 | ns | No | |
| ANOVA table | SS (Type III) | DF | MS | F (DFn, DFd) | P value |
| Interaction | 9252 | 1 | 9252 | F (1, 23) = 0.09360 | P=0.7624 |
| Genotype | 57168 | 1 | 57168 | F (1, 23) = 0.5783 | P=0.4547 |
| Treatment | 19677 | 1 | 19677 | F (1, 23) = 0.1991 | P=0.6597 |

**Supplemental Table 5 – Neuronal (NeuN) antibody density across the hippocampus.** Values represent percent count per square nm across the hippocampus and its subregions for both male and female rats across four experimental groups (WTNT, TGNT. WTTR, and TGTR). Data was analyzed using two- way ANOVAs. Abbreviations: HC= hippocampus; CA= cornu ammonis; DG= dentate gyrus; SB= subiculum; HL= hilar region, GCL= granule cell layer, WTNT= wild type not treated, males n=7; females n=7; TGNT= transgenic not treated, males n=7; females n=7); WTTR= wild type BT-11 treated, males n=8; females n=6; TGTR= transgenic BT-11 treated, males n=7; females n=7.

**Supplemental Table 6A – Phosphorylated tau across Dorsal HC across genotype and treatment.**

| **Two-way ANOVA analysis** | **Males** | | | | |
|---|---|---|---|---|---|
| **Source of Variation** | **% of total variation** | **P value** | **P value summary** | **Significant?** | |
| Interaction | 3.445 | 0.3163 | ns | No | |
| Genotype | 6.982 | 0.1576 | ns | No | |
| Treatment | 8.111e-005 | 0.9961 | ns | No | |
| **ANOVA table** | **SS (Type III)** | **DF** | **MS** | **F (DFn, DFd)** | **P value** |
| Interaction | 29169 | 1 | 29169 | F (1, 27) = 1.043 | P=0.3163 |
| Genotype | 59111 | 1 | 59111 | F (1, 27) = 2.113 | P=0.1576 |
| Treatment | 0.6867 | 1 | 0.6867 | F (1, 27) = 2.455e-005 | P=0.9961 |

| **Two-way ANOVA analysis** | **Females** | | | | |
|---|---|---|---|---|---|
| **Source of Variation** | **% of total variation** | **P value** | **P value summary** | **Significant?** | |
| Interaction | 0.005314 | 0.9711 | ns | No | |
| Genotype | 1.758 | 0.5119 | ns | No | |
| Treatment | 6.804 | 0.2030 | ns | No | |
| **ANOVA table** | **SS (Type III)** | **DF** | **MS** | **F (DFn, DFd)** | **P value** |
| Interaction | 27.39 | 1 | 27.39 | F (1, 23) = 0.001341 | P=0.9711 |
| Genotype | 9065 | 1 | 9065 | F (1, 23) = 0.4437 | P=0.5119 |
| Treatment | 35072 | 1 | 35072 | F (1, 23) = 1.717 | P=0.2030 |

**Supplemental Table 6B – Phosphorylated tau across Dorsal HC across treatment in TG rats.**

| **Unpaired t test** | **Males** |
|---|---|
| P value | 0.3802 |
| P value summary | ns |
| Significantly different (P < 0.05)? | No |
| One- or two-tailed P value? | Two-tailed |
| t, df | t=0.9061, df=14 |

| **Unpaired t test** | **Females** |
|---|---|
| P value | 0.4356 |
| P value summary | ns |
| Significantly different (P < 0.05)? | No |
| One- or two-tailed P value? | Two-tailed |
| t, df | t=0.8065, df=12 |

**Supplemental Table 6C – Phosphorylated tau across CA1 across treatment in TG rats.**

| Unpaired t test | Males |
|---|---|
| P value | 0.4036 |
| P value summary | ns |
| Significantly different (P < 0.05)? | No |
| One- or two-tailed P value? | Two-tailed |
| t, df | t=0.8657, df=12 |

| Unpaired t test | Females |
|---|---|
| P value | 0.7289 |
| P value summary | ns |
| Significantly different (P < 0.05)? | No |
| One- or two-tailed P value? | Two-tailed |
| t, df | t=0.3548, df=12 |

### Supplemental Table 6D – Phosphorylated tau across CA3 across treatment in TG rats.

| Unpaired t test | Males |
|---|---|
| P value | 0.4000 |
| P value summary | ns |
| Significantly different (P < 0.05)? | No |
| One- or two-tailed P value? | Two-tailed |
| t, df | t=0.8726, df=12 |

| Unpaired t test | Females |
|---|---|
| P value | 0.4987 |
| P value summary | ns |
| Significantly different (P < 0.05)? | No |
| One- or two-tailed P value? | Two-tailed |
| t, df | t=0.6976, df=12 |

### Supplemental Table 6E – Phosphorylated tau across DG across treatment in TG rats.

| Unpaired t test | Males |
|---|---|
| P value | 0.5139 |
| P value summary | ns |
| Significantly different (P < 0.05)? | No |
| One- or two-tailed P value? | Two-tailed |
| t, df | t=0.6727, df=12 |

| Unpaired t test | Females |
|---|---|
| P value | 0.5390 |
| P value summary | ns |
| Significantly different (P < 0.05)? | No |
| One- or two-tailed P value? | Two-tailed |
| t, df | t=0.6324, df=12 |

### Supplemental Table 6F – Phosphorylated tau across SB across treatment in TG rats

| Unpaired t test | Males |
|---|---|
| P value | 0.7042 |
| P value summary | ns |
| Significantly different (P < 0.05)? | No |
| One- or two-tailed P value? | Two-tailed |
| t, df | t=0.3888, df=12 |

| Unpaired t test | Females |
|---|---|
| P value | 0.2171 |
| P value summary | ns |
| Significantly different (P < 0.05)? | No |

| | |
|---|---|
| One- or two-tailed P value? | Two-tailed |
| t, df | t=1.303, df=12 |

**Supplemental Table 6G – Phosphorylated tau across HL across treatment in TG rats.**

| Unpaired t test | Males |
|---|---|
| P value | 0.1878 |
| P value summary | ns |
| Significantly different (P < 0.05)? | No |
| One- or two-tailed P value? | Two-tailed |
| t, df | t=1.397, df=12 |

| Unpaired t test | Females |
|---|---|
| P value | 0.7289 |
| P value summary | ns |
| Significantly different (P < 0.05)? | No |
| One- or two-tailed P value? | Two-tailed |
| t, df | t=0.3548, df=12 |

**Supplemental Table 6H – Phosphorylated tau across GCL across treatment in TG rats.**

| Unpaired t test | Males |
|---|---|
| P value | 0.1573 |
| P value summary | ns |
| Significantly different (P < 0.05)? | No |
| One- or two-tailed P value? | Two-tailed |
| t, df | t=1.508, df=12 |

| Unpaired t test | Females |
|---|---|
| P value | 0.9686 |
| P value summary | ns |
| Significantly different (P < 0.05)? | No |
| One- or two-tailed P value? | Two-tailed |
| t, df | t=0.04021, df=12 |

**Supplemental Table 6 –Phosphorylated Tau (AT8 antibody) density across the hippocampus.** Values represent percent count per square nm across the hippocampus for both male and female rats across four experimental groups (WTNT, TGNT. WTTR, and TGTR). Data was analyzed using two- way ANOVA (A). Data was analyzed by unpaired t tests across subregions for both males and females across two experimental groups (TGNT and TGTR) (B-H). Abbreviations: HC= hippocampus; CA= cornu ammonis; DG= dentate gyrus; SB= subiculum; HL= hilar region, GCL= granule cell layer, WTNT= wild type not treated, males n=7; females n=7; TGNT= transgenic not treated, males n=7; females n=7); WTTR= wild type BT-11 treated, males n=8; females n=6; TGTR= transgenic BT-11 treated, males n=7; females n=7.

**Supplemental Table 7A – LANCL2 levels in total hippocampal lysates across genotype and treatment.**

| **Two-way ANOVA analysis** | **Males** | | | | |
|---|---|---|---|---|---|
| **Source of Variation** | **% of total variation** | **P value** | **P value summary** | **Significant?** | |
| Interaction | 9.245 | 0.3856 | ns | No | |
| genotype | 2.915 | 0.6203 | ns | No | |
| treatment | 0.006014 | 0.9819 | ns | No | |
| **ANOVA table** | **SS** | **DF** | **MS** | **F (DFn, DFd)** | **P value** |
| Interaction | 0.04452 | 1 | 0.04452 | F (1, 8) = 0.8421 | P=0.3856 |

| | | | | | |
|---|---|---|---|---|---|
| genotype | 0.01404 | 1 | 0.01404 | F (1, 8) = 0.2655 | P=0.6203 |
| treatment | 2.896e-005 | 1 | 2.896e-005 | F (1, 8) = 0.0005478 | P=0.9819 |

| **Two-way ANOVA analysis** | **Females** | | | | |
|---|---|---|---|---|---|
| **Source of Variation** | **% of total variation** | **P value** | **P value summary** | **Significant?** | |
| Interaction | 11.43 | 0.3355 | ns | No | |
| genotype | 1.406 | 0.7286 | ns | No | |
| treatment | 0.04467 | 0.9505 | ns | No | |
| **ANOVA table** | **SS** | **DF** | **MS** | **F (DFn, DFd)** | **P value** |
| Interaction | 0.08662 | 1 | 0.08662 | F (1, 8) = 1.050 | P=0.3355 |
| genotype | 0.01065 | 1 | 0.01065 | F (1, 8) = 0.1292 | P=0.7286 |
| treatment | 0.0003384 | 1 | 0.0003384 | F (1, 8) = 0.004102 | P=0.9505 |

### Supplemental Table 7B – LANCL2 levels in hippocampal cytoplasmic fractions across genotype and treatment.

| **Two-way ANOVA analysis** | **Males** | | | | |
|---|---|---|---|---|---|
| **Source of Variation** | **% of total variation** | **P value** | **P value summary** | **Significant?** | |
| Interaction | 2.554 | 0.5922 | ns | No | |
| genotype | 14.25 | 0.2241 | ns | No | |
| treatment | 17.55 | 0.1818 | ns | No | |
| **ANOVA table** | **SS** | **DF** | **MS** | **F (DFn, DFd)** | **P value** |
| Interaction | 0.009154 | 1 | 0.009154 | F (1, 8) = 0.3112 | P=0.5922 |
| genotype | 0.05107 | 1 | 0.05107 | F (1, 8) = 1.736 | P=0.2241 |
| treatment | 0.06290 | 1 | 0.06290 | F (1, 8) = 2.139 | P=0.1818 |

| **Two-way ANOVA analysis** | **Females** | | | | |
|---|---|---|---|---|---|
| **Source of Variation** | **% of total variation** | **P value** | **P value summary** | **Significant?** | |
| Interaction | 0.1148 | 0.8896 | ns | No | |
| genotype | 52.84 | 0.0153 | * | Yes | |
| treatment | 2.269 | 0.5421 | ns | No | |
| **ANOVA table** | **SS** | **DF** | **MS** | **F (DFn, DFd)** | **P value** |
| Interaction | 0.0003705 | 1 | 0.0003705 | F (1, 8) = 0.02052 | P=0.8896 |
| genotype | 0.1705 | 1 | 0.1705 | F (1, 8) = 9.442 | P=0.0153 |
| treatment | 0.007320 | 1 | 0.007320 | F (1, 8) = 0.4054 | P=0.5421 |

### Supplemental Table 7C – LANCL2 levels in hippocampal nuclear fractions across genotype and treatment.

| **Two-way ANOVA analysis** | **Males** | | | | |
|---|---|---|---|---|---|
| **Source of Variation** | **% of total variation** | **P value** | **P value summary** | **Significant?** | |
| Interaction | 0.06776 | 0.9308 | ns | No | |
| genotype | 20.78 | 0.1551 | ns | No | |
| treatment | 11.72 | 0.2723 | ns | No | |
| **ANOVA table** | **SS** | **DF** | **MS** | **F (DFn, DFd)** | **P value** |
| Interaction | 0.0007363 | 1 | 0.0007363 | F (1, 8) = 0.008038 | P=0.9308 |
| genotype | 0.2258 | 1 | 0.2258 | F (1, 8) = 2.465 | P=0.1551 |
| treatment | 0.1273 | 1 | 0.1273 | F (1, 8) = 1.390 | P=0.2723 |

| **Two-way ANOVA analysis** | **Females** | | | |
|---|---|---|---|---|
| **Source of Variation** | **% of total variation** | **P value** | **P value summary** | **Significant?** |
| Interaction | 1.073 | 0.6458 | ns | No |
| genotype | 57.65 | 0.0081 | ** | Yes |

| | | | | | |
|---|---|---|---|---|---|
| treatment | 3.631 | 0.4053 | ns | No | |
| **ANOVA table** | **SS** | **DF** | **MS** | **F (DFn, DFd)** | **P value** |
| Interaction | 0.06163 | 1 | 0.06163 | F (1, 8) = 0.2280 | P=0.6458 |
| genotype | 3.312 | 1 | 3.312 | F (1, 8) = 12.25 | P=0.0081 |
| treatment | 0.2086 | 1 | 0.2086 | F (1, 8) = 0.7716 | P=0.4053 |

**Supplemental Table 7 – LANCL2 levels across genotype and treatment.** Values represent the percentage of the pixel ratio for LANCL2 over the respective loading controls, GAPDH for total and cytoplasmic fractions (A, B), and lamin B1 for the nuclear fraction (C). Two-way ANOVA analysis was performed for the four groups WTNT, WTTR, TGNT, TGTR for both males and females, three rats per group. Abbreviations: WTNT= wild type not treated, TGNT= transgenic not treated, WTTR= wild type BT-11 treated, TGTR= transgenic BT-11 treated.

**Supplemental Table 8 – pCREB levels in total hippocampal lysate across genotype and treatment.**

| **Two-way ANOVA analysis** | **Males** | | | | |
|---|---|---|---|---|---|
| **Source of Variation** | **% of total variation** | **P value** | **P value summary** | **Significant?** | |
| Interaction | 19.72 | 0.1758 | ns | No | |
| genotype | 2.490 | 0.6120 | ns | No | |
| treatment | 6.258 | 0.4271 | ns | No | |
| **ANOVA table** | **SS** | **DF** | **MS** | **F (DFn, DFd)** | **P value** |
| Interaction | 2.310 | 1 | 2.310 | F (1, 8) = 2.206 | P=0.1758 |
| genotype | 0.2917 | 1 | 0.2917 | F (1, 8) = 0.2785 | P=0.6120 |
| treatment | 0.7331 | 1 | 0.7331 | F (1, 8) = 0.7000 | P=0.4271 |

| **Two-way ANOVA analysis** | **Females** | | | | |
|---|---|---|---|---|---|
| **Source of Variation** | **% of total variation** | **P value** | **P value summary** | **Significant?** | |
| Interaction | 8.182 | 0.3132 | ns | No | |
| genotype | 26.61 | 0.0882 | ns | No | |
| treatment | 8.712 | 0.2990 | ns | No | |
| **ANOVA table** | **SS** | **DF** | **MS** | **F (DFn, DFd)** | **P value** |
| Interaction | 5.971 | 1 | 5.971 | F (1, 8) = 1.158 | P=0.3132 |
| genotype | 19.42 | 1 | 19.42 | F (1, 8) = 3.767 | P=0.0882 |
| treatment | 6.358 | 1 | 6.358 | F (1, 8) = 1.233 | P=0.2990 |

**Supplemental Table 8 – pCREB levels across genotype and treatment.** Values represent the percentage of the pixel ratio for pCREB over CREB normalized to a GAPDH loading control. Two-way ANOVA analysis was performed for the four groups WTNT, WTTR, TGNT, TGTR for both males and females, three rats per group. Abbreviations: WTNT= wild type not treated, TGNT= transgenic not treated, WTTR= wild type BT-11 treated, TGTR= transgenic BT-11 treated.

**Supplemental Table 9 – Path length during habituation in aPAT across genotype and treatment.**

| **Two-way ANOVA analysis** | **Males** | | | | |
|---|---|---|---|---|---|
| **Source of Variation** | **% of total variation** | **P value** | **P value summary** | **Significant?** | |
| Interaction | 0.4738 | 0.6085 | ns | No | |
| Genotype | 0.3289 | 0.6694 | ns | No | |
| Treatment | 4.642 | 0.1127 | ns | No | |
| **ANOVA table** | **SS (Type III)** | **DF** | **MS** | **F (DFn, DFd)** | **P value** |
| Interaction | 3.330 | 1 | 3.330 | F (1, 53) = 0.2656 | P=0.6085 |
| Genotype | 2.312 | 1 | 2.312 | F (1, 53) = 0.1843 | P=0.6694 |
| Treatment | 32.62 | 1 | 32.62 | F (1, 53) = 2.602 | P=0.1127 |

| Two-way ANOVA analysis | Females | | | | |
|---|---|---|---|---|---|
| **Source of Variation** | **% of total variation** | **P value** | **P value summary** | **Significant?** | |
| Interaction | 0.03803 | 0.8712 | ns | No | |
| Genotype | 3.638 | 0.1160 | ns | No | |
| Treatment | 0.3886 | 0.6045 | ns | No | |
| **ANOVA table** | **SS (Type III)** | **DF** | **MS** | **F (DFn, DFd)** | **P value** |
| Interaction | 0.3085 | 1 | 0.3085 | F (1, 67) = 0.02651 | P=0.8712 |
| Genotype | 29.52 | 1 | 29.52 | F (1, 67) = 2.536 | P=0.1160 |
| Treatment | 3.152 | 1 | 3.152 | F (1, 67) = 0.2709 | P=0.6045 |

**Supplemental Table 9 – Path length was unchanged across genotype and treatment across trials.** Path length was determined for each experimental group (WTNT, TGNT, WTTR, and TGTR) for both males and females during a habituation trial. 2-way ANOVA analysis is used to assess effects of drug treatment and genotype (Abbreviations: WTNT= wild type not treated, TGNT= transgenic not treated, WTTR= wild type BT-11 treated, TGTR= transgenic BT-11 treated.

A

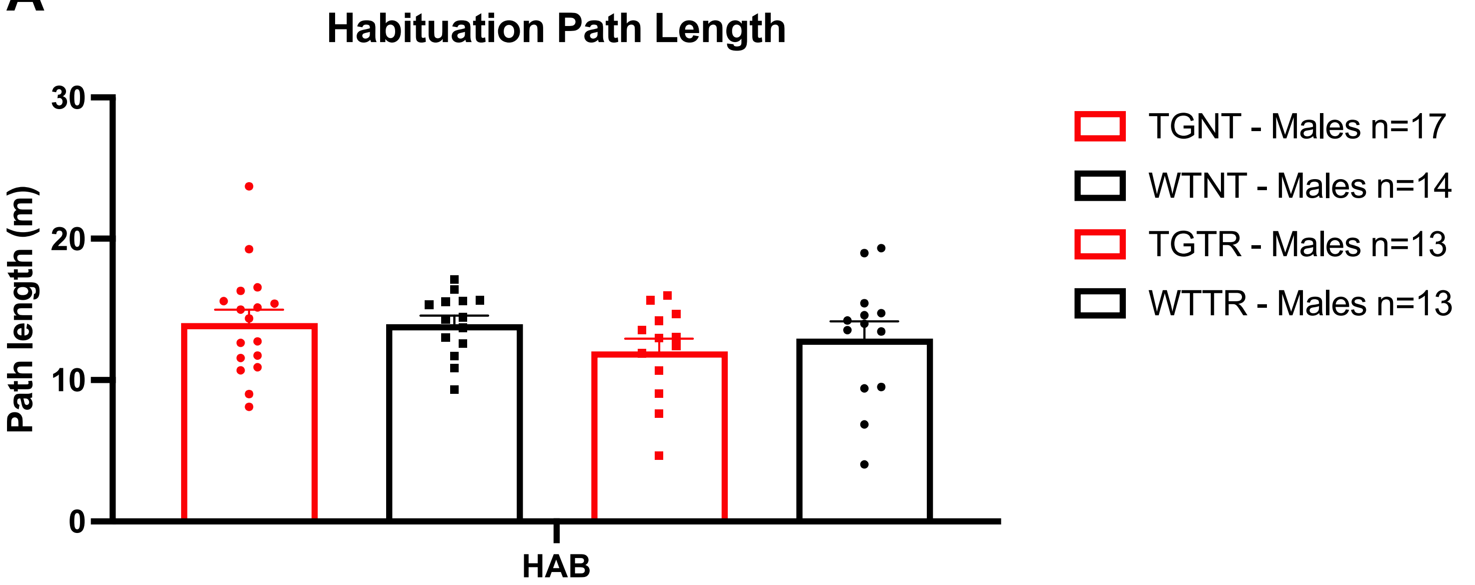

B

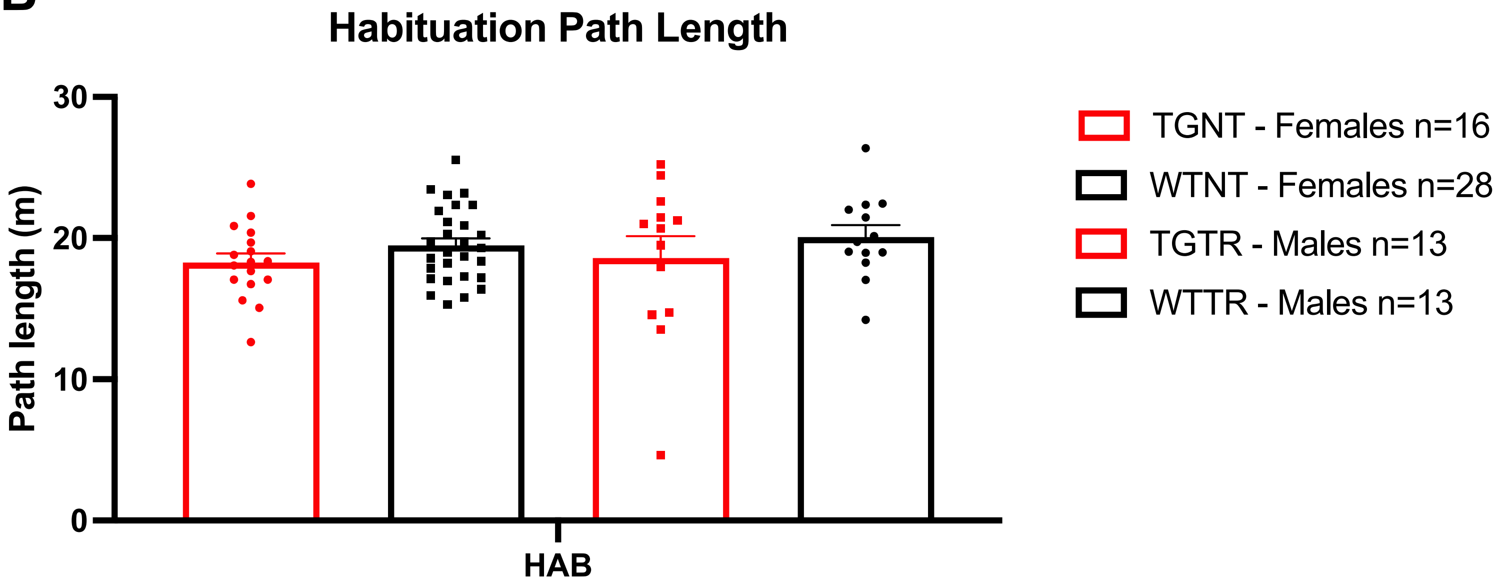

**Supplemental Figure 1**: Path length was determined for each experimental group (WTNT, TGNT, WTTR, and TGTR) for both males **(A)** and females **(B)** during a habituation trial. The Y-axis represents distance traveled in meters over a 10-minute period. 2-way ANOVA analysis was used to assess the effects of drug treatment and genotype Abbreviations: HAB= habituation, WTNT= wild type not treated, TGNT= transgenic not treated, WTTR= wild type BT-11 treated, TGTR= transgenic BT-11 treated.

A

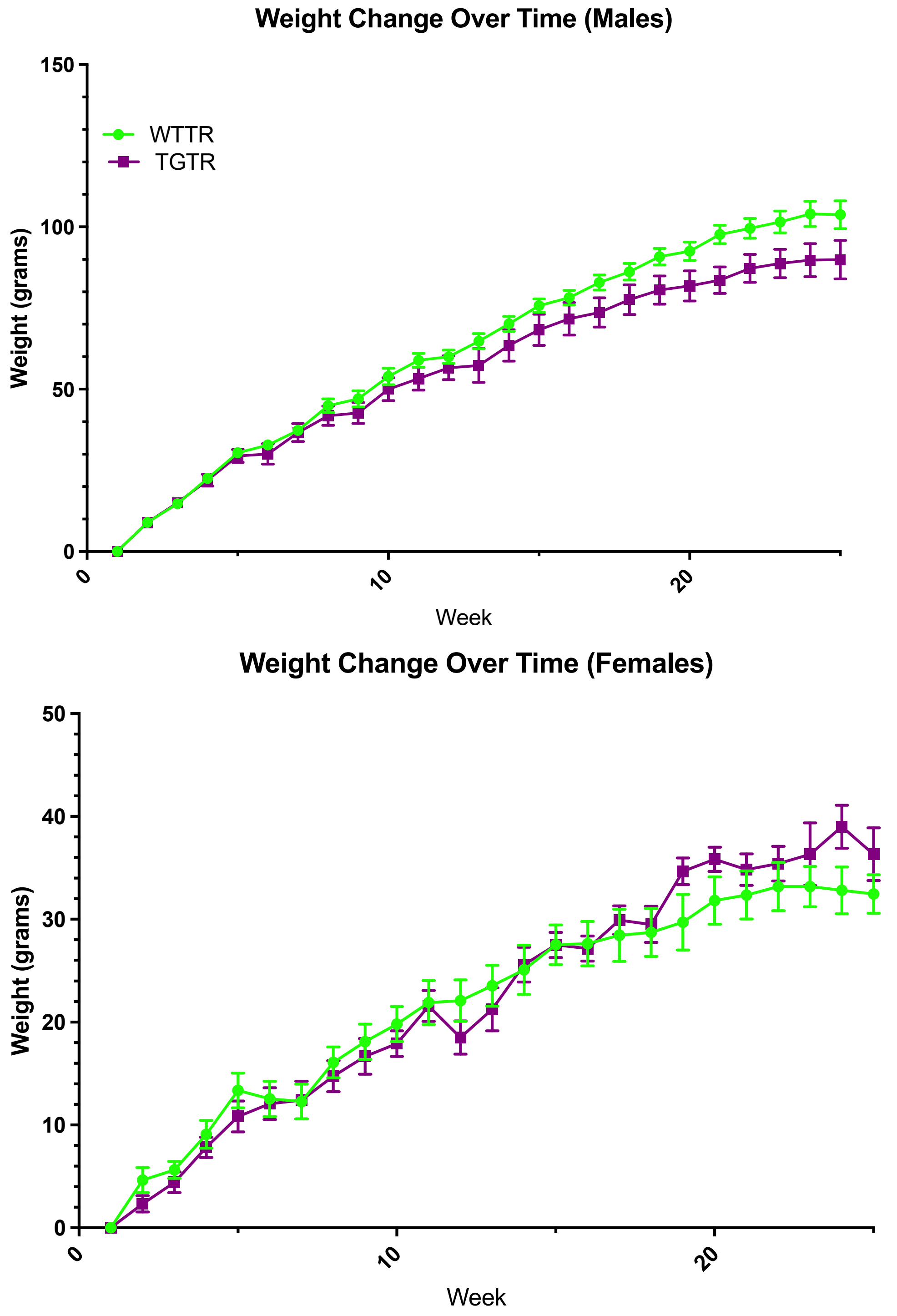

**Supplemental Figure 2**: Males (WTTR vs TGTR) show significance in weight gain over time [F (3.414, 85.36) = 492.1, p<0.0001] but no change in weight gain by genotype [F (1, 25) = 3, p=0.0956] **(A)**. Females (WTTR vs TGTR) show a significance in weight gain over time [F (3.463, 72.71) =149.4, p<0.0001] and no genotype significance [F (1, 21) = 0.08520], p=0.7732) **(B).** Results were analyzed by 2-way repeated-measures ANOVAs. Abbreviations: WTNT= wild type not treated, TGNT= transgenic not treated, WTTR= wild type BT-11 treated, TGTR= transgenic BT-11 treated.

A

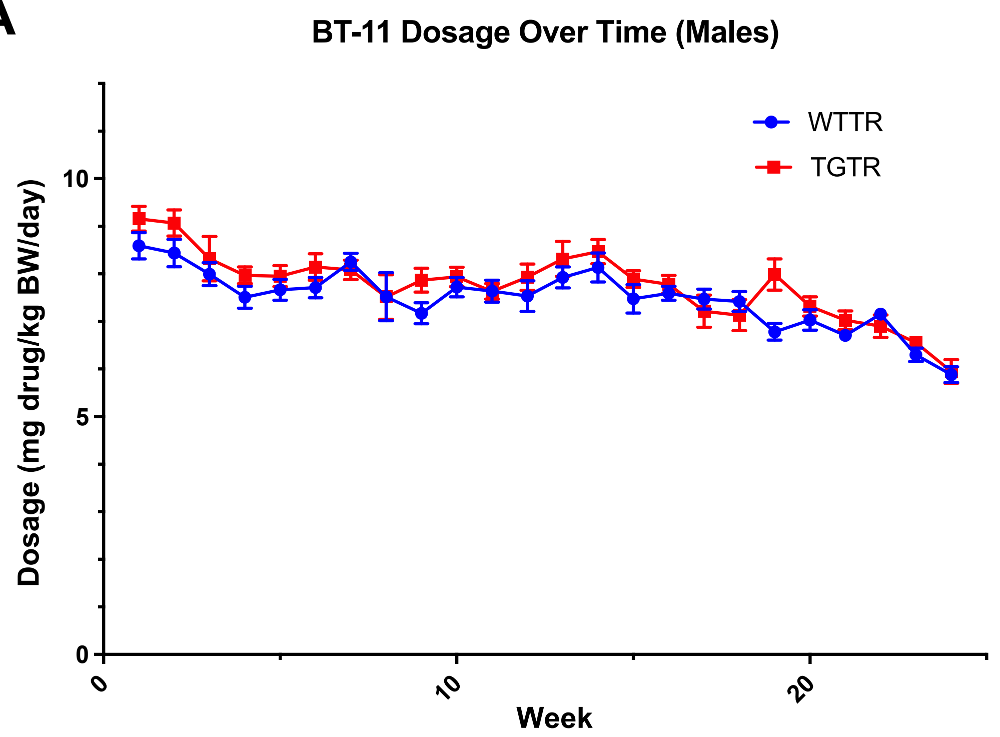

B

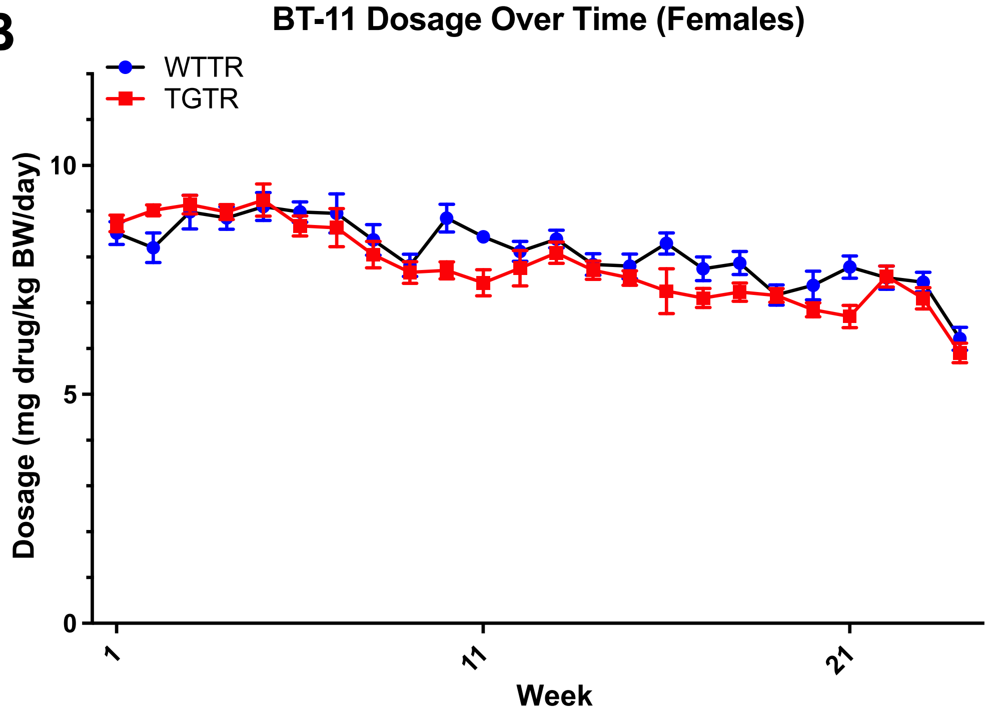

**Supplemental Figure 3**: Based on weekly body weights and food containing BT-11 the actual average of BT-11 consumed by WTTR and TGTR male rats respectively was 7.437 mg/kg bw/day, and 7.672 mg/kg bw/day. WTTR and TGTR showed no genotype difference [F (1, 25) =2.026, p=0.1669]. There was a significant difference by week of treatment [F (4.068, 101.7) =17.41, p<0.0001] **(A)**. The average of BT-11 consumed by WTTR and TGTR female rats respectively was 8.099 mg/kg bw/day and 7.649 mg/kg bw/day WTTR and TGTR showed no genotype difference [F (1, 21) = 3.163, p=0.0898]. There was a significant difference by week of treatment [F (5.394, 113.3) = 19.89, p<0.0001] **(B)**. Data was analyzed using 2-way repeated-measures ANOVA; bw=body WTNT= wild type not treated, TGNT= transgenic not treated, WTTR= wild type BT-11 treated, TGTR= transgenic BT-11 treated.

**Male Neuron (Neun) and Phosphorylated Tau (AT8) Representative Images**

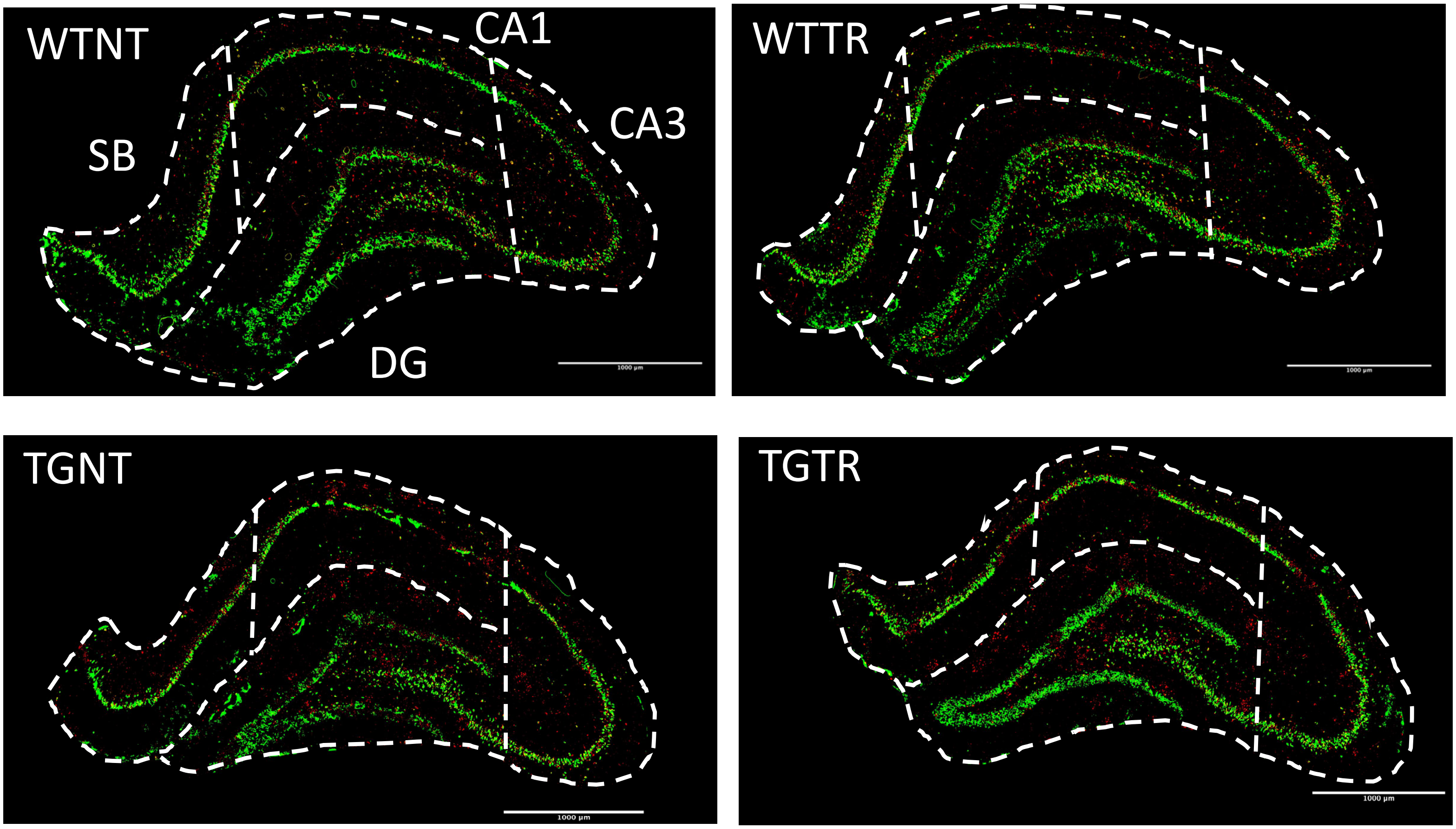

**Supplemental Figure 4**: Representative IHC images of phosphorylated tau (AT8) in red and neurons (NeuN) in green in male rats. WTNT= wild type not treated, TGNT= transgenic not treated, WTTR= wild type BT-11 treated, TGTR= transgenic BT-11 treated. 1000µm scale bars.

### Transgenic Male Neuron (Neun) and Phosphorylated Tau (AT8) Representative Images

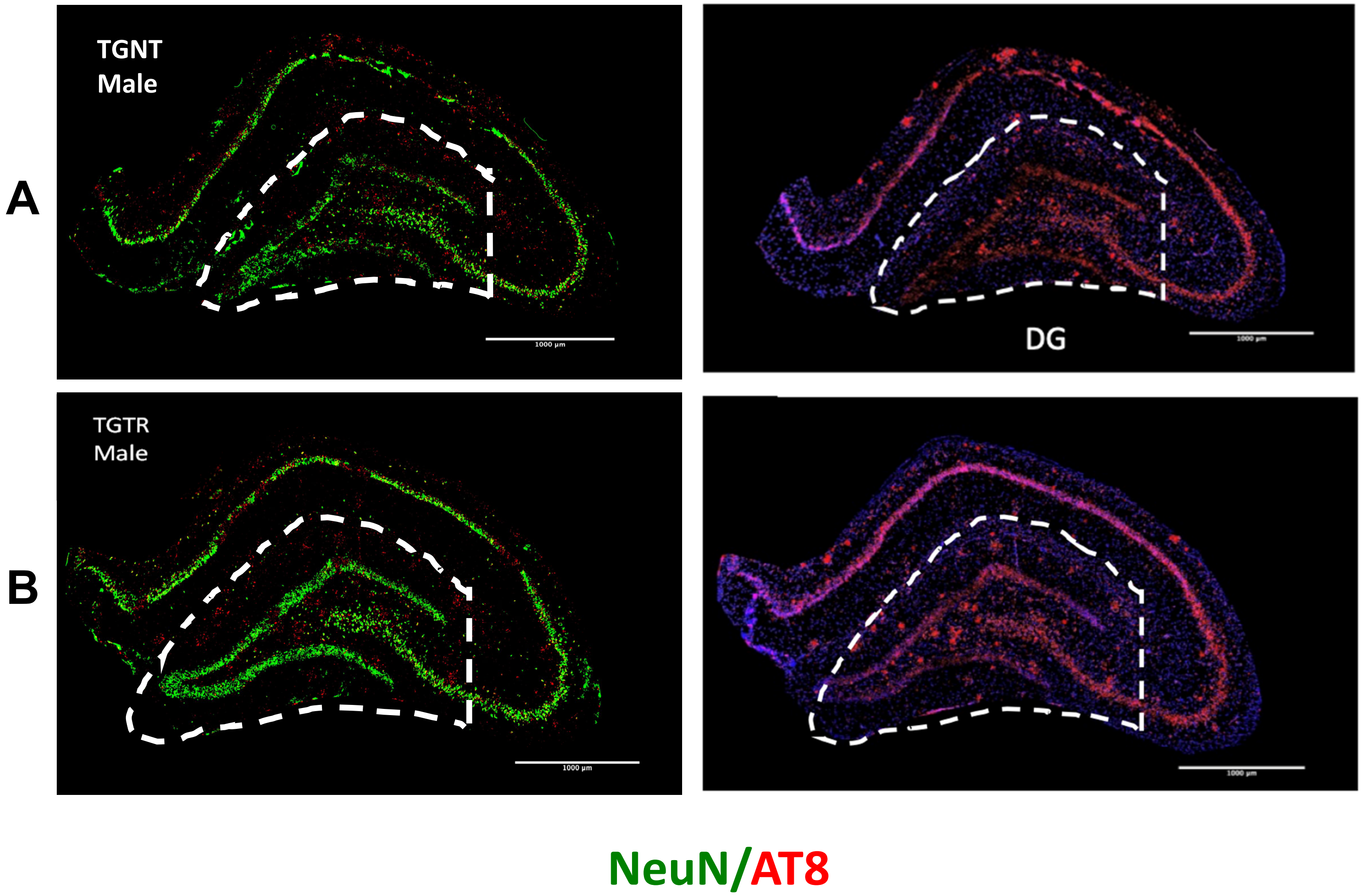

**Supplemental Figure 5**: Representative IHC images of neurons (NeuN, green) and Phosphorylated tau (AT8, red) in TGNT (A), TGTR (B) male rats**.** Abbreviations: TGNT= transgenic not treated, TGTR= transgenic BT-11 treated, DG= Dentate Gyrus. 1000µm scale bars.

**A**

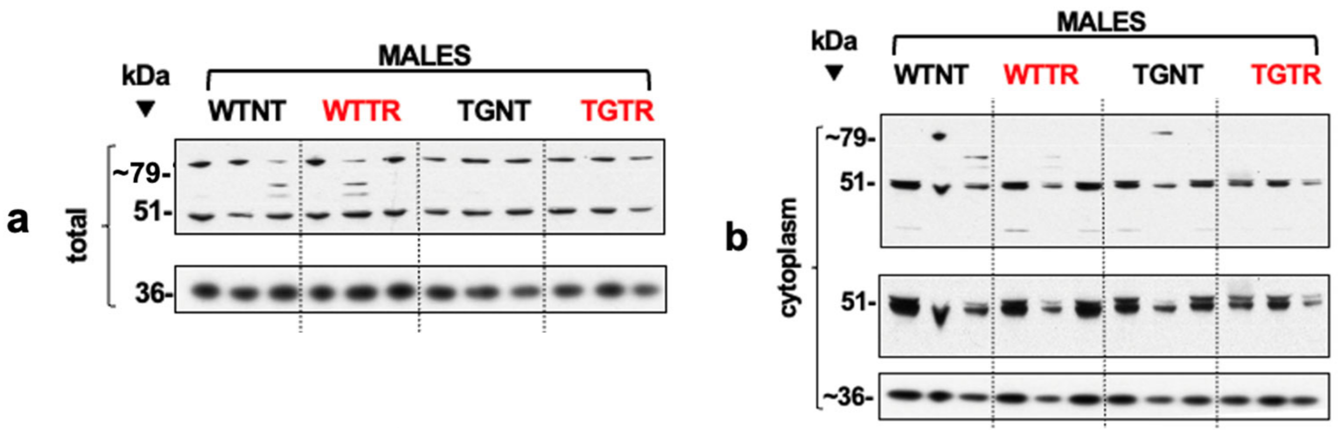

**B**

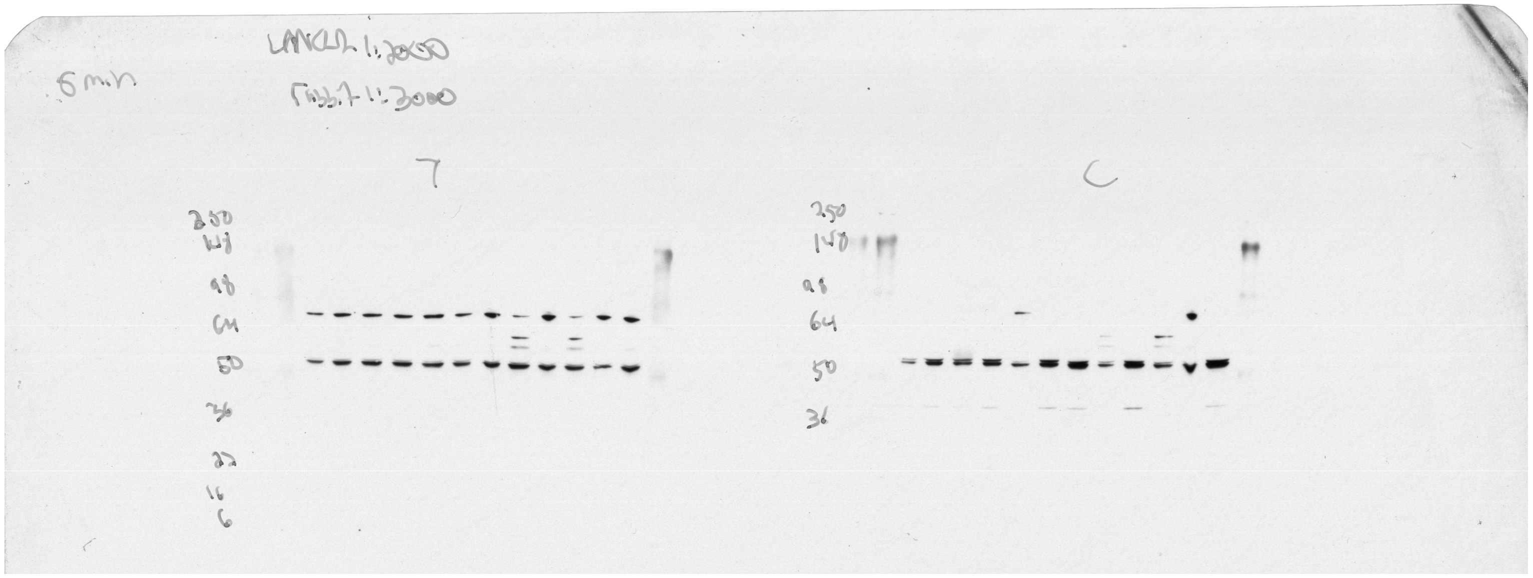

**Supplemental Figure 6: A –** Panel shown in Figure 7 of the manuscript. **B -** Scan of the film used to visualize LANCL2 western blots. Molecular weight markers are labelled on the left of the blots. The diagonal line on the corner was used to mark the film's orientation. The panels from figure 7 are flipped horizontally from the original blots to show WTNT, WTTR, TGNT, and TGTR samples from left to right. Hippocampal tissue was analyzed by western blotting. Total (left gel) and cytoplasmic fraction (right gel) from male rats are shown. Abbreviations: WTNT – wild-type not treated, TGNT – transgenic not treated, WTTR – wild-type BT-11 treated, TGTR – transgenic BT-11treated.

**A**

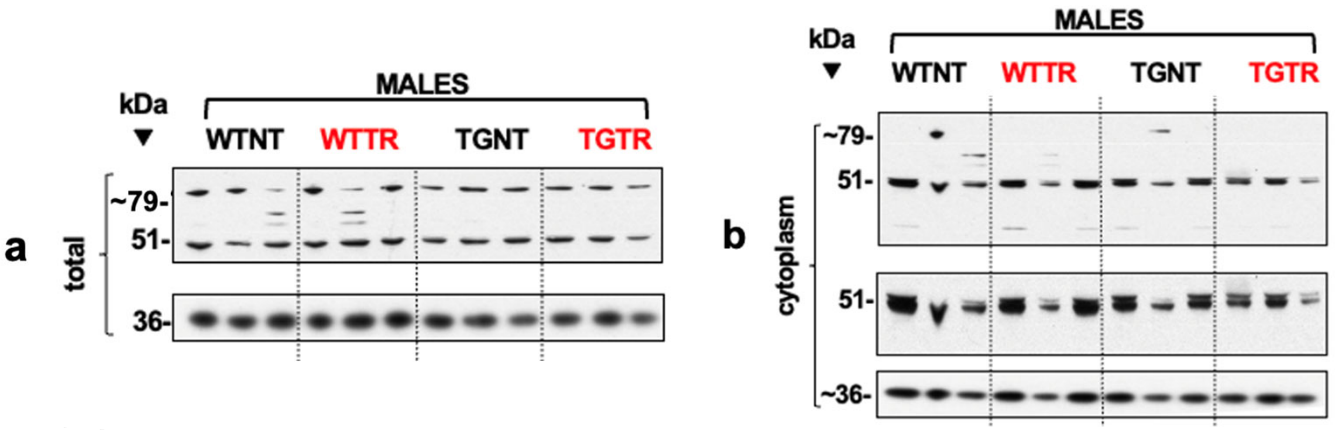

**B**

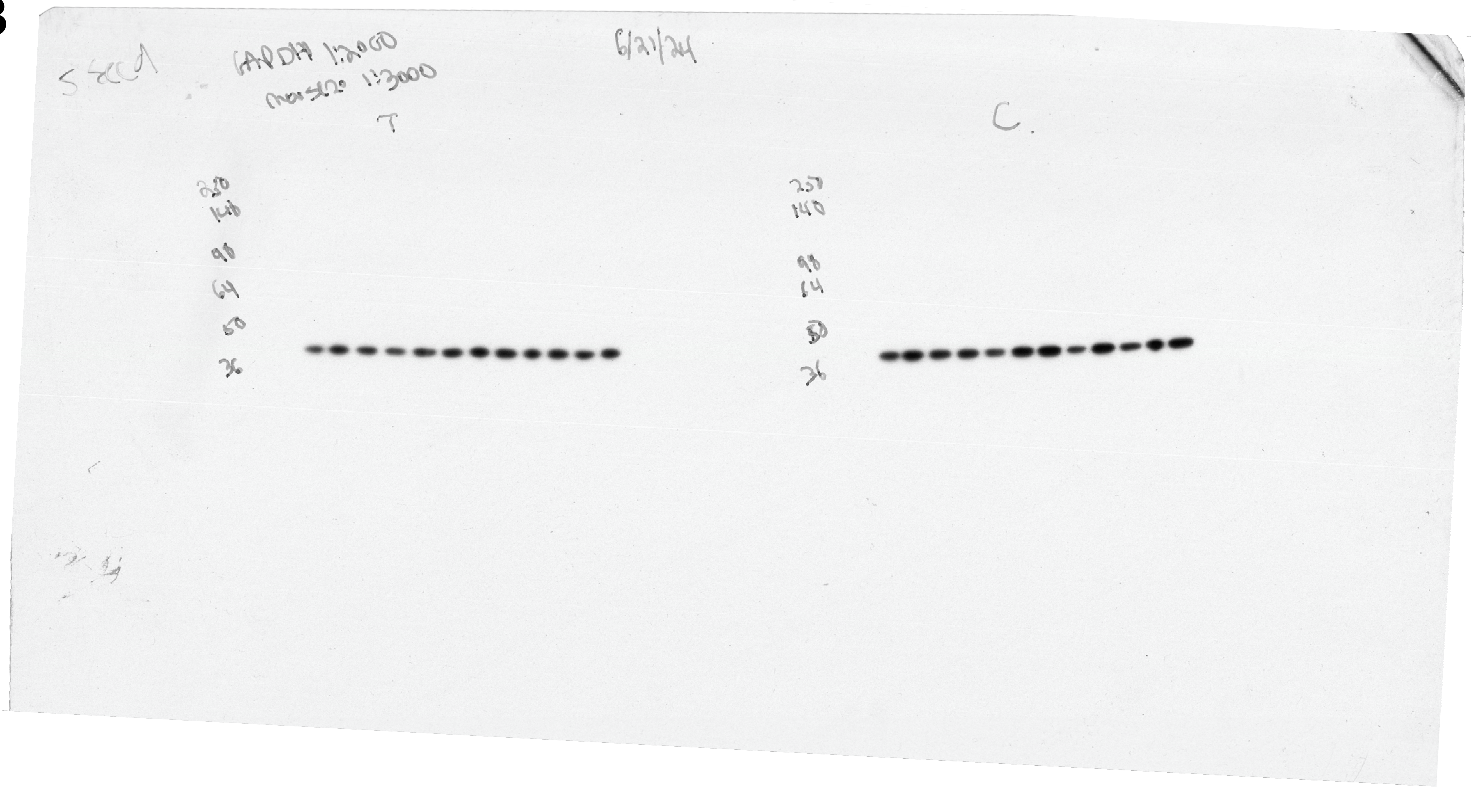

**Supplemental Figure 7: A –** Panel shown in Figure 7 of the manuscript. **B** - Scan of the film used to visualize GAPDH western blots. Molecular weight markers are labelled on the left of the blots. The diagonal line on the corner was used to mark the film's orientation. The panels from figure 7 are flipped horizontally from the original blots to show WTNT, WTTR, TGNT, and TGTR samples from left to right. Hippocampal tissue was analyzed by western blotting. Total (left gel) and cytoplasmic fraction (right gel) from male rats are shown Abbreviations WTNT – wild-type not treated, TGNT – transgenic not treated, WTTR – wild-type BT-11 treated, TGTR – transgenic BT-11treated.

## A

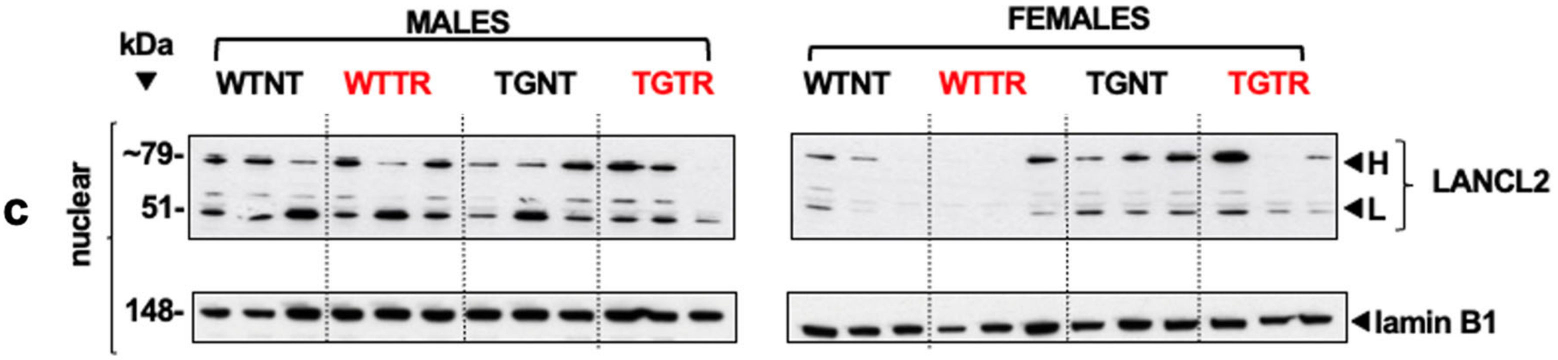

## B

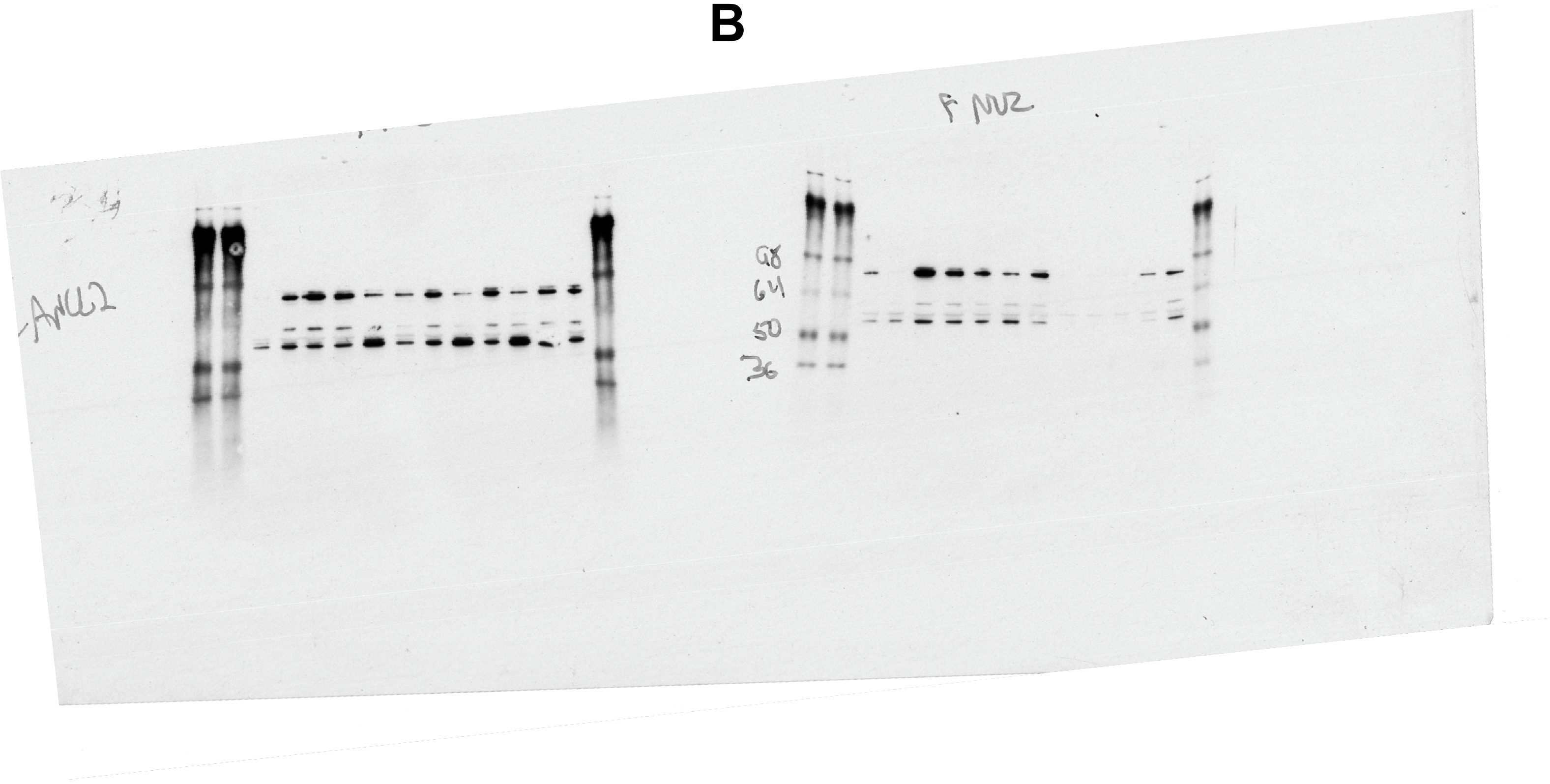

**Supplemental Figure 8: A –** Panel shown in Figure 7 of the manuscript. **B -** Scan of the film used to visualize LANCL2 western blots. The panels from figure 7 are flipped horizontally from the original blots to show WTNT, WTTR, TGNT, and TGTR samples from left to right. Hippocampal tissue was analyzed by western blotting. Male nuclear (left gel) and female nuclear (right gel) fractions are shown. Abbreviations WTNT – wild-type not treated, TGNT – transgenic not treated, WTTR – wild-type BT-11 treated, TGTR – transgenic BT-11treated.

A

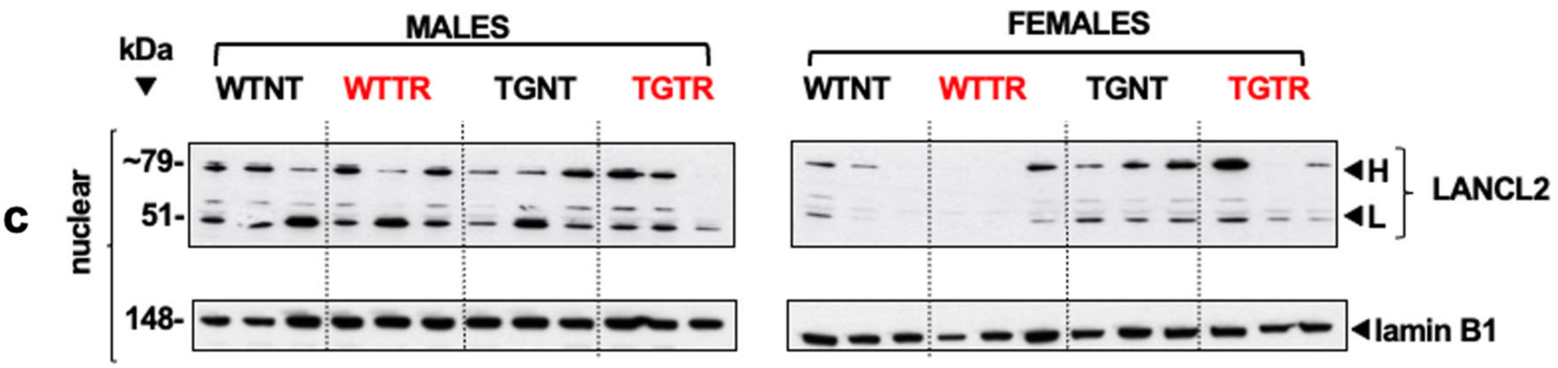

B

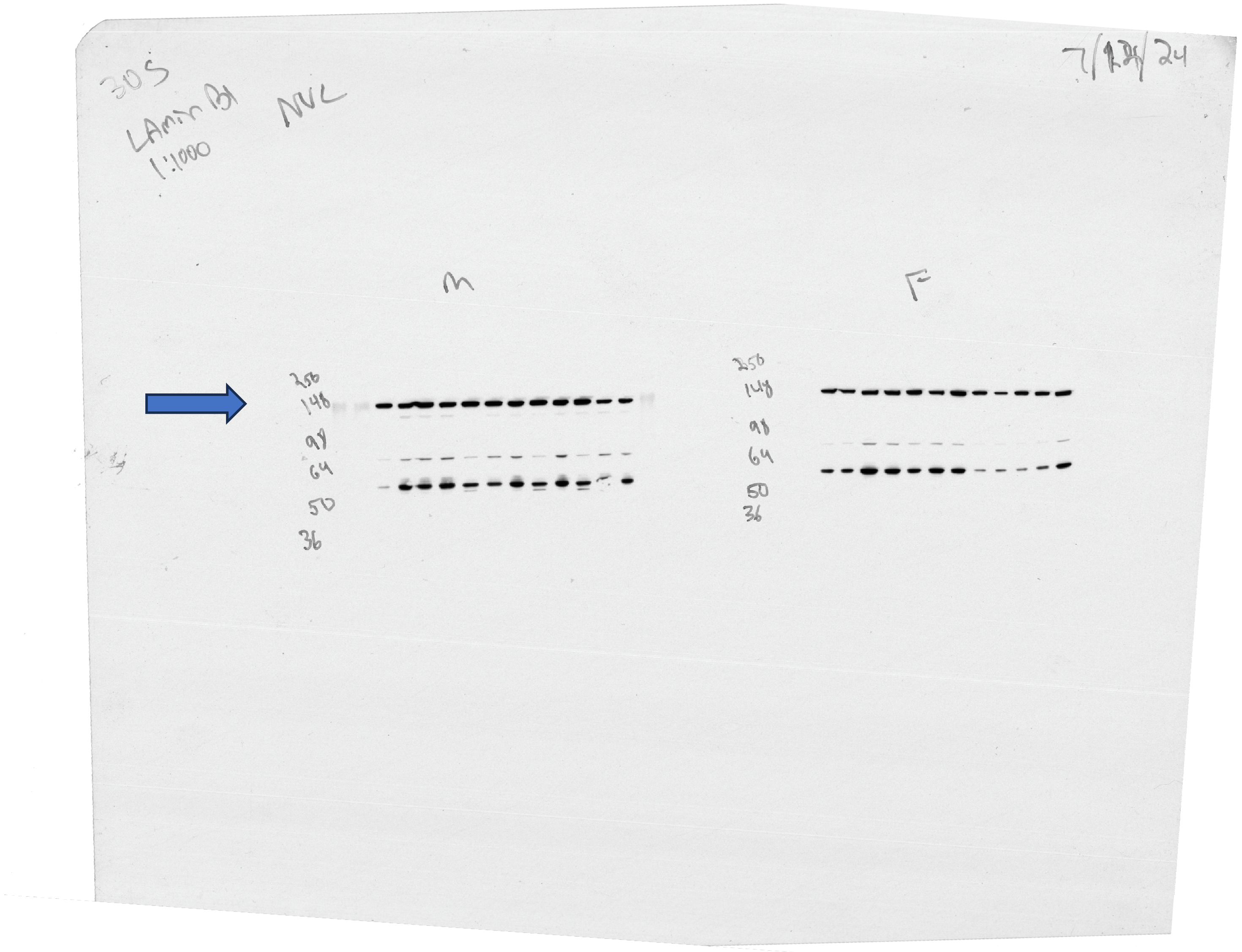

**Supplemental Figure 9: A –** Panel shown in Figure 7 of the manuscript. **B** - Scan of the film used to visualize Lamin B1 western blots (blue arrow). The panels from figure 7 are flipped horizontally from the original blots to show WTNT, WTTR, TGNT, and TGTR samples from left to right. Hippocampal tissue was analyzed by western blotting. Male nuclear (left gel) and female nuclear (right gel) fractions were included. Abbreviations WTNT – wild-type not treated, TGNT – transgenic not treated, WTTR – wild-type BT-11 treated, TGTR – transgenic BT-11treated.

## A

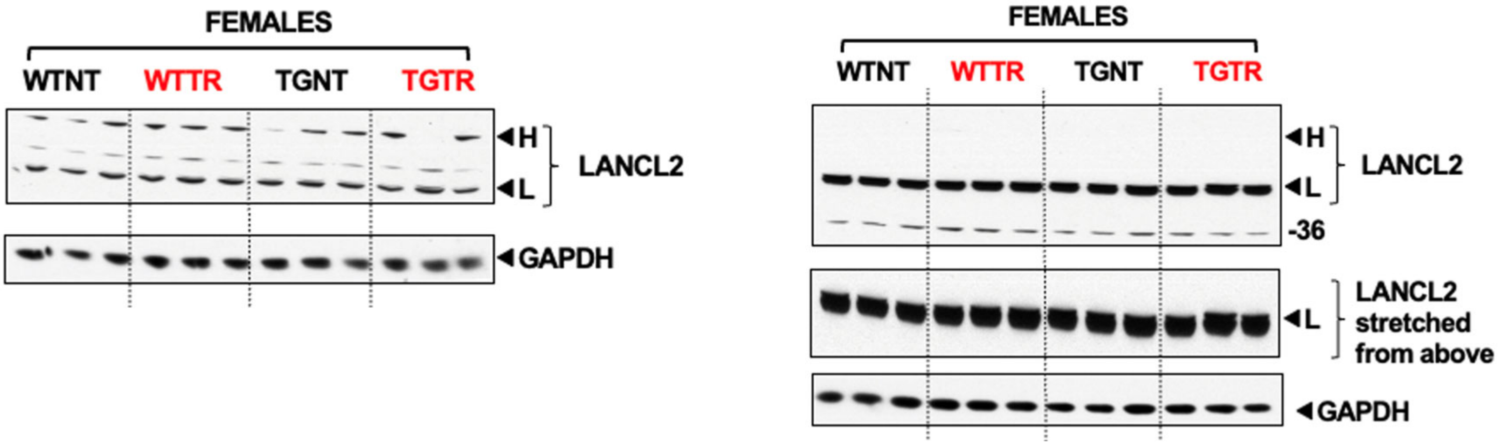

## B

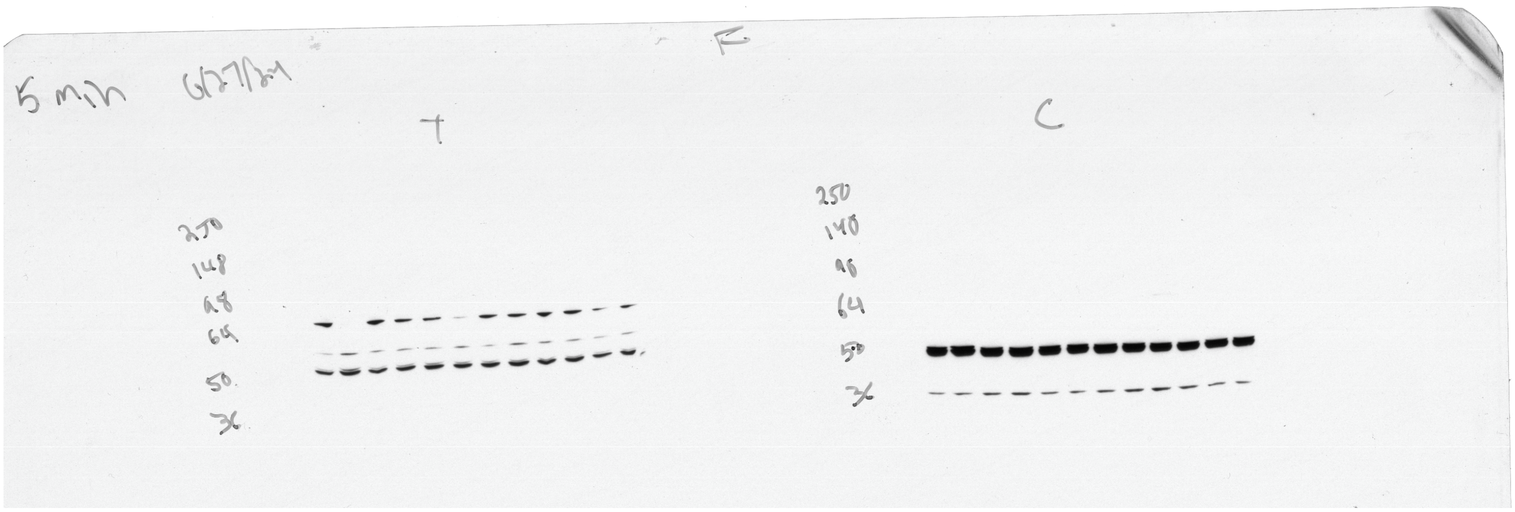

**Supplemental Figure 10: A –** Panel shown in Figure 7 of the manuscript. **B** - Scan of the film used to visualize LANCL2 western blots. The panels from figure 7 are flipped horizontally from the original blots to show WTNT, WTTR, TGNT, and TGTR samples from left to right. Hippocampal tissue was analyzed by western blotting. Female total (left gel) and cytoplasmic (right gel) fractions were included. Abbreviations WTNT – wild-type not treated, TGNT – transgenic not treated, WTTR – wild-type BT-11 treated, TGTR – transgenic BT-11treated.

## A

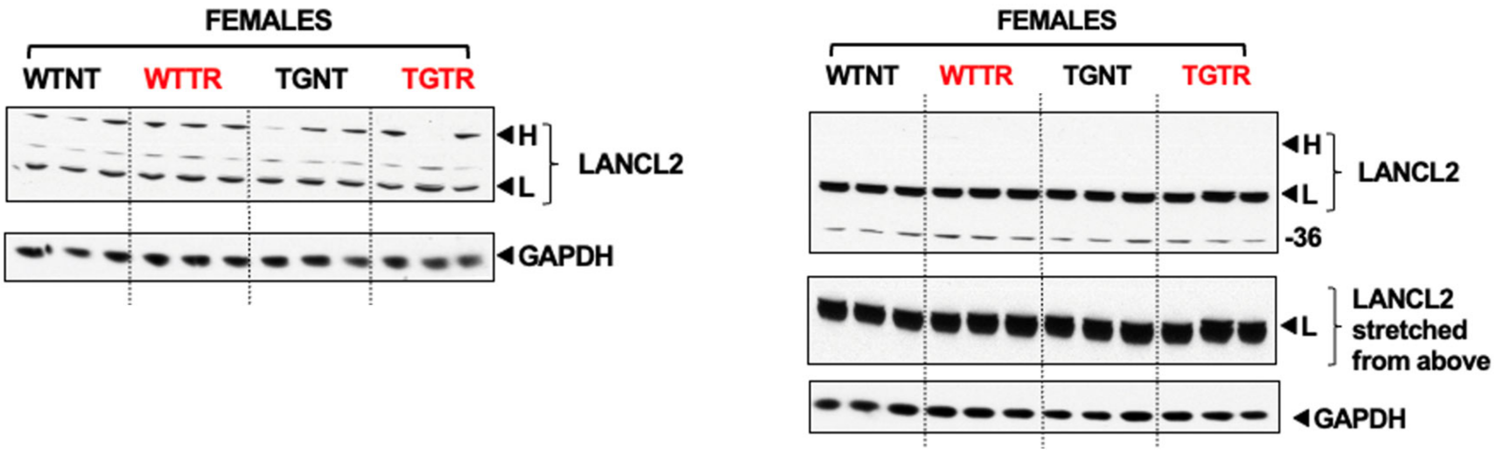

## B

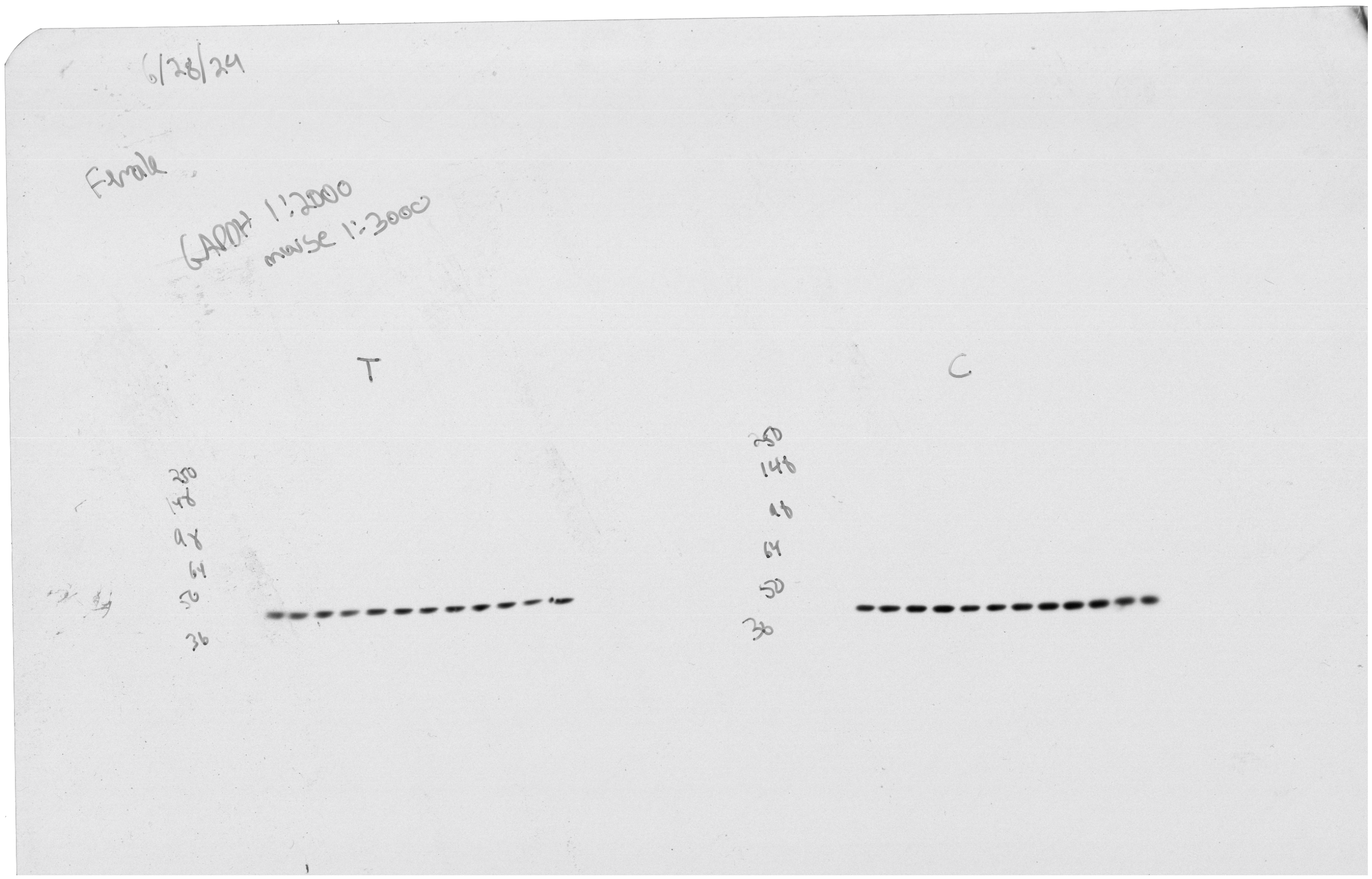

**Supplemental Figure 11: A –** Panel shown in Figure 7 of the manuscript. **B** - Scan of the film used to visualize GAPDH western blots. The panels from figure 7 are flipped horizontally from the original blots to show WTNT, WTTR, TGNT, and TGTR samples from left to right. Hippocampal tissue was analyzed by western blotting. Female total (left gel) and female cytoplasmic (right gel) fractions were included. Abbreviations WTNT – wild-type not treated, TGNT – transgenic not treated, WTTR – wild-type BT-11 treated, TGTR – transgenic BT-11treated.

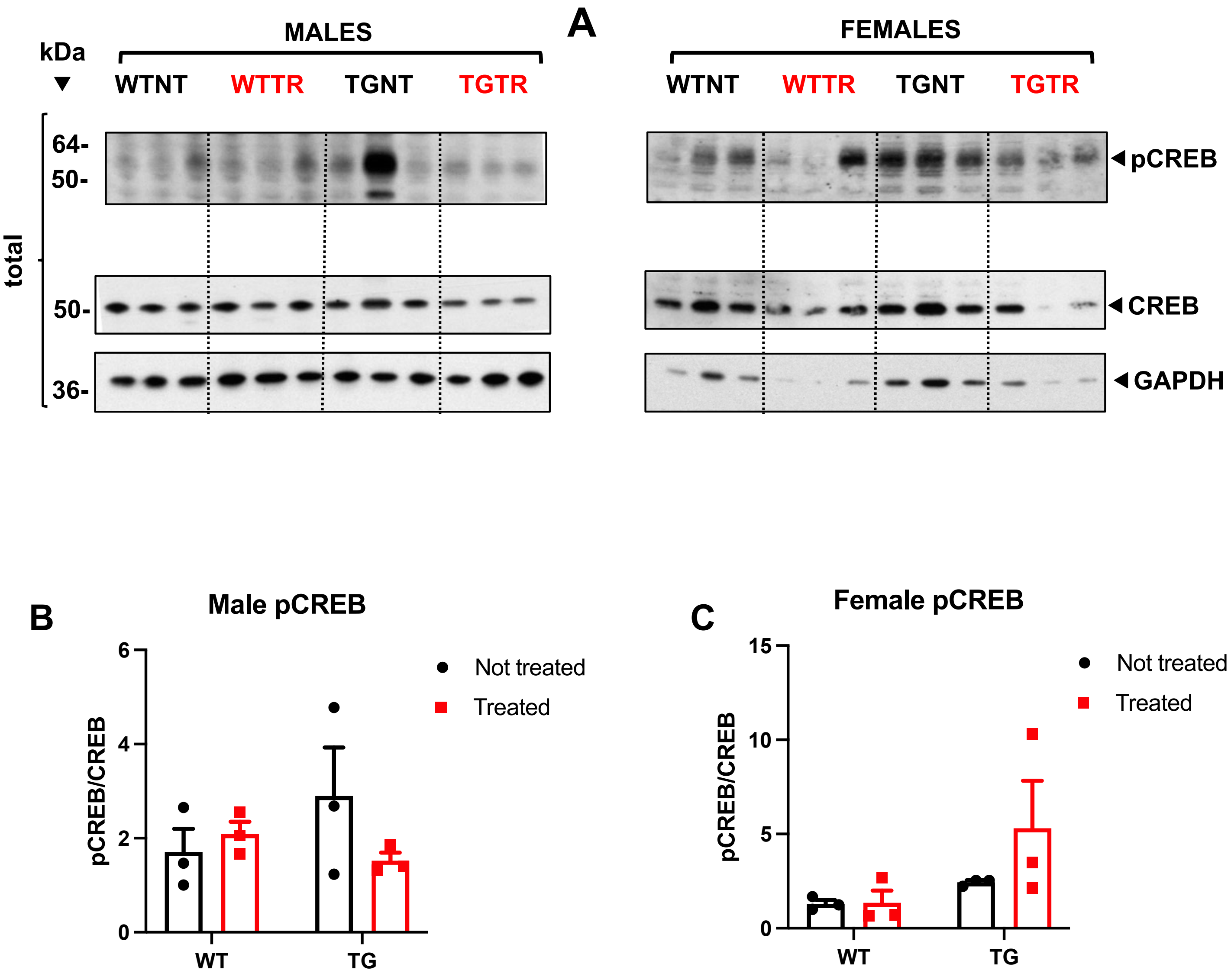

**Supplemental Figure 12:** There are no significant differences in pCREB and total CREB levels across genotype and treatment groups for males and females. Images of western blots across the four groups WTNT, WTTR, TGNT and TGTR for male and female rats. Dorsal hippocampal total, cytoplasmic and nuclear fractions were probed for pCREB and total CREB followed by the molecular marker GAPDH **(A)**. Signal was normalized with GAPDH and was quantified using relative density measurement on ImageJ. Statistical analysis was done on GraphPad Prism using ANOVAs.
WTNT – wild-type not treated, TGNT – transgenic not treated, WTTR – wild-type BT-11 treated, TGTR – transgenic BT-11treated.

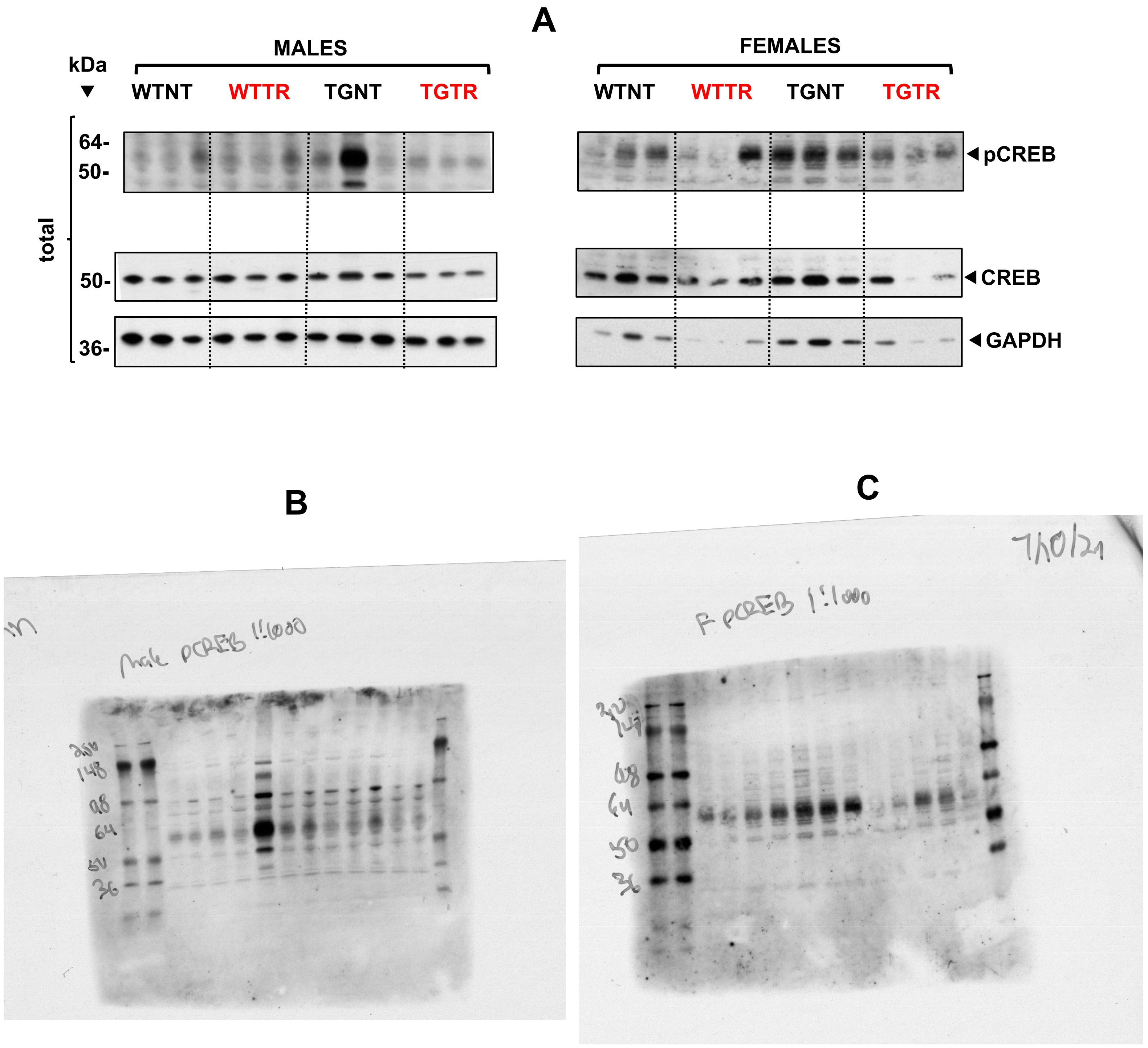

**Supplemental Figure 13: A –** Panel shown in Supplemental Figure 12. **B and C -** Scans of the films used to visualize pCREB western blots. The panels from supplemental figure 12 are flipped horizontally from the original blots to show WTNT, WTTR, TGNT, and TGTR samples from left to right. Hippocampal tissue was analyzed by western blotting. Male (left gel) and female (right gel) rats were included. Abbreviations WTNT – wild-type not treated, TGNT – transgenic not treated, WTTR – wild-type BT-11 treated, TGTR – transgenic BT-11treated.

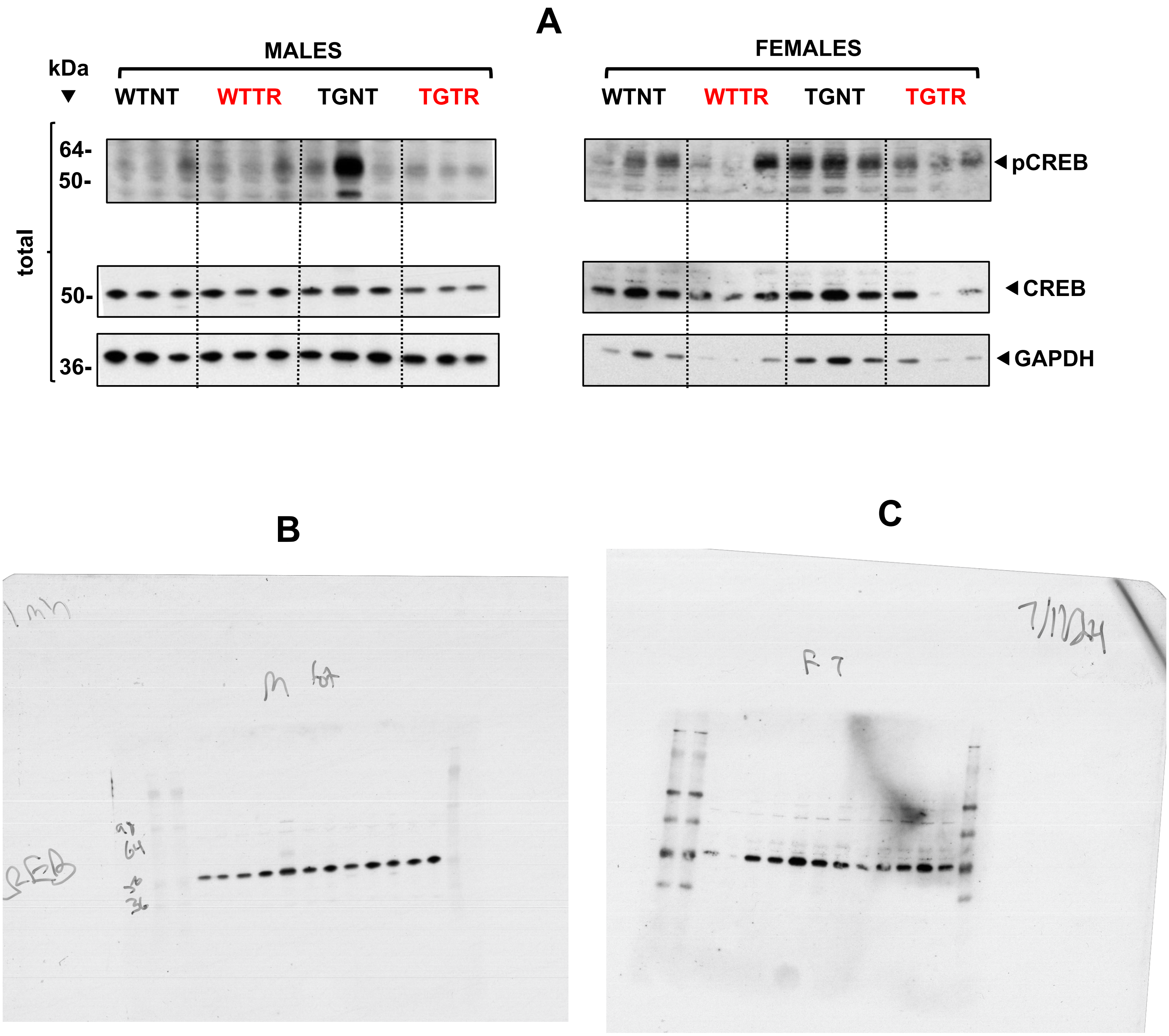

**Supplemental Figure 14: A –** Panel shown in Supplemental Figure 12. **B and C -** Scans of the films used to visualize CREB western blots. The panels from supplemental figure 12 are flipped horizontally from the original blots to show WTNT, WTTR, TGNT, and TGTR samples from left to right. Hippocampal tissue was analyzed by western blotting. Male (left gel) and female (right gel) rats were included. Abbreviations WTNT – wild-type not treated, TGNT – transgenic not treated, WTTR – wild-type BT-11 treated, TGTR – transgenic BT-11treated.

**A**

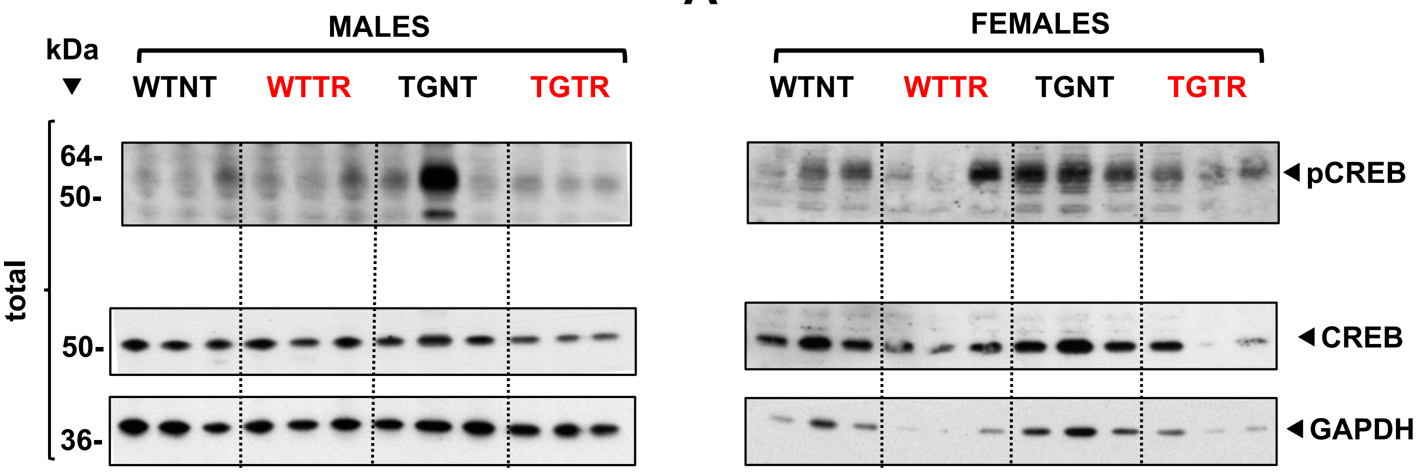

**B**

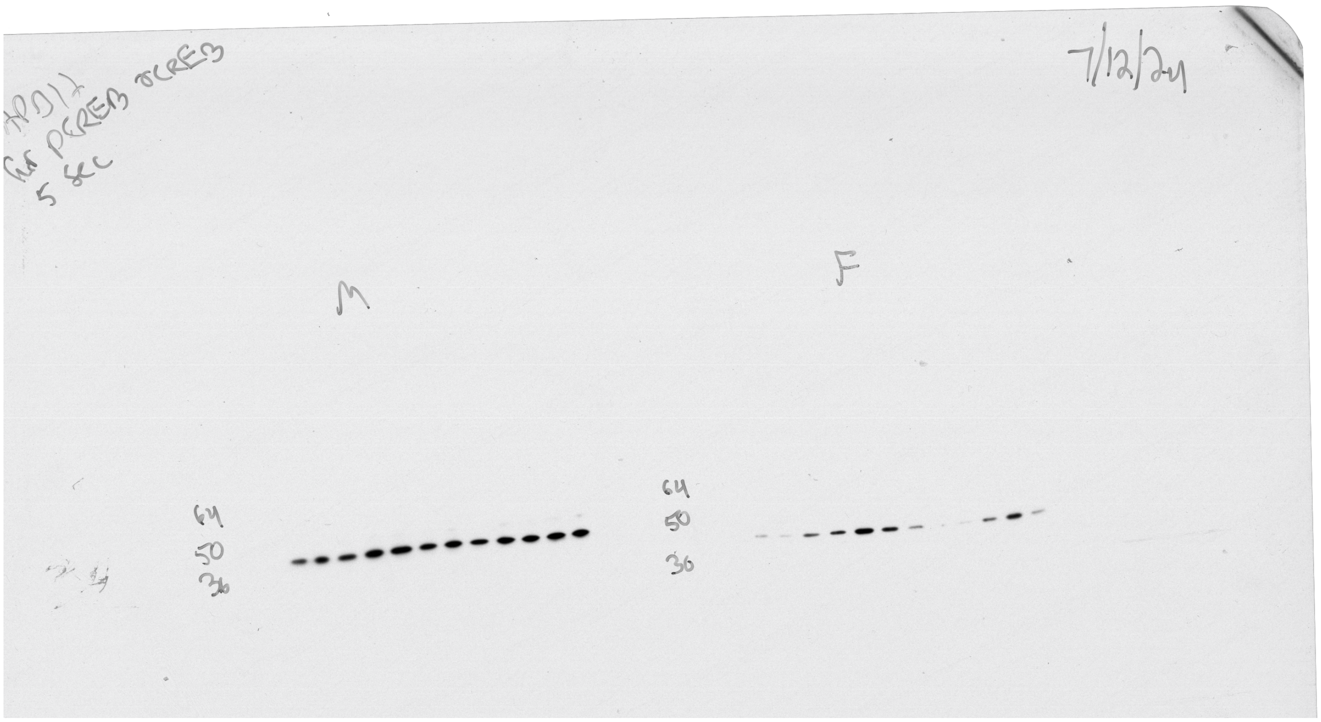

**Supplemental Figure 15: A –** Panel shown in Supplemental Figure 12. **B** - Scan of the film used to visualize GAPDH western blots. The panels from supplemental figure 12 are flipped horizontally from the original blots to show WTNT, WTTR, TGNT, and TGTR samples from left to right. Hippocampal tissue was analyzed by western blotting. Male (left gel) and female (right gel) rats were included. Abbreviations WTNT – wild-type not treated, TGNT – transgenic not treated, WTTR – wild-type BT-11 treated, TGTR – transgenic BT-11treated.

**Supplemental Figure 16**

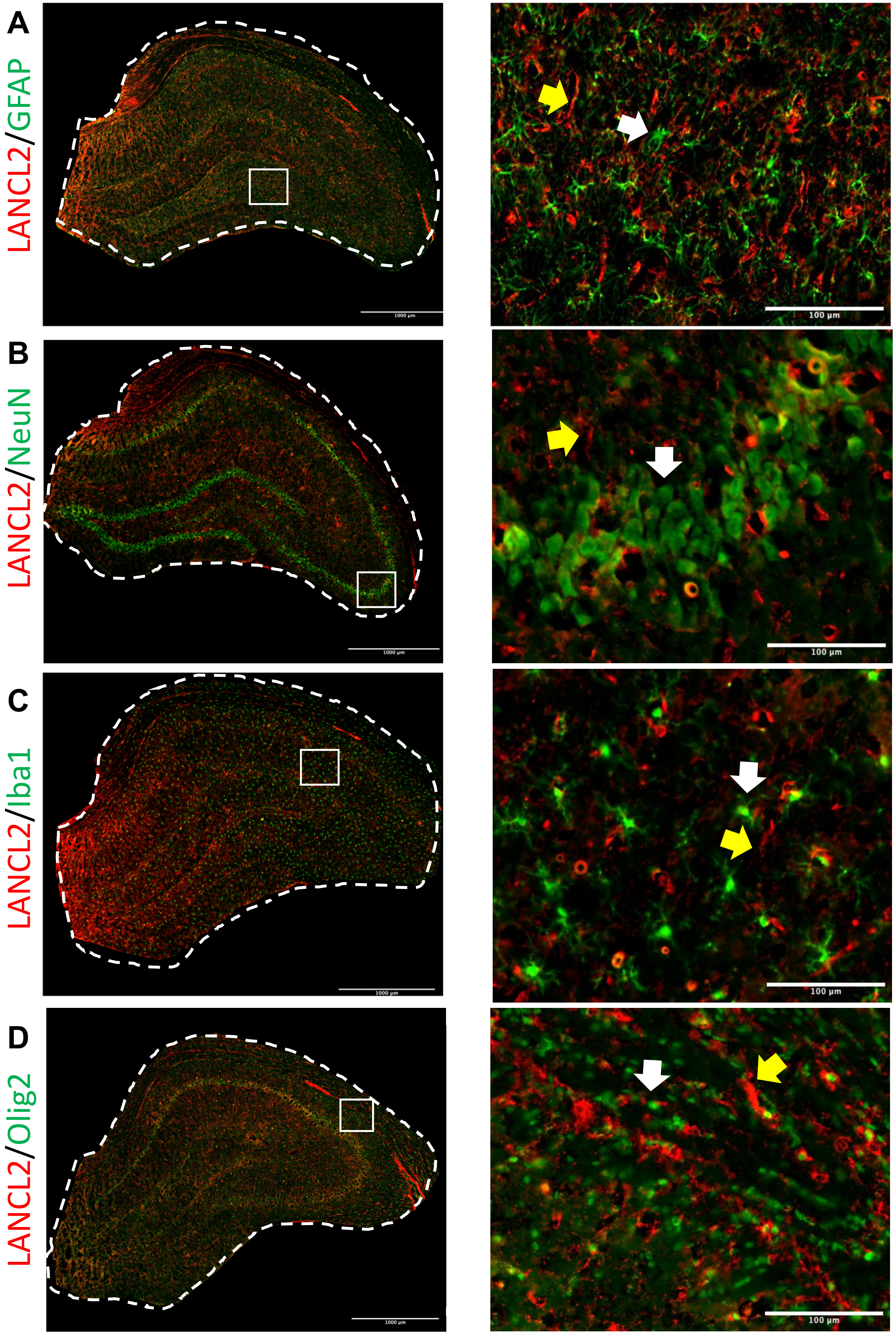

### Supplemental Figure 17

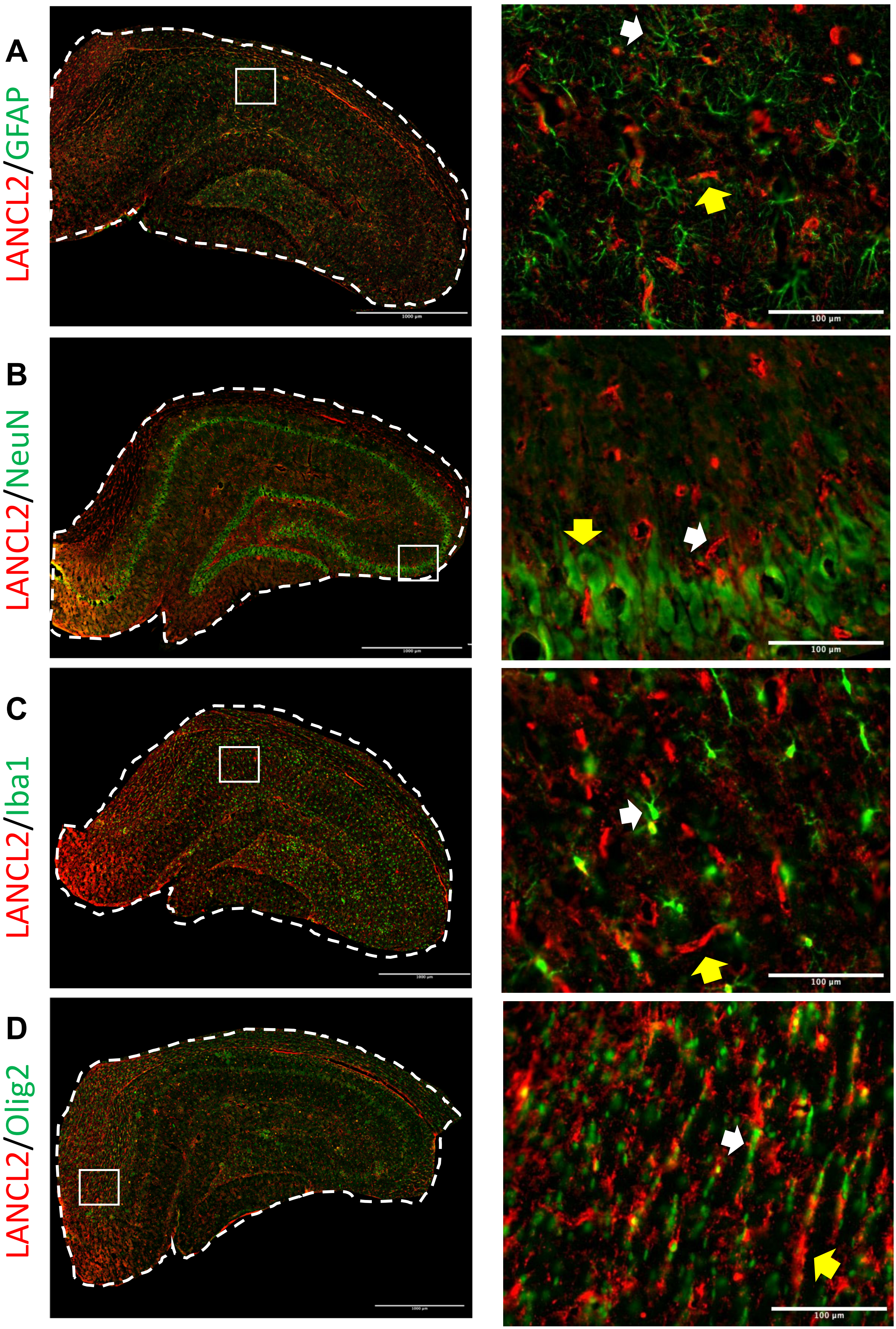

### LANCL2 is localized in oligodendrocytes in the dorsal hippocampus of WT male rats

Supplemental Figure 16: LANCL2 is detected in oligodendrocytes located in the dorsal hippocampus and neighboring corpus callosum of WTNT male rats. Representative images of LANCL2 (red) with GFAP for astrocytes (A), with NeuN for neurons (B), with Iba1 for microglia (C), and with Olig2 for oligodendrocytes (D). All cell type specific markers are shown in green. Left panels show a sagittal section of the dorsal hippocampus. Right panels show close ups in relevant areas to visualize individual cells. Higher magnification images are shown in the manuscript Figure 8 a-h. White arrows point to the cell type shown in green, yellow arrows point to LANCL2 staining shown in red.

### LANCL2 is localized in oligodendrocytes in the dorsal hippocampus of TG male rats

Supplemental Figure 17: LANCL2 is detected in oligodendrocytes located in the dorsal hippocampus and neighboring corpus callosum of TGNT male rats. Representative images of LANCL2 (red) with GFAP for astrocytes (A), with NeuN for neurons (B), with Iba1 for microglia (C), and with Olig2 for oligodendrocytes (D). All cell type specific markers are shown in green. Left panels show a sagittal section of the dorsal hippocampus. Right panels show close ups in relevant areas to visualize individual cells. Higher magnification images are shown in the manuscript Figure 8 i-p. White arrows point to the cell type shown in green, yellow arrows point to LANCL2 staining shown in red.
